# Autism in a dish: ES cell models of autism with copy number variations reveal cell-type-specific vulnerability

**DOI:** 10.1101/2022.02.02.478766

**Authors:** Jun Nomura, Amila Zuko, Keiko Kishimoto, Hiroaki Mutsumine, Kazumi Fukatsu, Yoshiko Nomura, Xiaoxi Liu, Nobuhiro Nakai, ES library team, Eiki Takahashi, Tsukasa Kouno, Jay W. Shin, Toru Takumi

## Abstract

Human genetics has identified numerous single nucleotide variations (SNVs) and copy number variations (CNVs) associated with autism spectrum disorders (ASD) and other psychiatric disorders. However, the lack of standardized biological resources impedes understanding of the common pathophysiology of ASD. Here, using next-generation chromosome engineering based on the CRISPR/Cas9 system, we established a biological resource including 65 genetically modified mouse embryonic stem cell (mESC) lines as genetic models of human SNVs and CNVs. To illustrate cell-type and CNV specific molecular features of ASD, we performed single-cell RNA sequencing (37,397 cells in total), morphological, and physiological analyses using 12 representative cell lines with CNVs highly associated with ASD. These results uncover gene ontology (GO) terms, canonical pathways, upstream regulators, and related neuropsychiatric disorders in a cell-type and CNV specific manner.

## INTRODUCTION

Autism spectrum disorder (ASD) is a highly heritable neurodevelopmental disorder characterized by social deficits with restricted interests and repetitive behaviors. Although considerable heterogeneity of genetics and clinical phenotypes have been reported in ASD^1^, remarkable advances in sequencing technologies have identified numerous *de novo* single nucleotide variants (SNVs) and copy number variations (CNVs) associated with ASD^2–4^. CNVs generally include multiple genes with regulatory elements such as promoter, enhancer, and repressor in the genome, which may contribute to the complexity and comorbidity of ASD pathology. Thus, it is ideal to analyze multiple CNVs in the same experimental platform to identify convergent pathways and molecular networks implicated in the pathophysiology of ASD. To date, these human genetic data are archived in web-based databases such as the Simons Foundation Autism Research Initiative (SFARI) and AutDB to adopt increasing genetic variants found in patients with ASD^5,6^. These human genetic data have also identified several CNVs derived from patients with ASD overlapping with those observed in other neuropsychiatric diseases such as schizophrenia and bipolar disorders^7,8^.

Genetic evidence-based biological research is still challenging because of the lack of standardized bioresources. To overcome these limitations, we developed an ASD-associated CNV cell bank as a biological resource for ASD using mouse embryonic stem cells (mESCs) and a next-generation chromosome engineering technique based on CRISPR/Cas9 system. This unique bioresource includes 65 CNVs with deletions and duplications covering 58 human chromosome loci and 175 additional vectors targeting these loci. These mESC cell lines provide major benefits as a biological resource for generating mutant mice, transplantation to living animals, and blastocyst complementation to assess neural development and morphogenesis of the *in vivo* brain environment^9^. Using neural cells derived from 12 representative cell lines we tried to identify the features commonly dysregulated in ASD by applying morphological, physiological, and single-cell transcriptome analyses.

## RESULTS

### Annotation of human CNVs to mouse genomic loci

To develop the ASD-associated CNV cell bank as a comprehensive biological platform for ASD research (Fig. 1a), we first referred to the SFARI database (https://gene.sfari.org/database/human-gene/) together with published data to survey ASD-associated CNVs. The SFARI database is a database for ASD by integrating published human genetic and animal model data. It contains more than 1,000 genes and 2,000 CNVs associated with ASD^6^. Using this database, we first listed 104 ASD-associated CNVs as candidates for the followed chromosome targeting (Extended Data Table 1). We analyzed syntenic regions between humans and mice for each CNV. Some CNVs, 35 out of 104 loci, such as 1q44, 14q11.2, and Xp22.31, were omitted from our chromosome targeting list because of the low conservation in the mouse genomic structure (Supplementary Table 1). Other CNVs which are recognized as significant risk loci across multiple psychiatric disorders have remained in our list, such as 1q21.1, 2p16.3 (*NRXN1*), 3q29, 7q11.23, 8p23.1, 15q11.2, 15q13.3, 16p11.2 (proximal region, from breakpoint (BP)4 to BP5), 16p13.2, 22q11.21, and 15q11-q13 Prader-Willi syndrome region (Extended Data Table 2)^7,8,10^. These CNVs contain genes associated with other psychiatric disorders such as schizophrenia, bipolar disorders, attention-deficit hyperactivity disorder (ADHD), and intellectual disability (ID). Other major CNVs, such as 3p26.3, 15q11.2-q13.1, 15q13.3, and 22q11.21, also include multiple psychiatric risk genes (Extended Data Table 3).

**Figure 1.**
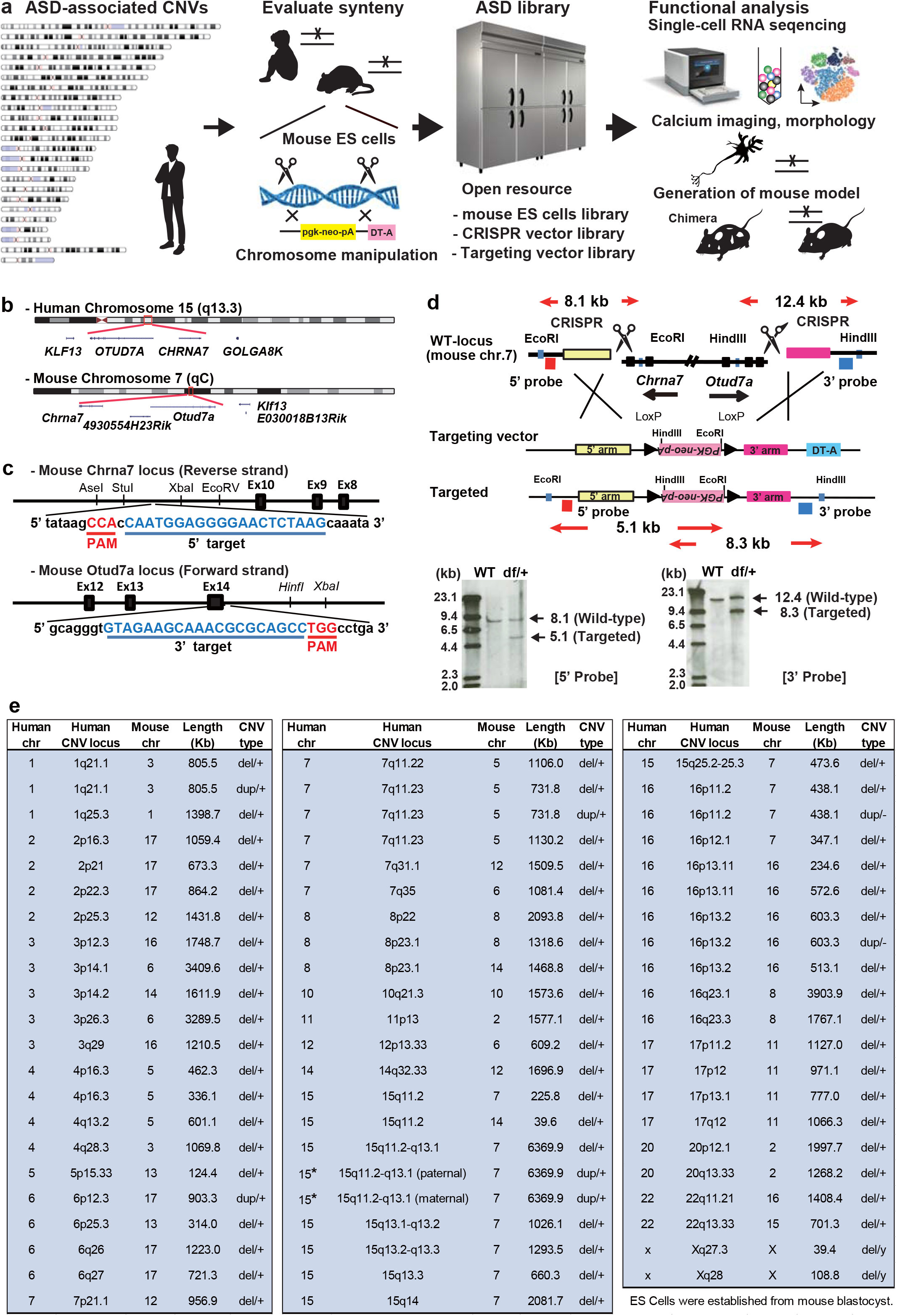
ASD cell bank for functional analysis. **a**, Schematic diagram to generate a cellular model of ASD with chromosomal abnormalities, and strategy to identify ASD-associated pathogenic phenotypes using representative ASD cell lines. **b**, Representative image of chromosome targeting locus in mouse chromosome 7 (7qC) corresponds to human 15q13.3 locus. **c**, Schematic of the sgRNA targeting sites in *Chrna7* and *Otud7a*, respectively. The target sequences are underlined and labeled in blue. The protospacer adjacent motif (PAM) sequence is underlined and labeled in red. **d**, Structures of the mouse 7qC genomic locus, the targeting vector, and the targeted locus mediated by genome editing technique. The restriction enzymes used for Southern blotting analysis are shown. The targeting vector contains the PGK-neo-pA cassette with diphtheria toxin A-fragment (DT-A) for positive and negative selection. The 5’ and 3’ probes are shown as red and blue boxes, respectively. EcoRI-digested genomic DNA was hybridized with a 5’ probe to produce an 8.1 kb band from wild-type (WT) and a 5.1 kb band from the targeted locus. HindIII-digested genomic DNA was hybridized with a 3’ probe to produce a 12.4 kb band from WT and an 8.3 kb from the targeted locus. e. Targeted mouse chromosome loci corresponding to human ASD-associated chromosome loci.

### Generation of cell models for ASD by using next-generation chromosome engineering

To generate cell models of ASD, we developed next-generation chromosome engineering using the CRISPR-based genome editing technique. First, we introduced two CRISPR/Cas9 vectors, pX330^11^ with a 20-nucleotide (nt) guide sequence and a targeting vector with short homology arms (1∼2 Kb) into murine C57BL/6J background ES cells (mESCs), CMTI-2. We introduced the CRISPR/Cas9 pX330 vectors without a 20-nt target sequence into mESCs as a control cell line. This next-generation chromosome engineering using a targeting vector together with CRISPR/Cas9 vectors based on homology-directed repair (HDR) brought ∼10% targeting efficiency with low false positives cells by a diphtheria toxin A (DT-A) fragment as a negative selection marker (Fig. 1b-d).

Using *in silico* and experimentally validated 120 CRISPR vectors and 55 targeting vectors (Supplementary Table 2)^12^, we obtained 65 cell lines, including 58 deletions (one-copy, or null mutation (knockout)), 2 tandem duplication (two-copies), and 5 duplications (three-copies) (Fig. 1e, Extended Data Tables. 4, 5). Two out of 65 cell lines, mouse chromosome 7 corresponding to human 15q11.2-q13.1 duplication (paternal or maternal duplication, respectively), were established from mouse blastocyst by crossing paternally inherited 15q11.2-q13.1 duplication male mice^13^ with C57BL/6J wild-type (WT) female mice or C57BL/6J WT male mice with maternally inherited 15q11.2-q13.1 female mice^13^, respectively. All cell lines were verified for targeted deletion or duplication by PCR, and some lines were followed by array comparative genomic hybridization (aCGH) and Southern blotting analysis.

One of the advantages of mESC is its application to generating mouse models. We developed a mouse model with CNV using a cell line corresponding to human chromosome 15q13.3. This human locus encompasses two genes, *CHRNA7* and *OTUD7A*, which are highly conserved in the mouse 7qC locus (Fig. 1b). The targeted 15q13.3 heterozygote mESCs were injected into blastocysts, and chimera offspring mice were generated (Extended Data Figs. 1a, b). The sperms of 60% chimera mice were then fertilized with C57BL/6J WT egg *in vitro* to produce mice lacking one copy of the 15q13.3 allele (15q13.3(+/-)). Genotype was confirmed by PCR (Extended Data Fig. 1c). Using adult cortices, the gene expression in 15q13.3 was assessed by quantitative real-time RT-PCR (RT-qPCR). The result showed approximately 50% reduction of *Chrna7* and *Otud7a* in 15q13.3(+/-) mice, respectively (Extended Data Fig. 1d).

Using 15q13.3 (+/-) mice, we performed a battery of behavioral tests and compared the results with previous studies^14,15^. Although 15q13.3(+/-) mice were healthy and fertile with no gross physical abnormalities, they showed social deficits in the three-chamber social interaction test, increased startle response to a sudden noise, and increased body weights in the developmental period, which was observed in human subjects with 15q13.3 deletion^14^. These results illuminate *Chrna7-Otud7a* genetic interaction relevant to ASD-like symptoms (Extended Data Fig. 1e-p). This test successfully narrowed down the critical region for social deficits in the 15q13.3 microdeletion syndrome from 1.5 Mb (from *Chrna7* to *Fan1*) to 0.7 Mb (from *Chrna7* to *Otud7a*) and demonstrate the significance of mice generated using mESCs for modelling human psychiatric disorders *in vivo*.

### Morphological and physiological analyses of cell models with ASD-associated CNV

To analyze biological aspects of ASD-associated CNVs, we selected 12 CNVs as representatives for the following experiments, duplication of 1q21.1 (MIM: 612475), deletion of 2p16.3 (MIM: 614332), 3q29 (MIM: 609425), duplication of 7q11.23 (MIM: 613729), deletion of 15q11.2 (MIM: 615656), 15q13.3 (MIM: 612001), 16p11.2 (MIM: 611913), 16p13.2 (MIM: 616863), 17p11.2 (MIM: 182290), 17q12 (MIM: 614527), Xq27.3 (MIM: 300624) and, Xq28 (MIM: 312750) (Extended Data Table 1). These CNVs were selected based on two previous genetic ASD cohorts studies, *de novo* CNVs from the Simons Simplex Collection (SSC) 2,591 families^16^ and both *de novo* and rare CNVs on ASD risk in multiplex 1,532 families from the Autism Genetic Resource Exchange (AGRE)^17^. Xq27.3 (*Fmr1*) and Xq28 (*Mecp2*) were selected as a monogenic cause of syndromic ASD^18^. Genes located in each CNV were highly conserved in mice (Extended Data Table 6). Targeted deletion or duplication in these representative CNVs was confirmed by a-CGH (Extended Data Fig. 2) and gene expression profiles in the targeted loci (Extended Data Table 7).

We next differentiated these ES cells into neurons. On day 1 of *in vitro* differentiation, differentiating neurons were transfected with a green fluorescent protein (GFP) expression vector and fixed on day 3 to visualize them. Although we measured axon length, total neurite length (μm), and the number of neuronal branches, we found no significant difference between these mutants and control (Extended Data Fig. 3a-c). We then assessed neuronal response based on activity-dependent calcium influx. Differentiated neurons were loaded with the calcium indicator, Fluo-4, and their intracellular calcium mobilization was measured by application of 25 mM KCl, leading to membrane depolarization (Extended Data Fig. 3d). One-way ANOVA showed a significant effect of genotype for response amplitude of the fluorescent intensity, (ΔF/F), F(12,185)=5.27, (p < 0.001). The Bonferroni multiple comparisons test for *post-hoc* comparisons revealed a significance in 3q29 deletion vs. control cells (p < 0.001) (Extended data Fig. 3e). The result is consistent with the previous report using 3q29 deletion model mice, which showed excitatory/inhibitory imbalance derived from increased excitatory neural activity in the cerebral cortex^19^. These results indicate that the ES cell models could be morphologically and physiologically differentiated.

### Cell-type-specificity of gene expression and genetic associations with ASD

To reveal cell-type-specific expression of our multiple ASD cell lines, we performed single-cell RNA-sequencing (scRNA-seq) on the 10x Genomics platform using differentiated neuronal cells of 12 representative ASD-associated CNVs and a control mESCs line. We collected 37,397 cells in total, 2,858 cells from control and 34,539 cells from cell models across 12 CNVs. We then visualized all single-cells in a uniform manifold approximation and projection (UMAP) space to analyze cell-type-specific features in each cell model with CNV. The cell-type was annotated according to gene expression of canonical cell-type-specific markers (e.g., *Slc17a6* for glutamatergic neurons, *Gad1* for GABAergic neurons, *Fabp7* for neural stem cells and immature astrocytes, *Neurod6* for neural progenitors, *Ifrd1* for microglia, *Pdgfra* for OPC (oligodendrocyte precursor cell), *Col1a1* for endothelial cells, and *Pou5f1* for ES cells) and we finally identified 17 cell-types in total (Fig. 2a-d, Supplementary Table 3)^20–23^. The cell-type-specific differentially expressed genes (DEGs) include cortical upper-layer (2/3) specific glutamatergic neuronal genes such as *Cux2, Hap1*, and *Tmem145*^24,25^. In addition, UMAP revealed two different types of GABAergic neuronal clusters, one with *Gad1*+*Sst*+*Npy*+ and the other one with *Gad1*+*Sst*+*Npy*-. Both of these clusters expressed GABAergic marker *Gad1* with a subtype marker *Sst*, which has been classified as Martinotti cells^26^. Although GABAergic *Npy+* cells are widely expressed throughout the cortex, Martinotti-like cells co-expressed with *Sst* (*Sst+Npy+*) are regarded as the most excitable type^27^.

**Figure 2.**
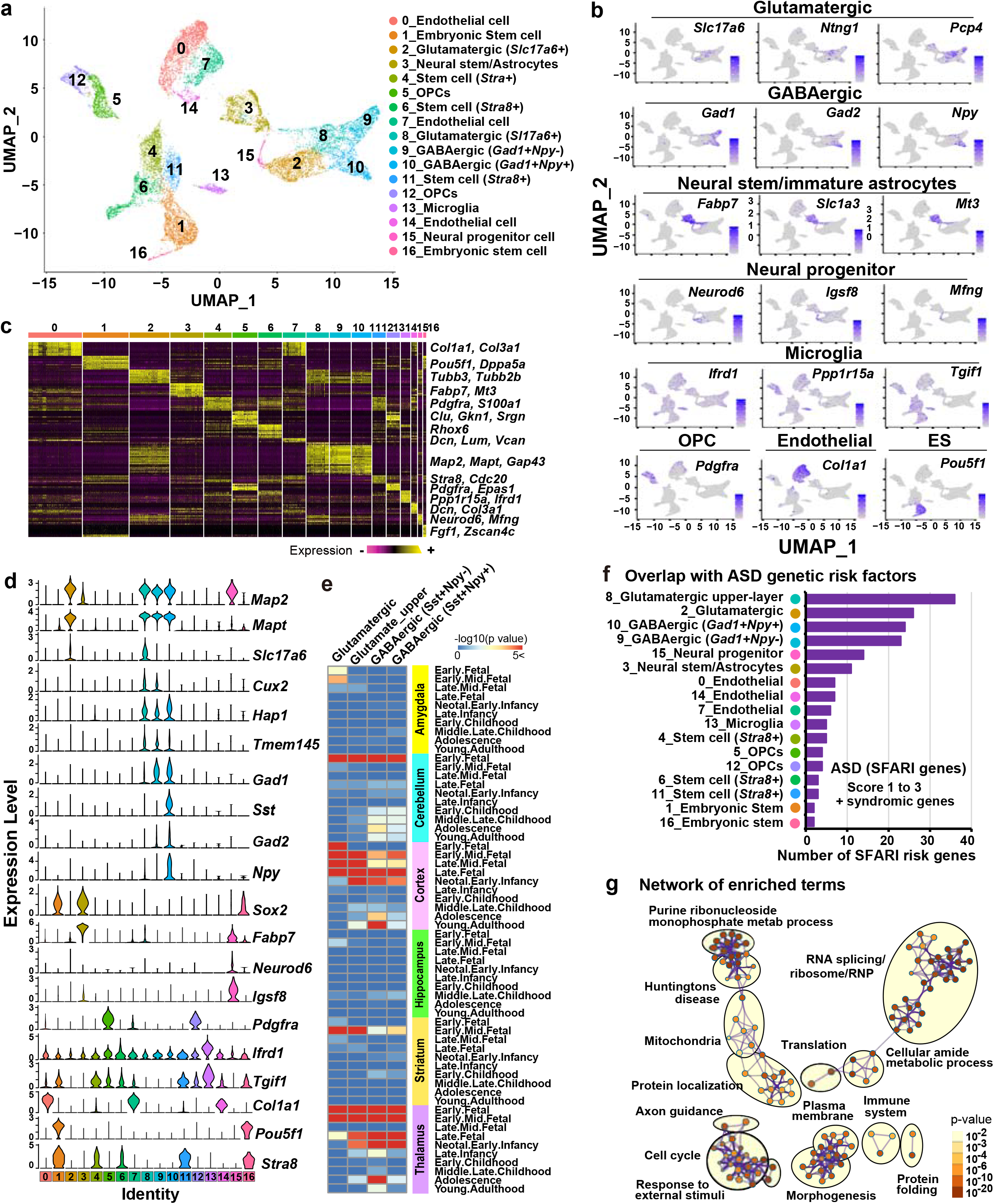
Transcriptional features and heterogeneity of cells harboring ASD-associated CNVs. **a**. UMAP dimensionality reduction embedding of differentiated neurons and the derivatives from mouse ES cells harboring ASD-associated CNVs. Whole cells were colored by annotated cell types. **b**. Feature plots of characteristic marker genes for the cell types are shown. Cells are color-coded according to gene expression levels. **c**. Heatmaps show the expression pattern of cell-type-specific genes. Columns represent individual cells, and rows represent individual genes. The representative differentially expressed genes are listed to the right. **d**. Violin plot showing the expression profile of each gene in various cell types. **e**. Cell-type-specific Expression Analysis (CSEA) using each glutamatergic or GABAergic cell-cluster-specific genes (top 300 independent genes, q < 0.001). It highlights the brain area with the development period in each neuronal cluster. **f**. The number of overlapping genes between genetic risk factors (SFARI genes, risk score 1, 2, 3, and syndromic genes) and cell-type-specific DEGs (top 300 genes, q < 0.001) in each cell cluster. Color-coding is consistent with **(a and c). g**. Network visualization of enriched terms. The network is visualized using Metascape and Cytoscape. Each node represents an enriched GO/pathway term and is colored by their p-values. Edges link similar terms.

The cell type with the annotation was also confirmed by Gene Ontology (GO), the Kyoto Encyclopedia of Genes and Genomes (KEGG), and Reactome enrichment analysis (Extended Data Table 8). Brain regions and developmental stages were determined by cell-type-specific expression enrichment analysis (CSEA) using 300 cluster-specific DEGs (adjusted p < 0.001)^28^. These neuronal gene sets were mainly enriched in both cortex and thalamic regions from the early fetal to late infancy period (Fig. 2e).

We also analyzed the interaction between ASD genetic risk factors (SFARI genes, risk score 1 to 3 plus syndromic genes) and cell-type-specific DEGs (top 300, adjusted p < 0.001) to assess the cell-type-specific contribution to ASD pathophysiology and pathogenesis (Fig. 2f). Remarkably, the upper-layer specific glutamatergic neuronal cluster largely overlapped with SFARI ASD risk genes (Fig. 2f, top rank). The tendency was consistent with a single-cell transcriptome study using postmortem cortical tissues from autistic individuals^29^. ASD risk genes were also highly expressed in neural stem and progenitor cells (Fig. 2f, 5th and 6th ranks), suggesting the pathological primed stage in neural fate^30^. In contrast, the effects of microglia (1.7%), OPCs (1.3%), and endothelial cells (2.3-1.7%) on ASD pathology were likely weaker than those of neuronal cells. However, biological evidence supports the importance of glial cells, and even in endothelial cells in the brain microenvironment for ASD pathology^31,32^. Indeed, vascular contributions of the 16p11.2 CNV have been recently published^33^.

### Molecular signatures of cell models with ASD-associated CNVs

Several convergent pathways underlying ASD pathophysiology have been suggested from studies using lymphoblasts, postmortem brains, and iPSCs from patients with ASD^34–36^. We investigated convergent features of ASD as well as each CNV from GO and pathway analysis using 12 representative ASD cell models. First, CNV specific DEGs were analyzed by GO and GO-network analysis. As expected, representative ASD-associated CNVs were significantly enriched in neuron, transcription, and translation-associated terms regardless of genotype, consistent with previous studies^2^ (Fig. 2g and Extended Data Table 9). In this enrichment analysis, immune systems were enriched in the ASD network. They were also repeatedly reported as a risk factor for ASD and other psychiatric disorders^37^. Interestingly, enrichment terms such as mitochondria dysfunction, chromatin, and synapse in GO analysis, and mTOR pathway, ubiquitin pathway, oxidative phosphorylation, and DNA damage in canonical pathway analysis have also been recognized as dysregulated factors for other psychiatric disorders^38,39^ (Extended Data Fig. 4).

### Cell-type-specific features across cell models with ASD-associated CNVs

To analyze cell-type-specific features in ASD-associated CNVs, we first examined the gene-disease association based on GWAS. Although neuronal cells were enriched in both neuropsychiatric and neurological features, GABAergic (*Gad1+Sst+Npy-*) cluster was especially more remarkable than that of other subtypes (Extended Data Fig. 5). The pathogenesis of neuropsychiatric disorders, including ASD, is highly associated with synaptic dysfunctions^40,41^. Thus, we next focused on the genes related to glutamatergic synapse by analyzing the relationship between glutamatergic cluster-specific genes (GCG) (p < 0.05 and log_2_-fold change > 0.4) and the SFARI genes. We referred datasets of postsynaptic density (PSD) complex proteins^42^, and realized that the PSD complex genes were substantially enriched in glutamatergic cluster (total average 24.0%, 1q21.1 dup, 31.0-32.7%; 2p16.3 del, 12.7-15.0%; 3q29 del, 26.7-27.9%; 7q11.23 dup, 23.7-28.3%; 15q11.2 del, 28.0-29.1%; 15q13.3 del, 22.4-23.8%; 16p11.2 del, 24.7-26.2%; 16p13.2 del, 15.8-27.5%; 17p11.2 del, 21.0-24.8%; 17q12 del, 19.1-22.3%; Xq27.3 del, 24.1-24.8%; Xq28 del, 19.6-24.0%) (Extended Data Fig. 6a).

We then assessed the “upstream regulators” that potentially affect the GCG as their downstream genes by using Ingenuity Pathway Analysis (IPA) (Extended Data Fig. 6b). Overall, neural development and neuronal function-related genes, such as *MAPT, PSEN1*, and *APP* (these are implicated in the etiology of Alzheimer’s disease), *POLG*, a mitochondrial DNA polymerase, *ADORA2A*, an adenosine A(2A) receptor, *MKNK1*, a MAPK interacting serine/threonine kinase, *RTN4*, a potent neurite outgrowth inhibitor, *SOD1*, a major cytoplasmic antioxidant enzyme, and *FMR1*, a negative translational regulator, were highly enriched regardless of cell-type or CNV associated with ASD. Genes significantly enriched in ASD-associated CNVs, such as *MAPT, FMR1, PSEN1, APP, ADORA2A, SOD1, POLG, KMT2A, YWHAZ, CREBZF, SERPINA1, and TP53*, have been reported as ASD risk or susceptible genes.

We performed canonical pathway analysis using IPA (Fig. 3a). Of note, three major translational pathways, EIF2 signaling, regulation of eIF4 and p70S6K signaling, and mTOR signaling, were enriched through all 12 cell lines containing ASD-associated CNVs, followed by mitochondria dysfunctions, oxidative phosphorylation, and protein ubiquitination pathway. These pathways seemed common and convergent regardless of cell type and psychiatric disorders. The analysis also identified cell-type-specific enriched pathways, unfolded protein response (UPR) and endoplasmic reticulum (ER) stress for *Gad1+Sst+Npy+* GABAergic neurons. p53 signaling for non-neuronal cells such as microglia and endothelial cells. These data indicate that p53, a tumor suppressor, is involved in the various biological process regardless of the cell-types^43^ and affects non-neuronal cell lineage in ASD-associated CNVs.

**Figure 3.**
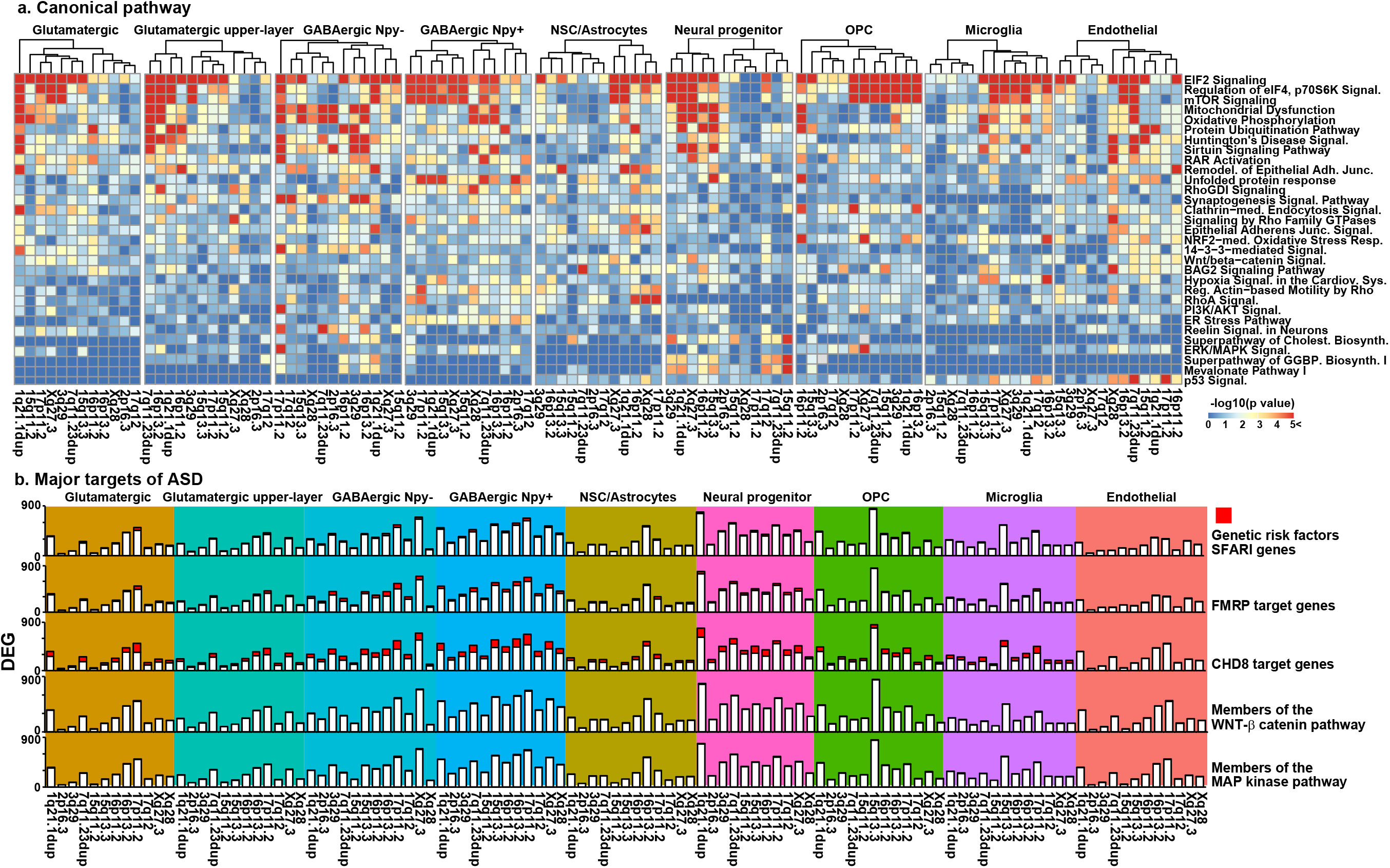
Cell-type and CNV- specific pathway analysis. **a**. Heatmap for the cell-type-specific enriched canonical pathways was performed by using Ingenuity Canonical Pathways Analysis. Each column in the figure represents an ASD-associated CNV, and each row represents a canonical pathway. The colors in the graph indicate the -log_10_ (p-value). **b**. Cell-type-specific single-cell RNA-seq data were used for the analysis. Numbers of DEGs and proportion of genes involved in major targets of ASD (SFARI ASD risk genes, FMRP targets, CHD8 targets, members of WNT β-catenin pathway, and members of MAP kinase pathway, respectively) were analyzed.

We then analyzed major targets of ASD such as FMRP, CHD8, WNT-β catenin, and the MAP kinase pathway^44^ (Fig. 3b). This analysis found cell-type-specific pathway regulation; in particular, FMRP target genes were mostly enriched in neuronal clusters but not in other cell types. Meanwhile, CHD8 target genes were enriched in all cell types except for endothelial cells, although both FMRP and CHD8 protein are expressed ubiquitously.

## DISCUSSION

This study has established an ASD cell bank in which 65 ASD-associated chromosome loci were targeted as a standardized platform for ASD research. These targeted CNVs and genes are highly associated with not only ASD but also other neurodevelopmental and neuropsychiatric disorders such as schizophrenia and bipolar disorder. Thus, the cell models developed in this study contribute to the research for multiple neurodevelopmental and psychiatric disorders. Moreover, the model was established using multipotent mESCs. Thus, as demonstrated in this study, we can generate mouse models of ASD containing CNVs of interest by introducing these mESCs into blastocyst-stage host embryos. It is also possible to differentiate the mESCs into multiple cell lineages of different tissues or organs *in vitro* as well as three-dimensional (3D) organoid cultures. In addition, these cell models allow us to perform transplantation and blastocyst complementation studies^9^. Therefore, our cell models would be valuable tools for various organogenesis studies with disease modeling *in vivo* and *vitro*.

Recent advances in genome editing techniques enable us to generate mutant animal models faster and efficiently than before. With genome editing tools such as the CRISPR/Cas9 system, gene targeting can be achieved directly in zygotes via microinjection of genome editing reagents without requiring mutant ES cells and the injection step of the mutant ES cells into the blastocysts^45^. The latest advances in genome editing tools can generate mutant mice within a single generation without the use of ES cells. Our crRNA (Crispr RNA) sequences designed *in silico*^11,46^ are verified by *in vitro* Surveyor assay. They can be used to synthesize oligonucleotides as crRNA, which can be utilized for zygote injection. Our original protocol named “next-generation chromosome engineering” designed to target a specific chromosome locus of interest successfully generated mega-base scale CNVs of the chromosomal regions such as 3p14.1 (3.4 Mb), 3p26.3 (3.2 Mb), and 15q11-q13 (6.3 Mb).

Previous bulk transcriptome analyses identified converging biological processes and pathways of ASD^35^. A cell-type-specific study using patient samples suggested the importance of synaptic and neurodevelopmental genes in cortical upper layer 2/3 and clinical association with microglia^29^. In this study, using scRNA-seq analysis, we (1) identified CNV- and cell-type heterogeneity of ASD (17 cell-types from pathogenic 12 different CNVs); (2) confirmed the importance of synaptic, PSD complex, and neurodevelopmental genes on ASD pathogenesis; (3) showed that FMRP targets were enriched in neuronal cells, whereas CHD8 targets were broadly affected in various cells including non-neuronal cells, although both genes were ubiquitously expressed; (4) showed a significance of neuronal genes as upstream regulators; (5) identified enriched terms implicated in ASD, such as translation, transcriptional regulation, cell cycle, morphogenesis, and immune system; (6) showed two faces of ASD-associated CNVs, psychiatric and neurological disorders

Our isogenic cellular platform for ASD research can be extended for 3-dimensional culture organoid, imaging, and transplantation as well as drug screening. In addition, our targeting and CRISPR/Cas9 vector collection and validated genome editing information could contribute to generating novel mouse models with CNVs. Our biological resource can be accessed through the web browser: https://www.med.kobe-u.ac.jp/asddb/

## Acknowledgments

We thank T. Yoshikawa and M. Toyoshima for the neural differentiation method, N. Mataga, K. Fukumoto, F. Sakai, N. Ito, C. Nishioka, M. Kadota and A. Kwon for experimental supports, Y. Okumura and M. Toki for database development, and all technical staff of the Takumi Lab for their technical assistance. This work was supported in part by KAKENHI (16H06316, 16H06463, 16F16110, 17K07119, 21H00202, 21H04813, 21K07820, 21K19351) from Japan Society for the Promotion of Science (JSPS) and Ministry of Education, Culture, Sports, Science, and Technology, Japan Agency for Medical Research and Development (JP21wm0425011), Intramural Research Grant (30-9) for Neurological and Psychiatric Disorders of NCNP, Takeda Science Foundation, Smoking Research Foundation, SENSHIN Medical Research Foundation, Tokyo Biochemical Research Foundation, Research Foundation for Opto-Science and Technology, Hyogo Science and Technology Association, Kawano Masanori Memorial Public Interest Incorporated Foundation for Promotion of Pediatrics, Taiju Life Social Welfare Foundation, and Naito Foundation. AZ was supported by JSPS Postdoctoral Fellowship for Research in Japan.

## Author contributions

J.N. and T.T. conceived and designed the study. T.T. supervised this project. J.N. drafted the manuscript. J.N. developed & implemented a chromosome manipulating technique. J.N., K.K., C.M., H.M., Y.N., Y.K., A.H., K.M., I.S., R.F., E.B., and K.Y., developed ASD cell models and vector library. A.Z., K.F., H.M., Y.S., I.S., Y.N., K.K., Y.K., C.M., N.N., and J.N. performed neural morphological and physiological experiments. T.A., E.T., J.N., K.K., C.M., Y.K., A.H., H.M., K.M., and K.Y., prepared ES cells for blastocyst injection and generated 15q13.3 chromosome mutant mice. J.N., M.S., Q.E., R.F., and C.M. performed behavioral tests. Y.K., C.M., and Y.N. performed cDNA library preparation. J.N., T.K., H.M., S.F., S.K., P.C., and J.S performed NGS and scRNA-seq analysis. J.N., H.M., Y.K., C.M., Y.N., A.H., K.K., K.M., R.F., X.L., and T.T. developed ASD CNV database.

## METHODS

### Cell culture

CMTI-2 mouse embryonic stem cell line (derived from murine strain C57BL/6J, normal male genotype, Millipore) and their derivatives were used for all experiments. mESCs were cultured in Dulbecco’s modified Eagle’s medium (DMEM, Gibco) supplemented with ES-qualified 15% FBS, 2 mM L-glutamine (Gibco), 0.1 mM MEM nonessential amino acids (NEAA, Gibco), nucleoside (Adenosine, Guanosine, Cytidine, Uridine, and Thymidine 30 µM each, Sigma), 0.1 mM 2-mercaptoethanol (Sigma), and 1,000 IU/ml LIF (ESGRO, Millipore) on mitomycin C-treated mouse embryonic fibroblasts (MEF) feeder cells. MEF cells were grown in Glasgow minimum essential medium (GMEM, Sigma) supplemented with 10% FBS.

### Establishment of ES cell lines from a blastocyst

Preparation of 15q11-q13 mouse ES cells from blastocyst was established as previously described with some modification^47^. Briefly, fertilized embryos were collected from C57BL/6J female mice mated with 15q11-q13 duplication males for paternal 15q11-q13 duplication, or 15q11-q13 duplication female mice mated with C57BL/6J male mice for maternal 15q11-q13 duplication, respectively. The separated cells were transferred to the individual wells of 96 well plates coated with MEFs. ES cell establishment medium with 20% Knockout Serum Replacement (KSR; Invitrogen), 0.1 mg/ml adrenocorticotropic hormone (ACTH; fragments, American Peptide Company), instead of fetal bovine serum (FBS). After 10 days, proliferating outgrowths were dissociated and cultured until stable cell lines grew out.

### Vector construction

CRISPR-Cas9 vectors for genome editing, pX330 (Addgene plasmid # 42230) was used. Annealed oligonucleotide for sgRNA was inserted into the BbsI site of pX330. Target sgRNAs were designed using the CRISPR design tool (http://crispr.mit.edu/) or CRISPRdirect (https://crispr.dbcls.jp/). To avoid off-target effects, only a higher score (>80; CRISPR design tool) or highly specific sequence (CRISPRdirect) were selected. The targeting vector cassette (DT-A/Neo #09, RIKEN BDR) contains a neomycin resistance gene and a DT-A cassette to avoid random integration. 5’ and 3’ homology arms were inserted into the outside of the neomycin resistance gene cassette in this vector. Target-specific crRNA sequence and primer set to make homology arms for targeting vectors are listed in Supplementary Table 2. All clones used in this study are available from RIKEN Bioresource Center, Japan (BRC, Gene engineering division: https://dna.brc.riken.jp/en/).

### Chromosome manipulation in mouse ES cells

The donor targeting vector was linearized by Asp718 (Roche). Mouse ES cells, CMTI-2 (Millipore), were electroporated using the Nucleofector II (Lonza). Program #A-23 was used for all electroporation. In each experiment, 5 × 10^6^ trypsinized cells were resuspended in 93.5 µl solution (mouse ES cell nucleofector kit, VPH-1001, Lonza). The solution includes 2 µg of each CRISPR/Cas9 vector (4 µg total) and 20 µg of the linearized targeting vector. After mixing with the DNA solution, the suspension was transferred to cuvettes, and the vectors were quickly electroporated into the cells. On the second day post electroporation, 500 µg/ml G418 (Nacalai Tesque) was applied to select neomycin-resistant ES cells, and each colony was cultured in a 96 well plate. Chromosome modification was verified by Southern blotting, PCR, quantitative RT-PCR, or array CGH.

Successfully targeted cell clones in this study are listed in Extended Data Table 4. All clones used in this study are available from RIKEN Bioresource Center, Japan (BRC, Cell engineering division: https://cell.brc.riken.jp/en/).

### Array comparative genomic hybridization

According to the manufacturer’s protocol, the Array comparative genomic hybridization (aCGH) was performed using SurePrint G3 Mouse CGH Microarray Kit, 1 × 1 M (Agilent Technologies). Genomic DNA from wild-type mouse ES cells (CMTI-2) was used as a reference. Signals were then analyzed using R.

### Neural differentiation from mouse ES cells

After being cultured on gelatin-coated dishes, mESCs were used to form an embryoid body. Cells were cultured on a nonadherent bacterial dish for 8 days in this suspension culture condition. Retinoic acid (5 µM) was added on days 4 and 6, respectively^48^. After suspension culture, cells were dissociated and seeded onto the Poly-L-ornithine (PLO) and laminin-coated culture dish, and then started differentiation by using neuronal media including NeuroCult NSC Basal Medium (StemCell Technologies) supplemented with 2% (v/v) B27 (Life Technologies), 10 ng/ml brain-derived neurotrophic factor (R&D Systems), 10 ng/ml glial-derived neurotrophic factor (R&D Systems), 200 μM ascorbic acid (Sigma-Aldrich) and 400 μM dibutyryl-cAMP (Sigma-Aldrich)^49^. Typical neuronal morphology was observed within 2 days.

### Immunocytochemistry and morphological analysis

Differentiating neurons were transfected on day 1 with pβactin-GFP by lipofectamine LTX and PLUS Reagent (Thermo Fisher Scientific) and cultured for additional 2 days on laminin and PLO-coated glass coverslips. Immunocytochemical staining was performed on day 3. In this step, cells were fixed with 4% paraformaldehyde containing 4% sucrose for 20 min at 37°C. After fixation, cells were washed twice with PBS for 10 min. Then, cells were blocked with blocking buffer (2% goat serum, 1% Glycine, 0.1% Poly-D-Lysine, 0.3% Triton, 1% BSA in PBS) for 1 h. Then, cells were incubated with primary antibody (mouse anti-GFP, 1:1000 dilution, Thermo Fisher Scientific) in blocking buffer in a humidified chamber overnight at 4 °C. The next day, cells were washed three times with PBS for 10 min and then incubated with the secondary antibody (goat anti-mouse Alexa 488, 1:1000 dilution, Thermo Fisher Scientific) in a blocking buffer for 1.5 h. After washing three times with PBS, nuclei were stained with DAPI (VECTOR Laboratories) for 10 min. Cells were then washed once with PBS for 10 min and were mounted with FluorSaveTM Reagent (Sigma-Aldrich) overnight at room temperature. All following steps were performed as described previously^50^. Images were taken by BZ-9000 (Keyence). Axon length, total neurite length, and the branch numbers were analyzed on WIS-Neuromath (Weizmann Institute of Science, Israel). Twenty neurons were randomly chosen.

### Ca^2+^ imaging

Differentiated neurons were cultured on laminin and PLO-coated glass bottom-culture dish. The neurons cultured for 7-days on the dish were treated with 1 μM of the Ca^2+^- sensitive, membrane-permeable fluorescent dye Fluo-4-AM (Thermo Fisher Scientific) dissolved in recording buffer (135 mM NaCl, 5 mM KCl, 2 mM CaCl_2_, 2 mM MgCl_2_, 10 mM HEPES, 10 mM glucose, at pH 7.4) for 30 min at 37 °C. After loading, cells were washed three times with the recording buffer at room temperature. Green fluorescence images were acquired using an EMCCD camera (iXon Ultra 897, Andor) on a confocal microscope (IX81 FV1000, Olympus) through a 20× objective lens (Olympus) and 495-540 nm emission filters (Olympus). Time-lapse images of the neurons at room temperature were recorded for 24 s duration with 20 ms interval by MetaMorph (Molecular Devices). The neurons were stimulated by 25 mM KCl manually treated in the recording buffer at the middle time point of recoding (i.e., at 12 s). Images from randomly selected 20 neurons were analyzed by Image J. Fluorescent intensities of the soma were averaged to represent the signal of the neuron *F*. This value was divided by the baseline signal value *F0*, which was calculated as an average of *F* before KCl treatment (i.e., 0 ∼ 10 s), to obtain normalized fluorescence changes ΔF/F = (F-F0)/F0.

### Southern blotting

Genomic DNA (10 µg) was digested with EcoRI for the 5’ region and HindIII for the 3’ region, respectively. The digested DNA was electrophoresed on a 0.8% agarose gel and transferred to a Hybond-N+ membrane (GE Healthcare). The membrane was hybridized with a digoxigenin-labeled DNA probe generated with a PCR DIG Probe Synthesis Kit (Sigma). Hybridization with a 5’ probe produced an 8.1 Kb band from the WT and a 5.1 Kb band from the targeted locus, while hybridized with a 3’ probe produced a 12.4 Kb band from the WT and an 8.1 Kb band from the targeted locus.

### Animals

Mice were housed in a room with a 12-hour light/dark cycle (light on 8:00 a.m. and off 8:00 p.m.) and provided *ad libitum* access to water and food. All protocols for animal experiments were approved by the Animal Care and Use Committees of the RIKEN Brain Science Institute and performed under the institutional guidelines and regulations.

### Generation of 15q13.3 chromosome mutant mice

To generate human 15q13.3 microdeletion model mice, C57BL/6 background ES cells with 15q13.3 heterozygote deletion were microinjected into BALB/c blastocysts. Then, sperm from the chimera mouse were fertilized *in vitro* (IVF) to generate 15q13.3(+/-) mice. Genotype was confirmed by Southern blot and three-primer PCR. PCR primer used for genotyping were as follows: L-Neo1, 5’ - GTACTCGGATGGAAGCCGGTCTTGTC-3’, 9_15q13.3_Wt_Fw, 5’- ACGCAGGGTGTAGAAGCAAA-3’, and 9_15q13.3_Wt_Tg_Rv, 5’ - CCGGTCGATTGTGAGTTCA-3’.

PCR with 9_15q13.3_Wt_Fw and 9_15q13.3_Wt_Tg_Rv primer pair produces a 395-bp fragment from the wild-type locus, whereas the L-Neo1 and 9_15q13.3_Wt_Tg_Rv primer pair have a 999-bp fragment from the targeted locus. PCR was performed with the Taq DNA Polymerase (NEB) following the manufacturer’s protocol. The PCR program consisted of initial denaturation at 96°C for 1 min; 40 cycles of 96°C for 10 s, 60°C for 10 s, and 68°C for 1 min.

### RT-qPCR analysis

Total RNA (0.5 μg) isolated from mouse cortices was subjected to reverse transcription (RT) with a SuperScript II Reverse Transcriptase (Thermo Fisher Scientific). The cDNA was subjected to quantitative PCR using SYBR Green PCR Master Mix (Applied Biosystems) and specific primers in a StepOnePlus (Applied Biosystems). PCR primer sequences (sense and antisense, respectively) were as follows: Chrna7, 5’-GCATGAAGACAGTCAGAGAAAGTAA -3’ and 5’-CCCTGGCTTTGCTGGTATT -3’; Otud7a, 5’-TCTTCCTTCGCCTCATGC -3’ and 5’-CACCTCCAGAGAGTGAGGAGTC -3’; and β-actin, 5’-TGGATGCCACAGGATTCCAT -3’ and 5’-CGTGCGTGACATCAAAGAGAA -3’. The amount of *Chrna7* and *Otud7a* were normalized using β-actin.

### Behavioral experimental design

C57BL6/J strain mice older than 10 weeks were used for behavioral tests. In the cage, 2-5 mice were housed together with littermates. Mice were maintained in a 12-h light/dark cycle (light on at 8:00) with *ad libitum* access to water and food. All behavioral tests were performed between 9:00 AM and 6:00 PM. Mice were habituated to the testing room for at least 30 min before starting the behavioral experiments to allow acclimatization.

### Open field test

Locomotor activity was measured in the open field apparatus (50 × 50 × 30 cm, O’Hara) illuminated at 100 lux light for 1 hour. Total distance traveled, time spent in the center area, and the rearing were recorded using TimeOFCR4 (O’Hara). The center area was defined as 36% of the field.

### Y-maze spontaneous alternation test

Spatial working memory was measured by the Y-maze test. The Y-maze apparatus consists of three arms. The apparatus was illuminated at 100 lux lights for 5 min. Each mouse was allowed to explore freely in the Y-maze in this test. The alternation rate was analyzed using TimeYM2 for Y-maze (O’Hara).

### Ultrasonic vocalization (USV)

Postnatal day 6 (P6) pups were assessed for ultrasonic vocalization (USV). A pup separated from their mother and littermates was put into a plastic beaker (8 cm diameter, 12 cm height, the bottom covered with gauze) with a condenser ultrasound microphone (CM16/CMPA, Avisoft Bioacoustics) and placed in a soundproof box. USVs were recorded for 5 min at a sampling rate of 250 kHz. The recorded data were analyzed using SASLab Pro (Avisoft Bioacoustics).

### Grooming

Mice were placed in a clean, transparent plastic cage. The self-grooming behavior was recorded by a video camera for a 10 min test period following 10 min habituation. The amount of time spent self-grooming was counted using a stopwatch.

### Acoustic startle response

Mice were placed in a Plexiglas cylinder and acclimated for 5 min. A test session was composed of 49 trials, and each trial was composed of prepulse sounds (0, 72, 74, 78, 82, and 86 dB respectively) pulse - (120 dB) paired stimulus or a no prepulse - no pulse pair. The average acoustic startle response was calculated by no prepulse – 120 dB pair.

### Elevated plus maze test

The elevated plus-maze apparatus (O’Hara) consists of two open arms (25 × 5 cm) and two enclosed arms of the same size with 15 cm high transparent walls. The arms and central square were made of white plastic plates and were elevated to a height of 55 cm above the floor. Mouse behavior was recorded during a 10 min test period. Anxiety-like behavior was measured by the percentage of time spent in the open arms. The maze was illuminated with 100 lux.

### Three-chamber social interaction test

The testing apparatus consisted of a rectangular three-chamber box (O’Hara). Mice can move to each chamber (20 × 40 × 20 cm) through small square openings (5 × 3 cm). The apparatus was illuminated at 10 lux. This test consists of three sessions. The first session for habituation to the apparatus for 10 min. Then, an age-matched unfamiliar male mouse (C57BL/6J) was placed in the wire cage in one of the two side chambers, and the subject mouse was allowed to move freely in the test box for 10 min to assess sociability. Finally, a second unfamiliar male mouse (C57BL/6J) was placed in the wire cage at the other side of the chamber to assess social novelty. In this session, the previously used stranger mouse was considered a familiar mouse. The subject mouse was placed in the chamber and allowed to move freely in the test box for 10 min. Time spent in the area was analyzed using TimeCSI1 for the three-chamber social interaction test system (O’Hara).

### Flurothyl-induced seizures

Flurothyl-induced seizure experiments were performed according to the previous study^51^. Briefly, the mice were placed individually in an air-tight glass chamber (2 L volume) in a ventilated chemical hood and then habituated to the chamber for 1 min. After habituation in a glass chamber, the mouse was exposed to 10% flurothyl (bis(2,2,2-trifluoroethyl) ether) in 95% ethanol. 10% Flurothyl solution was infused through a 5 ml syringe by using a microsyringe pump (KD Scientific) onto a gauze pad at the top of the chamber at a rate of 200 µl/min. We analyzed seizure behaviors using a video camera. In the case of observation of a generalized seizure, we immediately removed the lid of the chamber, exposing the mouse to fresh air to stop the seizure assay.

### Single-cell cDNA library preparation and RNA-sequencing (scRNA-seq)

Differentiated cells on day 15 (8 days suspension culture plus 7 days adherent culture) were harvested and dissociated using TrypLE Select (Gibco). Cells were then passed through a 20 µm strainer and resuspended in 1 x PBS with 0.04% BSA buffer. Then, cell suspensions (concentration and viability were assessed using TC20 Automated Cell Counter (Bio-Rad)) were loaded on a Chromium Single Cell Controller (10x Genomics) to generate single-cell gel beads in emulsion (GEMs) by using Chromium Single Cell 3’ Library and Gel Bead Kit v2 (10x Genomics) following the manufacture’s introduction. Captured cells were lysed and the RNA was barcoded through reverse transcription in each GEM. Reverse-transcribed cDNAs were purified by using Myone DynaBeads. To ensure successful amplification and accurate concentration of cDNA, we used Agilent Bioanalyzer 2100 using a High Sensitivity DNA chip (Agilent). Post library construction quantification was performed by KAPA Library Quantification Kits (Roche) according to the manufacture’s protocol. Sequencing was performed on an Illumina HiSeq 2000 or 1500 with pair-end using the following read length: 26 cycles Read1, 8 cycles i7 Index, and 98 cycles Read2.

### Single-cell sequence data pre-processing

The Illumina sequencer’s base call files (BCLs) were demultiplexed and converted to sample-specific FASTQ files using the cellranger mkfastq pipeline. Then, raw reads were processed using cellranger count pipeline, which takes FASTQ files generated by cellranger mkfastq and performs alignment to the mouse reference data (mm10), filtering, barcode counting, and Unique Molecular Identifier (UMI) counting. All scRNA-seq data were processed using R (v3.6.1). The datasets were processed following the pipeline of the Seurat (v3.0). Cells with unique feature counts of more than 2,500 or less than 200 and more than 5% mitochondrial counts were filtered out. Finally, a total of 14,396 cells (Ctrl, 1159; 1q21.1 dup, 969; 2p16.3, 655; 3q29, 738; 7q11.23 dup, 907; 15q11.2, 357; 15q13.3, 987; 16p11.2, 996; 16p13.2, 1112; 17p11.2, 750; 17q12, 2057; Xq27.3, 2361; Xq28, 1348 cells) were used for downstream analysis. The 13 datasets were integrated using the Seurat pipeline and performed dimensionality reduction for visualization by using Uniform Manifold Approximation and Projection (UMAP). For bulk analysis, cellranger aggr pipeline was used to identify each CNV’s features by calculating each gene expression by 10x Genomics Loupe (TM) Cell Browser v.2.0.0 (https://support.10xgenomics.com/single-cell-gene-expression/software/visualization/latest/what-is-loupe-cell-browser).

### Gene ontology (GO) and pathway analysis

Gene lists were submitted to the Metascape software (http://metascape.org/gp/index.html#/main/step1) for functional Annotation^52^. Enrichment terms from Biological Process (BP), Cellular Component (CC), and Molecular Function (MF) as GO analysis, as well as KEGG (Kyoto Encyclopedia of Genes and Genomes), were examined for further enrichment analysis. Enrichment network visualization was used to detect enriched term network. ToppFun software in the ToppGene Suite (https://toppgene.cchmc.org/enrichment.jsp)^53^ was used to analyze DEGs to GWAS-based gene-disease associations. The Ingenuity Pathway Analysis (IPA: Qiagen Bioinformatics) was used for upstream and canonical pathway analysis. SFARI genes and scoring modules were referred 06-20-2019 version of the SFARI database.

### Statistics

Statistical analysis was conducted using R, and data were analyzed using one-way analysis of variance (ANOVA), two-way repeated-measures ANOVA. Bonferroni correction was applied to multiple comparisons. The significance level was set to p < 0.05.

### Data Availability

## Extended data figures and tables

**Extended Data Figure 1. Generation and characterization of mice mimicking human 15q13.3 microdeletion syndrome**

**a**. Mouse C57/BL6J strain CMTI-2 ES cell line with chromosome 7 (7qC) heterozygote deletion were grown on mitotically inactive primary mouse embryonic fibroblasts (MEF) were injected into 3.5-day old blastocysts and implanted into pseudo-pregnant BALB/c female mice. Chimera rate was determined by coat color. **b**. Male chimera mice with an approximately 60% coat color contribution of CMTI-2 ES cells. **c**. Genotypes of ES cells determined by PCR. PCR with the three primers produces a 395-bp band form the wild-type (WT) and a 999-bp band from the targeted locus. **d**. Quantitative real-time PCR (qRT-PCR) analysis. Total RNA was purified from male adult WT and 15q13.3 (+/-) mouse cortices, respectively. Expression level of *Chrna7* and *Otud7a* were normalized by *β-actin*. n = 3 mice/genotype were used for the analysis. Both *Chrna7* and *Otud7a* expression levels were significantly reduced in 15q13.3(+/-), *Chrna7*, F(1,4) = 27.344, p < 0.01; Otud7a, F(1,4) = 21.577, p < 0.01 **e**. Body weights growth curves of WT and 15q13(+/-) mice. In the test period, significant difference was observed between two genotypes. F(1, 285) = 4.453, p < 0.0357, two-way repeated ANOVA (WT, n = 30; 15q13.3(+/-), n = 29). **f**. Brain weights were assessed at 5 months age. No significant difference was observed between WT and 15q13.3(+/-). F(1, 19) = 0.17, p = 0.685, one-way ANOVA (n = 3 for each genotype). **g**. Open field test to analyze locomotor activity, anxiety, vertical activity in the novel environment. There is no significant difference between genotypes, total distance, F(1,38) = 0.203, p = 0.655; percentage of time spent in the center area, F(1,38) = 0.066, p = 0.799; and the number of rearing in the open field chamber, F(1,38) = 0.188, p = 0.667 (n = 20 for each genotype). **h**. Spontaneous alternation Y-maze test to analyze spatial working memory. There is no significant difference between genotypes, F(1,38) = 1.174, p = 0.285 (n = 20 for each genotype). **i**. Ultrasonic vocalization test (USV) to analyze an early communicative behavior between mother and their dams. There is no significant difference between genotypes, F(1,68) = 0.527, p = 0.47 (WT, n = 36; 15q13.3(+/-), n = 34). **j**. Grooming number was counted to analyze core autistic symptoms (e.g. repetitive and excessive self-grooming). There is no significant difference between genotypes, F(1,42) = 0.052, p = 0.82 (n = 20 for each genotype). **k**. Acoustic startle response was assessed to see an exaggerated startle response to an unexpected auditory stimulus. Startle response was significantly increased in 15q13.3(+/-) mice. F(1,38) = 5.622, p < 0.05 (n = 20 for each genotype). **l**. Elevated plus maze test was performed to analyze anxiety-like behavior. There is no significant difference between genotypes, F(1,38) = 0.098, p = 0.756, F(1,38) = 0.219, p = 0.642, closed and opened arms respectively (n = 20 for each genotype). **m**. Three-chamber social interaction test to assess sociability. Both ctrl and 15q13.3(+/-) mice significantly stay longer in stranger cage than empty cage, F(1,32) = 8.21, p < 0.01, F(1,38) = 48.69, p < 0.01, respectively. Although ctrl mice stay longer in stranger mice than familiar mice area, F(1,32) = 4.602, p < 0.05, no significant difference was observed in 15q13.3(+/-) mice, F(1,38) = 0.448, p = 0.507 (WT, n = 17; 15q13.3(+/-), n = 20). **n**. Schematic of GABA _A_R antagonist, flurothyl-induced seizure protocol and experimental setting. **o**. Latency to generalized seizure were analyzed in both genotypes and genders. Two-way ANOVA revealed no significant main effect of genotype (F(1, 52) = 0.073, p = 0.78) and gender (F(1,52) = 0.113, p = 0.738), and no significant interaction between genotype and gender (F(1, 52) = 3.08, p = 0.085) (Male WT, n = 9; male 15q13.3(+/-), n = 14; female WT, n = 17; female 15q13.3(+/-), n = 16). **p**. Frequency of generalized seizure were analyzed in both genotypes and genders. Two-way ANOVA revealed significant main effect of genotype (F(1, 52) = 4.923, p < 0.05), but not in gender (F(1,52) = 0.004, p = 0.95) (Male WT, n = 9; male 15q13.3(+/-), n = 14; female WT, n = 17; female 15q13.3(+/-), n = 16).

**Extended Data Figure 2. Analyzed targeted deletion or duplication of the cellular model of ASD by array-CGH**

Genomic DNA was extracted from control and twelve representative mutant mouse ES cells, respectively. Control genomic DNA was used as a reference. Each dot represents an oligonucleotide. A red shaded region indicates a deleted or duplicated region. Analysis was designed by referring to the mouse reference sequence mm9 (NCBI Build 37).

**Extended Data Figure 3. Morphological and physiological analyses of neurons derived from mES cells**.

**a, b, c, d**. Box plots of the axon length **(a)**, total neurite length **(b)**, and total branch number **(c)** in mouse ESC-derived neurons, respectively. (total n=1348 cells including; n=179 (Ctrl); 75 (1q21.1 dup); 82 (2p16.3); 155 (3q29); 69 (7q11.23 dup); n=79 (15q11.2); 160 (15q13.3); 92 (16p11.2); 65 (16p13.2); 71 (17p11.2); 82 (17q12); 85 (Xq27.3); 154 (Xq28)). **d**. Traces show the relative change in fluorescence intensity (ΔF/F) induced by 25 mM KCl at day 3. **e**. The averaged peak amplitude of Ca2+ response (ΔF/F) was evoked by 25 mM KCl. Box plots show median, quartiles (boxes), and range (whiskers). P-values are determined by One-way-ANOVA with post hoc Bonferroni multiple comparison test (**a, b, c, e**). ***p < 0.001.

**Extended Data Figure 4. Functional and Molecular signature of ASD-associated CNVs**.

**a-c**. Gene ontology (GO) analysis for clustering of CNVs. Biological Process (BP)**(a)**, Molecular Function (MF)**(b)**, and Cellular Component (CC)**(c)**, respectively. The rows of the heatmap represent the GO terms and the columns represent CNVs. **d**. Canonical pathway analysis was performed by using Ingenuity Canonical Pathways Analysis. The rows of the heatmap are the canonical pathway and the columns indicate CNVs. The gradient of color in the heatmap indicates the enrichment levels. The heatmap color indicates statistical significance (-Log10(p-value)).

**Extended Data Figure 5. Gene-disease association analysis identified psychiatric and neurological aspects of ASD-associated CNVs**.

The heatmap shows the interaction between neuropsychiatric, neurological disorders, and ASD-associated CNVs in a cell-type-specific manner. The rows of the heatmap are the major psychiatric and neurological disorders, and the columns are ASD-associated CNVs. The red line indicates a member of the psychiatric disorders, while the green line indicates a member of neurological disorders. The gradient of color in the heatmap indicates the enrichment levels. The heatmap color indicates statistical significance (-Log10(p-value)).

**Extended Data Figure 6. Significance of the glutamatergic postsynaptic density genes and upstream regulator in ASD**.

**a**. Overlap between glutamatergic (*Slc17a6*+) cell-cluster (Figure 2b, #8 and #10) specific DEGs and postsynaptic density (PSD) genes. The X-axis represents the number of DEGs in each cell cluster and Y-axis represents glutamatergic neuronal cells clusters in each CNV. The red column indicates PSD complex genes. SFARI genes (gene scores 1 to 3 and syndromic) in each DEGs are listed to the right. The numbers in parentheses indicate the risk gene score of ASD defined by SFARI, and genes without score indicate a syndromic gene. **b**. Cell-type-specific upstream-regulators. The numbers in parentheses indicate the risk gene score defined by SFARI, and genes without a score are not SFARI ASD risk genes. The rows of the heatmap are the upstream regulator genes, and columns are CNVs. The gradient of color in the heatmap indicates the enrichment levels. The heatmap color indicates statistical significance (-Log10(p-value)).

**Extended Data Table 1. Major CNVs associated with ASD**.

**Extended Data Table 2. Common CNVs among psychiatric disorders (targeted CNVs in the study)**

**Extended Data Table 3. Common genes among psychiatric disorders (target CNVs in our study)**.

**Extended Data Table 4. Cell library**.

**Extended Data Table 5. Consequence of chromosome targeting**.

**Extended Data Table 6. Synteny analysis (targeted CNVs)**.

**Extended Data Table 7. Gene expression in targeted loci**.

Gene expression of each cell-line. Genes located in targeted region were analyzed. Average expression, logFC, p-value were derived from scRNA-seq data. P-values were adjusted using the Benjamini-Hochberg correction for multiple tests. *p < 0.1, **p < 0.05, ***p < 0.01, ****p < 0.001.

**Extended Data Table 8. Cell-type (cluster) specific Gene Ontology (GO) terms**.

**Extended Data Table 9. Gene Ontology analysis; ASD associated 12 CNVs**.

**Supplementary Table 1. Synteny analysis; low syntenic CNV.**

**Supplementary Table 2. CRISPR and targeting vectors used in the study.**

**Supplementary Table 3. Cell-type markers in each cell cluster.**

