## Extended Data Fig. 1 for "Autism in a dish: ES cell models of autism with copy number variations reveal cell-type-specific vulnerability"

### Extended Data Figure 1.

a

| Locus | Cell ID# | Chimera ID# | Chimera rate (%) | # Progeny screened | Germline transmission |
|---|---|---|---|---|---|
| 15q13.3 | 19_A12 | 160212 | 0 | - | - |
| | | 160311 | 60 | 117 | 1 (0.8%) |

b

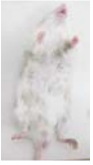

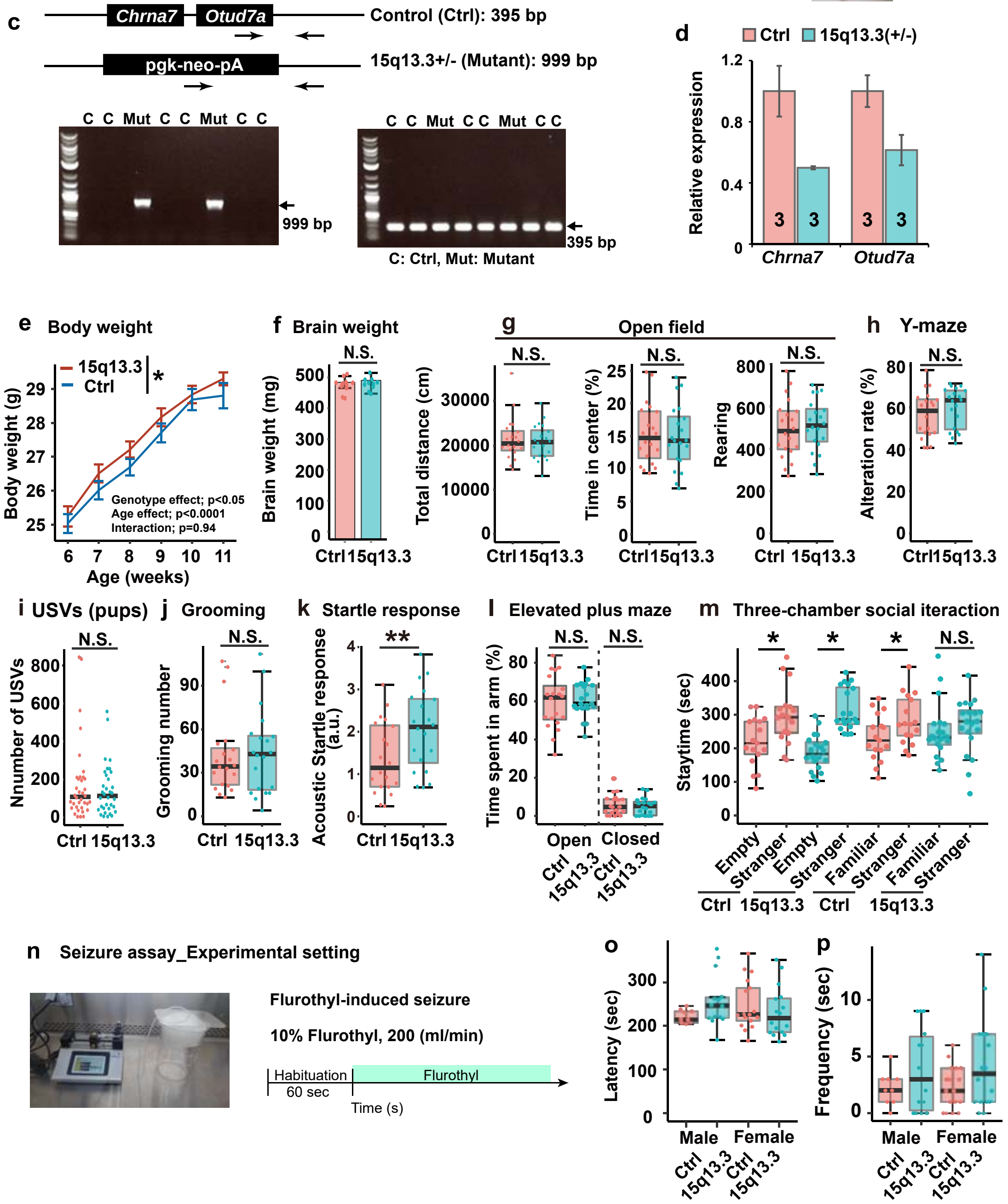
