## Supplementary figures and images for "Autism in a dish: ES cell models of autism with copy number variations reveal cell-type-specific vulnerability"

### Extended Data Fig. 2-1

#### Extended Data Figure 2.1.

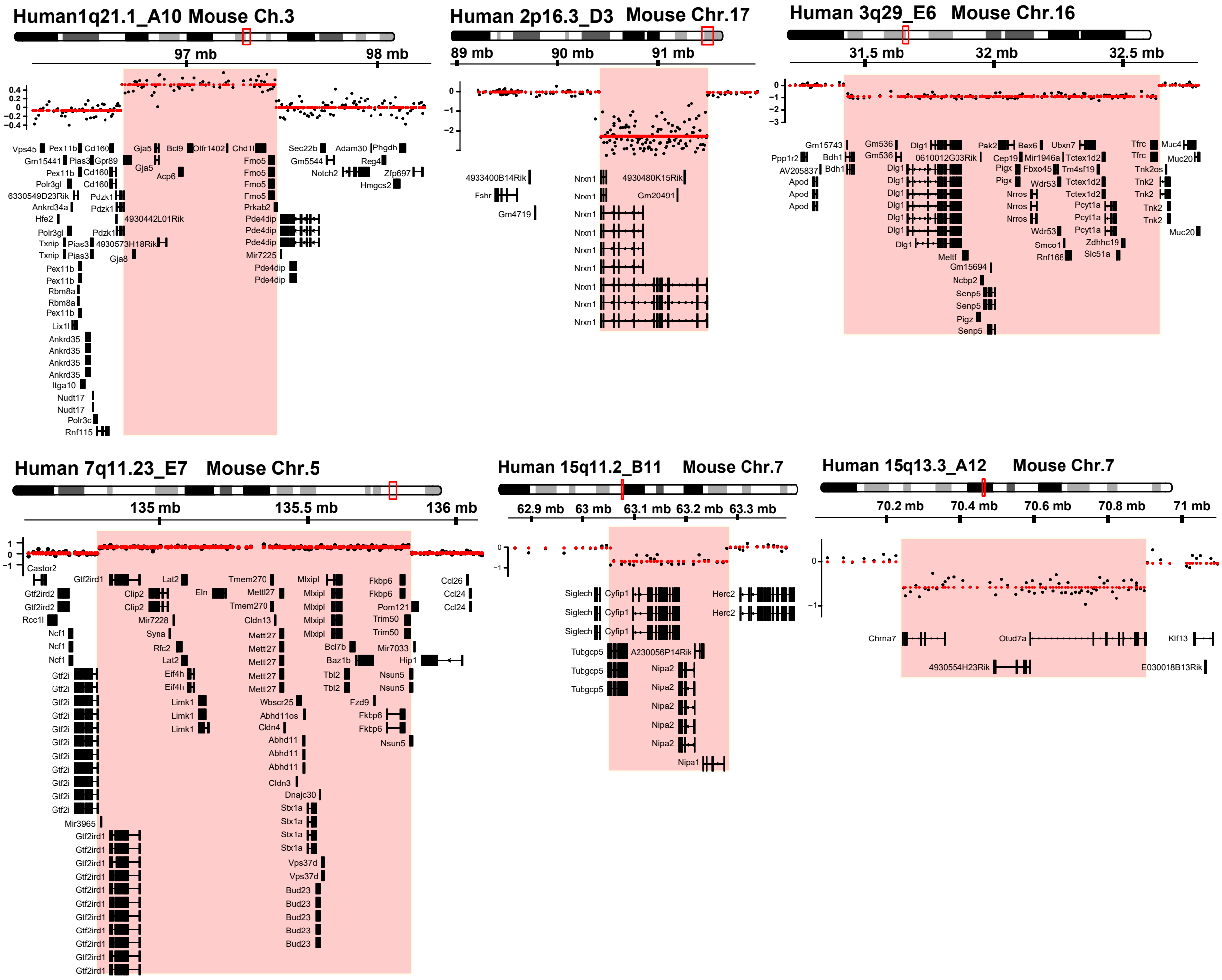

### Extended Data Fig. 2-2

#### Extended Data Figure 2.2.

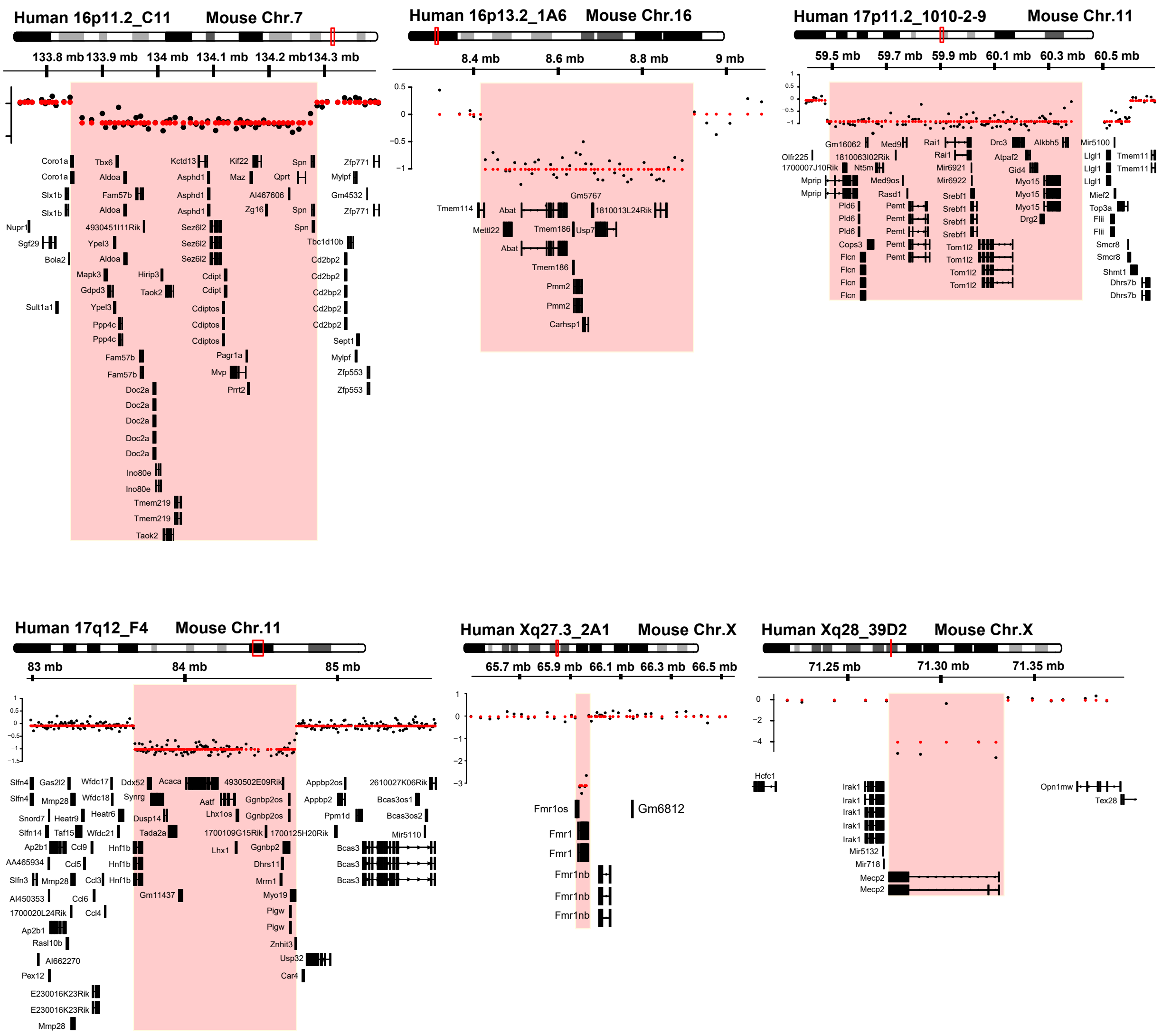

### Extended Data Fig. 3

**Extended Data Figure 3.**

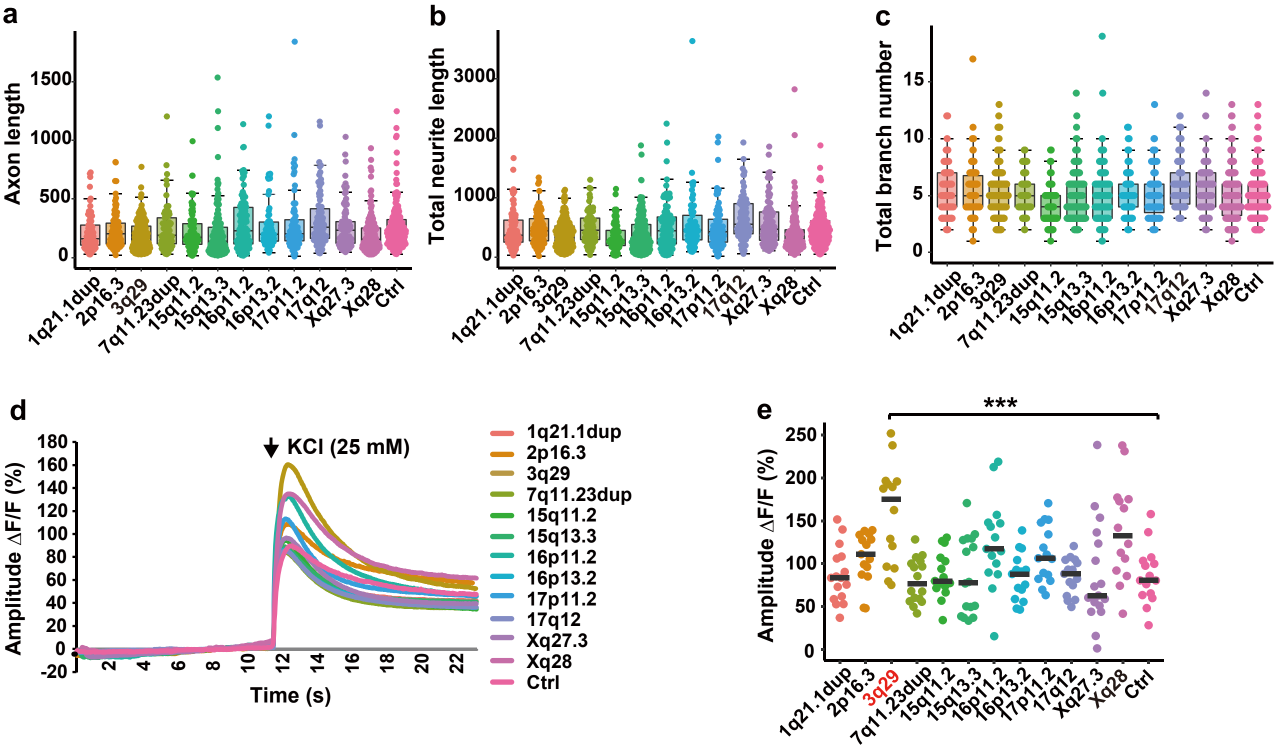

### Extended Data Fig. 4

#### Extended Data Figure 4.

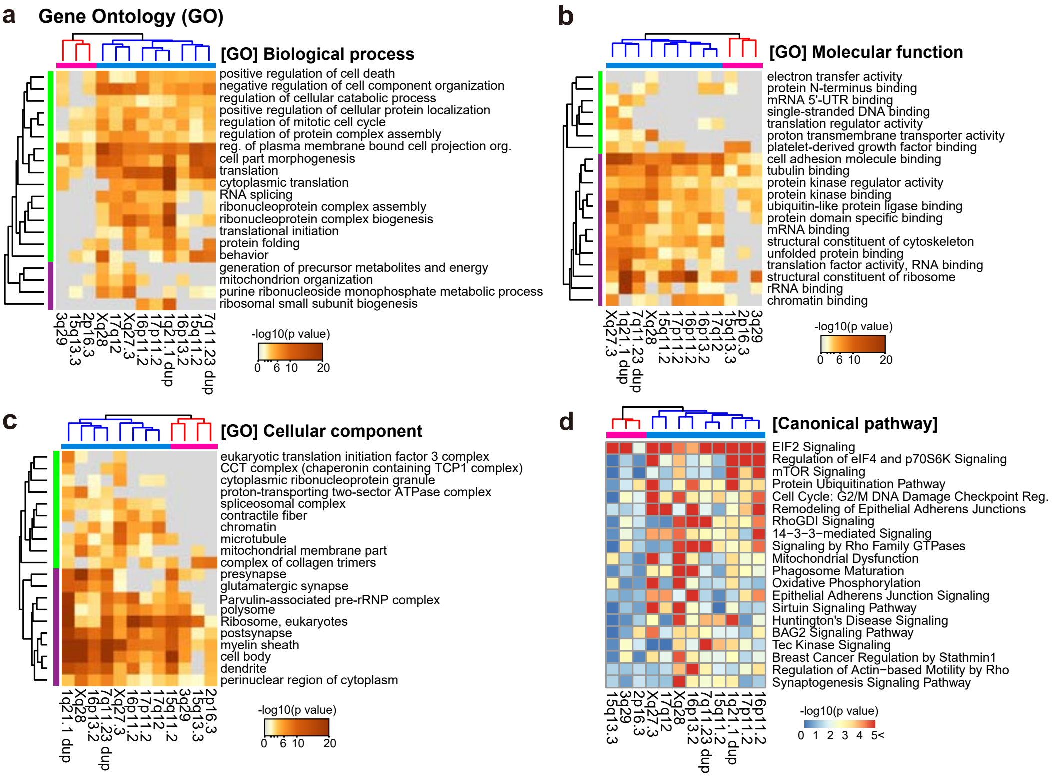

### Extended Data Fig. 5

##### Extended Data Figure 5.

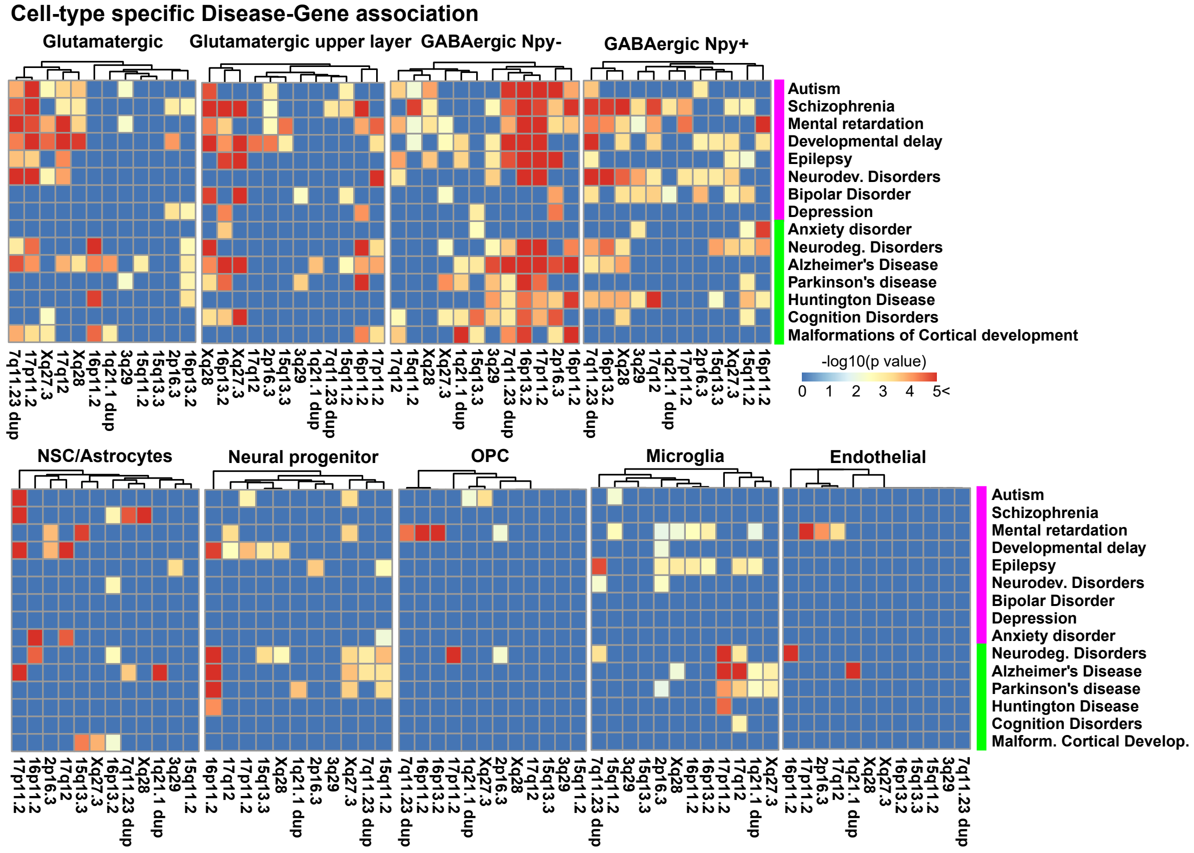

### Extended Data Fig. 6

##### Extended Data Figure 6.

###### a Glutamatergic postsynaptic density genes

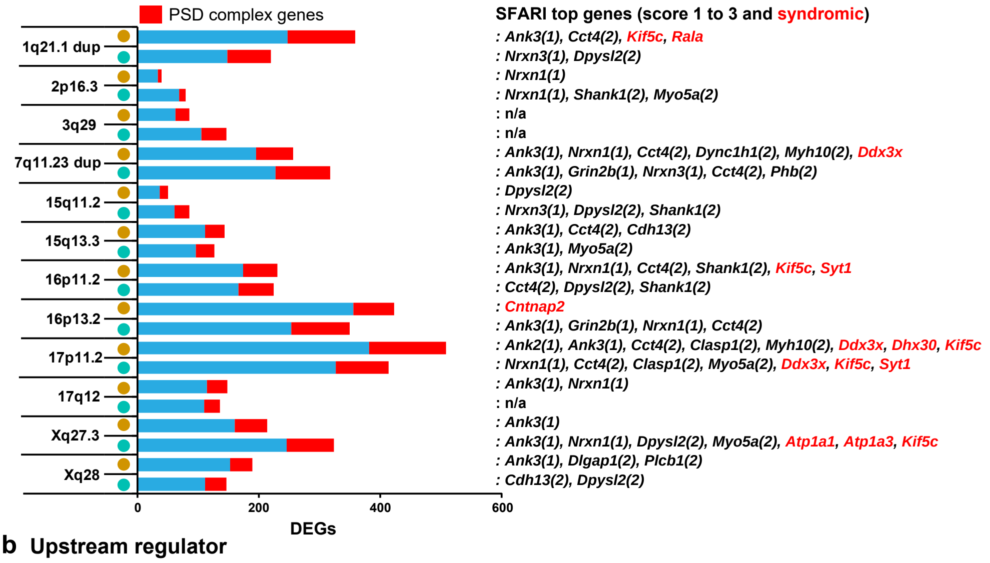


###### b Upstream regulator

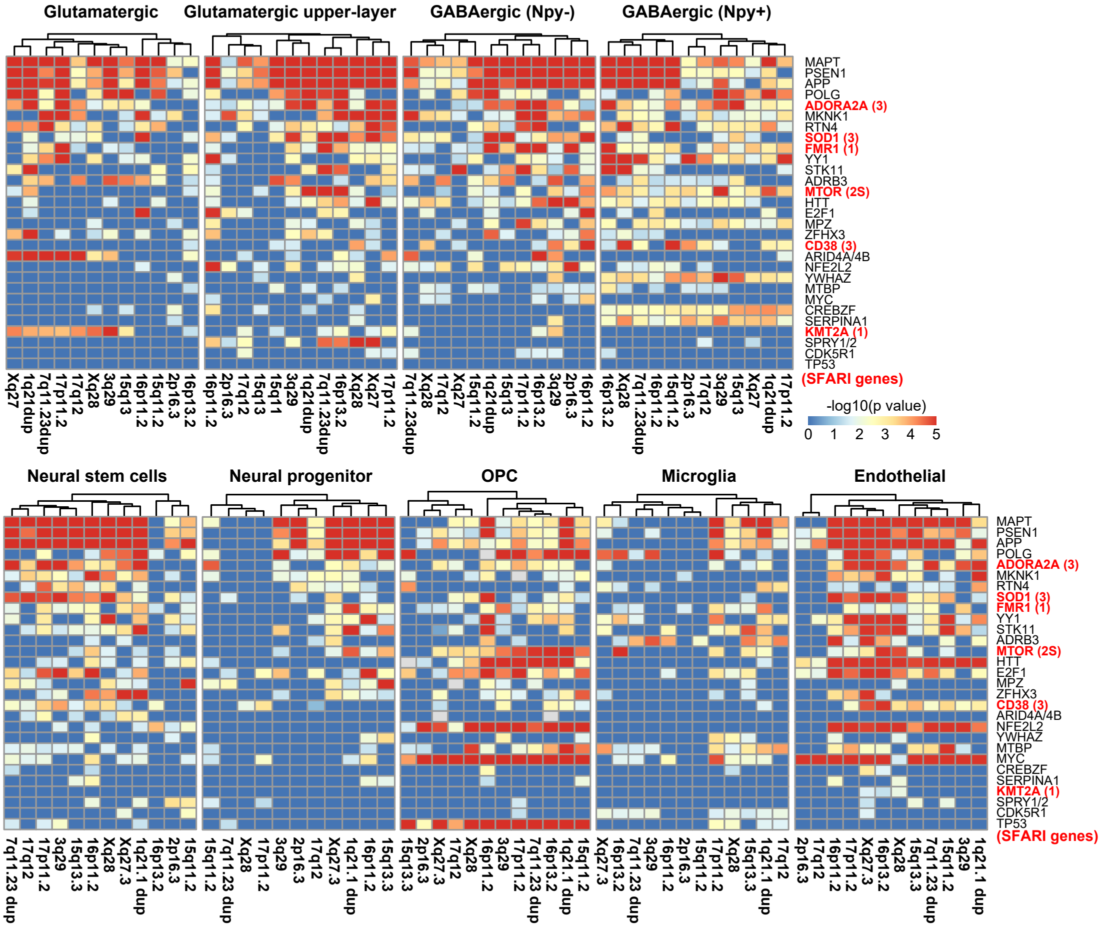
