## Extended Data Table 2 for "Autism in a dish: ES cell models of autism with copy number variations reveal cell-type-specific vulnerability"

**Extended Data Table2. Common CNVs among psychiatric disorders (targeted CNVs in our study).**

| Human chromosome | CNV type | Direction | p-value | OR (95% CI) | Associated disorders | Reference |
|---|---|---|---|---|---|---|
| 1q21.1 | Loss + gain | Risk | 1.50 × 10-6 | 3.8 (2.1–6.9) | **SCZ** | Marshall et al., 2017 |
| | gain | Risk | 2.20 x 10-2 | 2.64 (1.19–5.88) | **BPD** | Green et al., 2016 |
| | gain | Risk | 9.08× 10-4 | 2.17 (1.34-3.36) | **DP** | Kendall et al., 2019 |
| 2p16.3 (NRXN1) | Loss | Risk | 4.92 × 10-9 | 14.4 (4.2–46.9) | **SCZ** | Marshall et al., 2017 |
| | Loss | Risk | 5.70 x 10-3 | 2.01 (1.18-3.19) | **DP** | Kendall et al., 2019 |
| 3q29 | Loss | Risk | 1.86 × 10-6 | ∞ | **SCZ** | Marshall et al., 2017 |
| | Loss | Risk | 3.00 x 10-2 | 17.31 (1.57–190.97) | **BPD** | Green et al., 2016 |
| | Loss | Risk | 1.00 x 10-3 | 11.22 (2.27-46.52) | **DP** | Kendall et al., 2019 |
| 7q11.23 | Gain | Risk | 1.68 × 10-4 | 16.1 (3.1–125.7) | **SCZ** | Marshall et al., 2017 |
| 8p23.1 | Loss | Risk | 9.00 x 10-3 | 9.64 (1.32-50.21) | **DP** | Kendall et al., 2019 |
| 15q11.2 | Loss | Risk | 1.34 × 10-3 | 1.8 (1.2–2.6) | **SCZ** | Marshall et al., 2017 |
| Prader Willi syn | Gain | Risk | 4.61x10-5 | 8.14 (2.77-21.69) | **DP** | Kendall et al., 2019 |
| 15q13.3 | Loss | Risk | 2.13 × 10-7 | 15.6 (3.7–66.5) | **SCZ** | Marshall et al., 2017 |
| 16p11.2, proximal | Gain | Risk | 2.52 × 10-12 | 9.4 (4.2–20.9) | **SCZ** | Marshall et al., 2017 |
| | Gain | Risk | 2.3 × 10-4 | 4.37 (2.12–9.00) | **BPD** | Green et al., 2016 |
| | Gain | Risk | 2.04x10-4 | 2.65 (1.53-4.31) | **DP** | Kendall et al., 2019 |
| 16p11.2, distal | Loss | Risk | 5.52 × 10-5 | 20.6 (2.6–162.2) | **SCZ** | Marshall et al., 2017 |
| | Loss | Risk | 5.00 x 10-2 | 2.33 (0.92-4.63) | **DP** | Kendall et al., 2019 |
| 16p13.11 | Loss | Risk | 3.00 x 10-3 | 2.21 (1.25-3.63) | **DP** | Kendall et al., 2019 |
| 22q11.21 | Loss | Risk | 5.70 × 10-18 | 67.7 (9.3–492.8) | **SCZ** | Marshall et al., 2017 |

Marshall et al., Nat. Genetics., 49(1), 2017
Green et al., Mol. Psychiatry., 21(1), 2016
Kendall et al., JAMA Psychiatry., 2019
