## Extended Data Table 3 for "Autism in a dish: ES cell models of autism with copy number variations reveal cell-type-specific vulnerability"

**Extended Data Table 3. Common genes among psychiatric disorders (targeted CNVs in our study)**

| Gene_symbol | Human Chromosome | gene score | Syndromic | Genetic category | Associated disorders | Associated syndromes |
|---|---|---|---|---|---|---|
| Nrxn1 | 2p16.3 | 2 | - | Rare Single Gene Mutation | ADHD, EP, EPS, ASD, BPD, SCZ, ID | Pitt-Hopkins-like syndrome 2, Tourette syndrome |
| Birc6 | 2p22.3 | 4 | - | Rare Single Gene Mutation | - | - |
| Myt1l | 2p25.3 | 1 | - | Rare Single Gene Mutation | ASD | - |
| Pxdn | 2p25.3 | 4 | - | Rare Single Gene Mutation | - | - |
| Sntg2 | 2p25.3 | 4 | - | Rare Single Gene Mutation | - | - |
| Tpo | 2p25.3 | 4 | - | Genetic Association | - | - |
| Robo2 | 3p12.3 | 3 | - | Rare Single Gene Mutation | DD/NDD | - |
| Robo1 | 3p12.3 | 5 | - | Rare Single Gene Mutation | - | - |
| Foxp1 | 3p14.1 | 2 | S | Rare Single Gene Mutation | ASD, DD/NDD, ID | - |
| Fhit | 3p14.2 | 4 | - | Rare Single Gene Mutation | - | - |
| Cntn4 | 3p26.3 | 2 | - | Rare Single Gene Mutation | DD/NDD, ID | - |
| Cntn6 | 3p26.3 | 3 | - | Rare Single Gene Mutation | ADHD | Tourette syndrome |
| Pak2 | 3q29 | 3 | - | Rare Single Gene Mutation | - | 3q29 microdeletion syndrome |
| Dlg1 | 3q29 | 4 | - | Rare Single Gene Mutation | - | - |
| Tm4sf19 | 3q29 | 4 | - | Rare Single Gene Mutation | - | - |
| Pcdh10 | 4q28.3 | 4 | - | Rare Single Gene Mutation | - | - |
| Dusp22 | 6p25.3 | 6 | - | Rare Single Gene Mutation | ID | - |
| Auts2 | 7q11.22 | 3 | - | Rare Single Gene Mutation | ASD, DD/NDD, ID | - |
| Stx1a | 7q11.23 | 4 | - | Rare Single Gene Mutation | - | - |
| Gtf2i | 7q11.23 | 4 | - | Genetic Association | - | Williams syndrome |
| Stx1a | 7q11.23 | 4 | - | Rare Single Gene Mutation | - | - |
| Dock4 | 7q31.1 | 4 | - | Genetic Association | ASD | - |
| Immp2l | 7q31.1 | 4 | - | Rare Single Gene Mutation | ADHD, ASD, DD/NDD, ID | - |
| Pinx1 | 8p23.1 | 5 | - | Rare Single Gene Mutation | - | - |
| Ctnna3 | 10q21.3 | 4 | - | Genetic Association | ID, ADHD | - |
| Elp4 | 11p13 | 3 | - | Rare Single Gene Mutation | - | - |
| Pax6 | 11p13 | - | S | Syndromic | ASD, DD/NDD, ID | WAGR syndrome |
| Cacna1c | 12p13.33 | - | S | Syndromic | EPS, ASD, BPD, ID | Timothy syndrome |
| Prkd1 | 14q12 | - | S | Syndromic | EPS, DD/NDD | Rett syndrome |
| Pacs2 | 14q32.33 | - | S | Syndromic | ASD | - |
| Cyfip1 | 15q11.2 | 3 | - | - | ID | - |
| Nipa1 | 15q11.2 | 4 | - | - | ID | - |
| Nipa2 | 15q11.2 | 4 | - | - | ID | - |
| Tubgcp5 | 15q11.2 | 4 | - | Rare Single Gene Mutation | ID | - |
| Gabrb3 | 15q11.2-q13.1 | 2 | - | Rare Single Gene Mutation | ADHD, ASD, DD/NDD, ID | - |
| Magel2 | 15q11.2-q13.1 | 2 | S | Rare Single Gene Mutation | EPS, ASD, DD/NDD, ID | Schaaf-Yang syndrome, Chitayat-Hall syndrome |
| Atp10a | 15q11.2-q13.1 | 3 | - | Rare Single Gene Mutation | - | - |
| Gabrg3 | 15q11.2-q13.1 | 3 | - | Rare Single Gene Mutation | - | - |
| Ube3a | 15q11.2-q13.1 | 3 | S | Rare Single Gene Mutation | ID, EPS, ASD, DD/NDD, ADHD | Angelman syndrome |
| Gabra5 | 15q11.2-q13.1 | 5 | - | Functional | ASD, DD/NDD | - |
| Snrpn | 15q11.2-q13.1 | 5 | - | - | - | - |
| Herc2 | 15q11.2-q13.1 | - | S | Syndromic | EPS, ASD, DD/NDD | - |
| Apba2 | 15q13.1 | 4 | - | Rare Single Gene Mutation | - | - |
| Trpm1 | 15q13.2-q13.3 | 3 | - | Rare Single Gene Mutation | - | - |
| Fan1 | 15q13.2-q13.3 | 4 | - | Rare Single Gene Mutation | SCZ | - |
| Chrna7 | 15q13.3 | 3 | - | - | EP, EPS, ASD, DD/NDD, BPD, ID | - |
| Otud7a | 15q13.3 | 3 | - | Rare Single Gene Mutation | - | - |
| Taok2 | 16p11.2 | 2 | - | Rare Single Gene Mutation | - | - |
| Kctd13 | 16p11.2 | 4 | - | Rare Single Gene Mutation | - | - |
| Mapk3 | 16p11.2 | 4 | - | Rare Single Gene Mutation | - | - |
| Sez6l2 | 16p11.2 | 4 | - | - | - | - |
| Usp7 | 16p13.2 | 2 | S | Rare Single Gene Mutation | EPS, ASD, ADHD | - |
| Rbfox1 | 16p13.2 | 3 | - | Rare Single Gene Mutation | ASD, DD/NDD, ID | - |
| Abat | 16p13.2 | 4 | - | Genetic Association | - | - |
| Cntnap4 | 16q23.1 | 3 | - | Rare Single Gene Mutation | SCZ, ADHD | - |
| Adamts18 | 16q23.1 | 5 | - | Rare Single Gene Mutation | ASD | - |
| Chst5 | 16q23.1 | 6 | - | Rare Single Gene Mutation | - | - |
| Tmem231 | 16q23.1 | - | S | Syndromic | DD/NDD | Joubert syndrome-20, Meckel-Gruber syndrome |
| Cdh13 | 16q23.3 | 3 | - | Rare Single Gene Mutation | - | - |
| Rai1 | 17p11.2 | 3 | S | Rare Single Gene Mutation | EPS, ID | Smith-Magenis syndrome, Potocki-Lupski syndrome |
| Rasd1 | 17p11.2 | 5 | - | Functional | ASD | - |
| Myh4 | 17p13.1 | 4 | - | Rare Single Gene Mutation | - | Tourette syndrome |
| Ggnbp2 | 17q12 | 3 | - | Rare Single Gene Mutation | - | - |
| Macrod2 | 20p12.1 | 2 | - | Genetic Association | ADHD | - |
| Kcnq2 | 20q13.33 | 3 | - | Rare Single Gene Mutation | ADHD, ASD, DD/NDD, ID | - |
| Eef1a2 | 20q13.33 | - | S | Syndromic | ASD | - |
| Prodh | 22q11.21 | 3 | S | Rare Single Gene Mutation | EPS, ASD, EP | - |
| Gnb1l | 22q11.21 | 4 | - | Rare Single Gene Mutation | - | - |
| Tbx1 | 22q11.21 | 4 | - | Syndromic | ASD | DiGeorge syndrome |
| Shank3 | 22q13.33 | 1 | S | Rare Single Gene Mutation | EPS, ASD, DD/NDD, ID | Phelan-McDermid syndrome, Rett syndrome-like phenotype |
| Sbf1 | 22q13.33 | 3 | - | Rare Single Gene Mutation | - | Charcot-Marie-Tooth syndrome |
| Mapk8ip2 | 22q13.33 | 5 | - | Functional | - | - |

| | | | | | | |
|---|---|---|---|---|---|---|
| Chkb | 22q13.33 | - | S | Syndromic | ASD, DD/NDD, ID | - |
| Mapk12 | 22q13.33 | - | - | Rare Single Gene Mutation | ASD | - |
| Fmr1 | Xq27.3 | - | S | Syndromic | ID, EPS, ASD, DD/NDD, ADHD | Fragile X syndrome, Fragile X-associated tremor/ataxia syndrome |
| Mecp2 | Xq28 | 2 | S | Rare Single Gene Mutation | ID, EP, EPS, ASD, DD/NDD, SCZ, ADHD | Rett syndrome, X-linked intellectual disability, MECP2 duplication syndrome |

*ADHD, attention deficit with hyperactivity disorder; EPS, extrapyramidal symptoms; ID, intellectual disability; ASD, autism spectrum disorder; EP, epilepsy; SCZ, schizophrenia; DD/NDD, developmental delay/neurodevelopmental disorder; BPD, bipolar disorder.
