## Extended Data Table 4 for "Autism in a dish: ES cell models of autism with copy number variations reveal cell-type-specific vulnerability"

**Extended Data Table 4. Targeted CNVs in mouse ES cells**

| | | | | | 5' end gene | | | | 3' end gene | | | |
|---|---|---|---|---|---|---|---|---|---|---|---|---|
| **Human chr** | **Human CNV locus** | **Mouse chr** | **Length (Kb)** | **CNV type** | **Gene** | **Chr_start** | **Chr_end** | **strand** | **Gene** | **Chr_start** | **Chr_end** | **strand** |
| 1 | 1q21.1 | 3 | 805.5 | del/+ | Gpr89 | 96868281 | 96905346 | - | Prkab2 | 97658193 | 97673812 | + |
| 1 | 1q21.1 | 3 | 805.5 | dup/+ | Gpr89 | 96868281 | 96905346 | - | Prkab2 | 97658193 | 97673812 | + |
| 1 | 1q25.3 | 1 | 1398.7 | del/+ | Swt1 | 151367699 | 151428455 | - | Rgl1 | 152516760 | 152766351 | - |
| 2 | 2p16.3 | 17 | 1059.4 | del/+ | Nrxn1 | 90033631 | 91093071 | - | - | - | - | N/A |
| 2 | 2p21 | 17 | 673.3 | del/+ | Srbd1 | 85984665 | 86145175 | - | Prkce | 86167785 | 86657919 | + |
| 2 | 2p22.3 | 17 | 864.2 | del/+ | Birc6 | 74528295 | 74703356 | + | Ltbp1 | 75005568 | 75392512 | + |
| 2 | 2p25.3 | 12 | 1431.8 | del/+ | Myt1l | 29528384 | 29923213 | + | Sh3yl1 | 30911668 | 30960162 | + |
| 3 | 3p12.3 | 16 | 1748.7 | del/+ | Robo1 | 72663149 | 73046095 | + | Robo2 | 73892306 | 74411825 | - |
| 3 | 3p14.1 | 6 | 3409.6 | del/+ | Fam19a1 | 96113154 | 96657198 | + | Foxp1 | 98925338 | 99522721 | - |
| 3 | 3p14.2 | 14 | 1611.9 | del/+ | Fhit | 9550092 | 11162035 | - | - | - | - | N/A |
| 3 | 3p26.3 | 6 | 3289.5 | del/+ | Chl1 | 103510586 | 103750211 | + | Crbn | 106780201 | 106800077 | - |
| 3 | 3q29 | 16 | 1210.5 | del/+ | Bdh1 | 31422280 | 31458901 | + | Tfrc | 32608920 | 32632794 | + |
| 4 | 4p16.3 | 5 | 462.3 | del/+ | Pigg | 108312609 | 108349355 | + | Rnf212 | 108729308 | 108774953 | - |
| 4 | 4p16.3 | 5 | 336.1 | del/+ | Fam53a | 33600347 | 33629635 | - | Nelfa | 33897916 | 33936413 | - |
| 4 | 4q13.2 | 5 | 601.1 | del/+ | Ugt2b34 | 86889767 | 86906937 | - | Ugt2a1 | 87459490 | 87490871 | - |
| 4 | 4q28.3 | 3 | 1069.8 | del/+ | Pcdh10 | 45378398 | 45435623 | + | Pabpc4l | 46442197 | 46448219 | - |
| 5 | 5p15.33 | 13 | 124.4 | del/+ | Brd9 | 73937838 | 73960894 | + | Cep72 | 74036495 | 74062285 | - |
| 6 | 6p12.3 | 17 | 903.3 | dup/+ | Adgrf2 | 42708936 | 42742179 | - | Pla2g7 | 43568098 | 43612201 | + |
| 6 | 6p25.3 | 13 | 314.0 | del/+ | Dusp22 | 30659999 | 30711231 | + | Exoc2 | 30813919 | 30974047 | - |
| 6 | 6q26 | 17 | 1223.0 | del/+ | Park2 | 10840384 | 12063361 | + | - | - | - | N/A |
| 6 | 6q27 | 17 | 721.3 | del/+ | Tcte2 | 13482553 | 13761825 | - | Dact2 | 14195231 | 14203831 | - |
| 7 | 7p21.1 | 12 | 956.9 | del/+ | Snx13 | 35047189 | 35147469 | + | Agr2 | 35992907 | 36004087 | + |
| 7 | 7q11.22 | 5 | 1106.0 | del/+ | Auts2 | 131437333 | 132543344 | - | - | - | - | N/A |
| 7 | 7q11.23 | 5 | 731.8 | del/+ | Gm1620 | 134636240 | 134463449 | + | Trim50 | 135353295 | 135368005 | + |
| 7 | 7q11.23 | 5 | 731.8 | dup/+ | Gm1620 | 134636240 | 134463449 | + | Trim50 | 135353295 | 135368005 | + |
| 7 | 7q11.23 | 5 | 1130.2 | del/+ | Gtf2i | 134237834 | 134314760 | - | Trim50 | 135353295 | 135368005 | + |
| 7 | 7q31.1 | 12 | 1509.5 | del/+ | Dock4 | 40446053 | 40846486 | + | Immp2l | 41024090 | 41955588 | + |
| 7 | 7q35 | 6 | 1081.4 | del/+ | Tcaf3 | 42584866 | 42597692 | - | Tpk1 | 43345001 | 43666278 | - |
| 8 | 8p22 | 8 | 2093.8 | del/+ | Dlc1 | 36567751 | 36953143 | - | Sgcz | 37522298 | 38661508 | - |
| 8 | 8p23.1 | 8 | 1318.6 | del/+ | Tnks | 34829179 | 34965690 | - | D8Ertd82e | 36094828 | 36147787 | + |
| 8 | 8p23.1 | 14 | 1468.8 | del/+ | Ctsb | 63122462 | 63145919 | + | Mir3078 | 64591185 | 64591271 | + |
| 10 | 10q21.3 | 10 | 1573.6 | del/+ | Ctnna3 | 63430098 | 65003667 | + | - | - | - | N/A |
| 11 | 11p13 | 2 | 1577.1 | del/+ | Hipk3 | 104426481 | 104494446 | - | Dnajc24 | 105966709 | 106003549 | - |
| 12 | 12p13.33 | 6 | 609.2 | del/+ | Cacna1c | 118587240 | 119196418 | - | - | - | - | N/A |
| 14 | 14q32.33 | 12 | 1696.9 | del/+ | Ppp1r13b | 111828457 | 111908110 | - | Ighd3 | 113525313 | 113525329 | - |
| 15 | 15q11.2 | 7 | 225.8 | del/+ | Tubgcp5 | 55794148 | 55831447 | + | Nipa1 | 55977567 | 56019954 | - |
| 15 | 15q11.2 | 14 | 39.6 | del/+ | Olfr732 | 50281228 | 50282271 | - | Olfr734 | 50319836 | 50320868 | - |
| 15 | 15q11.2-q13.1 | 7 | 6369.9 | del/+ | Herc2 | 56050196 | 56231800 | + | Mkrn3 | 62417593 | 62420139 | - |
| 15* | 15q11.2-q13.1 (paternal) | 7 | 6369.9 | dup/+ | Herc2 | 56050196 | 56231800 | + | Mkrn3 | 62417593 | 62420139 | - |
| 15* | 15q11.2-q13.1 (maternal) | 7 | 6369.9 | dup/+ | Herc2 | 56050196 | 56231800 | + | Mkrn3 | 62417593 | 62420139 | - |
| 15 | 15q13.1-q13.2 | 7 | 1026.1 | del/+ | Apba2 | 64501706 | 64753878 | + | Tjp1 | 65296165 | 65527781 | - |
| 15 | 15q13.2-q13.3 | 7 | 1293.5 | del/+ | Chrna7 | 63098692 | 63212526 | - | Mphosph10 | 64376541 | 64392236 | - |
| 15 | 15q13.3 | 7 | 660.3 | del/+ | Chrna7 | 63098692 | 63212526 | - | Otud7a | 63444751 | 63759028 | + |
| 15 | 15q14 | 7 | 2081.7 | del/+ | Tarsl2 | 65644898 | 65692091 | + | Ttc23 | 67647410 | 67726576 | + |
| 15 | 15q25.2-25.3 | 7 | 473.6 | del/+ | Zscan2 | 80860920 | 80876516 | + | Pde8a | 81213596 | 81334533 | + |
| 16 | 16p11.2 | 7 | 438.1 | del/+ | Coro1a | 126699773 | 126707787 | - | Spn | 127132232 | 127137823 | - |
| 16 | 16p11.2 | 7 | 438.1 | dup/- | Coro1a | 126699773 | 126707787 | - | Spn | 127132232 | 127137823 | - |
| 16 | 16p12.1 | 7 | 347.1 | del/+ | Uqcrc2 | 120635176 | 120659478 | + | Cdr2 | 120957036 | 120982312 | - |
| 16 | 16p13.11 | 16 | 234.6 | del/+ | Pla2g10 | 13715057 | 13730983 | - | Mpv17l | 13903161 | 13949619 | + |
| 16 | 16p13.11 | 16 | 572.6 | del/+ | Mpv17l | 13903161 | 13949619 | + | Abcc1 | 14361558 | 14475737 | + |
| 16 | 16p13.2 | 16 | 603.3 | del/+ | Rbfox1 | 6809222 | 7412479 | + | - | - | - | N/A |
| 16 | 16p13.2 | 16 | 603.3 | dup/- | Rbfox1 | 6809222 | 7412479 | + | - | - | - | N/A |
| 16 | 16p13.2 | 16 | 513.1 | del/+ | Tmem114 | 8409276 | 8425136 | - | Gm10247 | 8922057 | 8922389 | - |
| 16 | 16q23.1 | 8 | 3903.9 | del/+ | Wdr59 | 111448797 | 111522092 | - | Wwox | 114439655 | 115352708 | + |
| 16 | 16q23.3 | 8 | 1767.1 | del/+ | Mphosph6 | 117791645 | 117801929 | - | Mbtps1 | 119508156 | 119558761 | - |
| 17 | 17p11.2 | 11 | 1127.0 | del/+ | Mprip | 59661305 | 59780860 | + | Smcr8 | 60777524 | 60788287 | + |
| 17 | 17p12 | 11 | 971.1 | del/+ | Cdrt4 | 62951193 | 62993095 | + | Hs3st3b1 | 63885792 | 63922290 | - |
| 17 | 17p13.1 | 11 | 777.0 | del/+ | Pirt | 66911981 | 66929876 | + | Gas7 | 67455437 | 67688990 | + |
| 17 | 17q12 | 11 | 1066.3 | del/+ | Hnf1b | 83850063 | 83905819 | + | Znhit3 | 84910950 | 84916366 | - |

| | | | | | | | | | | | | |
|---|---|---|---|---|---|---|---|---|---|---|---|---|
| 20 | 20p12.1 | 2 | 1997.7 | del/+ | Macrod2 | 140395309 | 142392966 | + | - | - | - | N/A |
| 20 | 20q13.33 | 2 | 1268.2 | del/+ | Ogfr | 180589245 | 180595836 | + | Pcmtd2 | 181837854 | 181857461 | + |
| 22 | 22q11.21 | 16 | 1408.4 | del/+ | P2rx6 | 17561885 | 17577800 | + | Hira | 18877037 | 18970309 | + |
| 22 | 22q13.33 | 15 | 701.3 | del/+ | Ttll8 | 88890633 | 88954418 | - | Rabl2 | 89582533 | 89591923 | - |
| x | Xq27.3 | X | 39.4 | del/y | Fmr1 | 68678541 | 68717963 | + | - | - | - | N/A |
| x | Xq28 | X | 108.8 | del/y | Mecp2 | 74026592 | 74135363 | - | - | - | - | N/A |

*15q11.2-q13.1 paternal duplication (patdp/+) cells were established from mouse blastocyst.
