## Extended Data Table 5 for "Autism in a dish: ES cell models of autism with copy number variations reveal cell-type-specific vulnerability"

**Extended Data Table 5. Chromosome Targeting**

| Chr | Cytoband | cnv-type | deletion-values | duplication-values | Number-case-individuals | Number-of-reports | Chrmosome targeting | Conservation of genes between human and mice |
|---|---|---|---|---|---|---|---|---|
| 16 | 16p11.2 | Deletion-Duplication | 765 | 760 | 1525 | 104 | Succeeded | High |
| 15 | 15q11.2 | Deletion-Duplication | 595 | 1559 | 2154 | 87 | Succeeded | High |
| 2 | 2p16.3 | Deletion-Duplication | 459 | 22 | 481 | 62 | Succeeded | High |
| 1 | 1q21.1 | Deletion-Duplication | 422 | 499 | 921 | 52 | Succeeded | High |
| 7 | 7q11.23 | Deletion-Duplication | 197 | 187 | 384 | 56 | Succeeded | High |
| 3 | 3q29 | Deletion-Duplication | 248 | 183 | 431 | 53 | Succeeded | High |
| 22 | 22q11.21 | Deletion-Duplication | 533 | 444 | 977 | 79 | Succeeded | High |
| X | Xq28 | Deletion-Duplication | 137 | 277 | 414 | 58 | Succeeded | High |
| 15 | 15q13.3 | Deletion-Duplication | 90 | 323 | 413 | 66 | Succeeded | High |
| 17 | 17q12 | Deletion-Duplication | 241 | 232 | 473 | 59 | Succeeded | High |
| 1 | 1q44 | Deletion-Duplication | 468 | 111 | 579 | 47 | - | Low |
| 3 | 3p26.3 | Deletion-Duplication | 193 | 195 | 388 | 48 | Succeeded | High |
| 17 | 17p11.2 | Deletion-Duplication | 168 | 136 | 304 | 45 | Succeeded | High |
| 22 | 22q13.33 | Deletion-Duplication | 184 | 31 | 215 | 54 | Succeeded | High |
| X | Xp22.31 | Deletion-Duplication | 92 | 195 | 287 | 40 | - | Low |
| 2 | 2q37.3 | Deletion-Duplication | 137 | 97 | 234 | 43 | - | Low |
| 8 | 8p23.1 | Deletion-Duplication | 221 | 190 | 411 | 41 | Succeeded | High |
| 14 | 14q11.2 | Deletion-Duplication | 231 | 133 | 364 | 45 | - | Low |
| 17 | 17q21.31 | Duplication | 124 | 57 | 181 | 39 | Failed | - |
| 17 | 17p13.3 | Deletion-Duplication | 195 | 229 | 424 | 45 | - | Low |
| 4 | 4q35.2 | Deletion-Duplication | 94 | 139 | 233 | 43 | Failed | - |
| 4 | 4q13.2 | Deletion-Duplication | 301 | 37 | 338 | 30 | Succeeded | High |
| 5 | 5q35.3 | Deletion-Duplication | 54 | 174 | 228 | 37 | Failed | - |
| 11 | 11p15.4 | Deletion-Duplication | 136 | 133 | 269 | 33 | - | Low |
| 12 | 12p13.31 | Deletion-Duplication | 163 | 104 | 267 | 27 | - | Low |
| 14 | 14q32.33 | Deletion-Duplication | 79 | 83 | 162 | 30 | Succeeded | High |
| 16 | 16p13.3 | Deletion-Duplication | 284 | 188 | 472 | 46 | - | Low |
| 7 | 7q11.22 | Deletion-Duplication | 119 | 82 | 201 | 34 | Succeeded | High |
| 8 | 8p22 | Deletion-Duplication | 500 | 70 | 570 | 36 | Succeeded | High |
| 16 | 16p13.2 | Deletion-Duplication | 129 | 98 | 227 | 35 | Succeeded | High |
| 2 | 2p22.3 | Deletion-Duplication | 128 | 99 | 227 | 32 | Succeeded | High |
| 2 | 2p25.3 | Deletion-Duplication | 49 | 66 | 115 | 33 | Succeeded | High |
| 2 | 2q13 | Deletion-Duplication | 186 | 136 | 322 | 46 | Failed | - |
| 6 | 6q26 | Deletion-Duplication | 210 | 109 | 319 | 36 | Succeeded | High |
| 6 | 6q27 | Deletion-Duplication | 97 | 194 | 291 | 28 | Succeeded | High |
| 7 | 7q35 | Deletion-Duplication | 70 | 54 | 124 | 38 | Succeeded | High |
| 8 | 8p23.2 | Deletion-Duplication | 188 | 119 | 307 | 32 | Failed | - |
| 9 | 9q34.3 | Deletion-Duplication | 135 | 171 | 306 | 37 | Failed | - |
| 10 | 10q26.3 | Deletion-Duplication | 54 | 249 | 303 | 37 | Failed | - |
| 15 | 15q11.2-q13.1 | Duplication | 117 | 244 | 361 | 55 | Succeeded | High |
| 16 | 16p13.11 | Deletion-Duplication | 207 | 449 | 656 | 50 | Succeeded | High |
| 22 | 22q11.23 | Deletion-Duplication | 51 | 80 | 131 | 29 | Failed | - |
| X | Xp22.33 | Deletion-Duplication | 56 | 156 | 212 | 37 | - | Low |
| 3 | 3q26.1 | Deletion-Duplication | 200 | 44 | 244 | 24 | - | Low |
| 4 | 4p16.3 | Deletion-Duplication | 127 | 168 | 295 | 35 | Succeeded | High |
| 5 | 5p15.2 | Deletion-Duplication | 220 | 35 | 255 | 28 | - | Low |
| 5 | 5p15.33 | Deletion-Duplication | 82 | 517 | 599 | 33 | Succeeded | High |
| 7 | 7p21.1 | Deletion-Duplication | 97 | 58 | 155 | 26 | Succeeded | High |
| 7 | 7p21.3 | Deletion-Duplication | 239 | 73 | 312 | 27 | Failed | - |
| 10 | 10q21.3 | Deletion-Duplication | 317 | 27 | 344 | 36 | Succeeded | High |
| 12 | 12p13.33 | Deletion-Duplication | 160 | 39 | 199 | 31 | Succeeded | High |
| 15 | 15q13.2-q13.3 | Deletion-Duplication | 223 | 77 | 300 | 39 | Succeeded | High |
| 16 | 16q23.1 | Deletion-Duplication | 163 | 97 | 260 | 37 | Succeeded | High |
| 20 | 20q13.33 | Deletion-Duplication | 93 | 80 | 173 | 30 | Succeeded | High |
| 2 | 2p21 | Deletion-Duplication | 192 | 71 | 263 | 27 | Succeeded | High |
| 5 | 5q23.1 | Deletion-Duplication | 132 | 48 | 180 | 27 | Failed | - |
| 6 | 6p25.3 | Deletion-Duplication | 76 | 102 | 178 | 18 | Succeeded | High |
| 9 | 9p23 | Deletion-Duplication | 363 | 86 | 449 | 27 | - | Low |
| 17 | 17p13.1 | Deletion-Duplication | 83 | 69 | 152 | 26 | Succeeded | High |
| 20 | 20p13 | Deletion-Duplication | 50 | 48 | 98 | 30 | Failed | - |
| 22 | 22q13.1 | Deletion-Duplication | 56 | 39 | 95 | 24 | - | Low |
| 2 | 2p11.2 | Deletion-Duplication | 82 | 120 | 202 | 25 | - | Low |
| 4 | 4q28.3 | Deletion-Duplication | 122 | 42 | 164 | 19 | Succeeded | High |
| 6 | 6p21.32 | Deletion-Duplication | 165 | 101 | 266 | 23 | Failed | - |
| 6 | 6q14.1 | Deletion-Duplication | 294 | 44 | 338 | 22 | - | Low |
| 7 | 7q31.1 | Deletion-Duplication | 218 | 55 | 273 | 34 | Succeeded | High |
| 11 | 11p13 | Deletion-Duplication | 61 | 31 | 92 | 25 | Succeeded | High |
| 12 | 12q12 | Deletion-Duplication | 66 | 33 | 99 | 20 | - | Low |
| 13 | 13q21.1 | Deletion-Duplication | 84 | 19 | 103 | 18 | - | Low |
| 14 | 14q12 | Deletion-Duplication | 59 | 37 | 96 | 20 | Failed | - |
| 15 | 15q14 | Deletion-Duplication | 59 | 27 | 86 | 22 | - | Low |
| 15 | 15q26.3 | Deletion-Duplication | 65 | 149 | 214 | 33 | Succeeded | High |
| 18 | 18p11.32 | Deletion-Duplication | 103 | 50 | 153 | 21 | - | Low |
| 20 | 20p12.1 | Deletion-Duplication | 246 | 33 | 279 | 34 | Succeeded | High |
| 22 | 22q12.3 | Deletion-Duplication | 44 | 51 | 95 | 22 | - | Low |
| 1 | 1p31.3 | Deletion-Duplication | 26 | 37 | 63 | 23 | - | Low |
| 2 | 2q33.1 | Deletion-Duplication | 46 | 29 | 75 | 19 | - | Low |
| 3 | 3p14.1 | Deletion-Duplication | 46 | 23 | 69 | 21 | Succeeded | High |
| 3 | 3p14.2 | Deletion-Duplication | 122 | 17 | 139 | 25 | Succeeded | High |
| 3 | 3p26.2 | Deletion-Duplication | 160 | 8 | 168 | 17 | - | Low |
| 4 | 4p15.1 | Deletion-Duplication | 100 | 25 | 125 | 18 | - | Low |
| 7 | 7q34 | Deletion-Duplication | 296 | 33 | 329 | 20 | - | Low |
| 8 | 8q24.3 | Deletion-Duplication | 69 | 78 | 147 | 35 | - | Low |
| 14 | 14q24.3 | Deletion-Duplication | 59 | 25 | 84 | 19 | - | Low |
| 16 | 16p12.1 | Deletion-Duplication | 194 | 43 | 237 | 20 | Succeeded | High |
| 16 | 16q23.3 | Deletion-Duplication | 39 | 31 | 70 | 20 | Succeeded | High |

| | | | | | | | | |
|---|---|---|---|---|---|---|---|---|
| 17 | 17p12 | Deletion-Duplication | 110 | 72 | 182 | 35 | Succeeded | High |
| 19 | 19q13.42 | Deletion-Duplication | 59 | 68 | 127 | 28 | Failed | - |
| X | Xp11.4 | Deletion-Duplication | 21 | 70 | 91 | 25 | - | Low |
| 1 | 1p21.1 | Deletion-Duplication | 64 | 51 | 115 | 23 | - | Low |
| 1 | 1p31.1 | Deletion-Duplication | 132 | 43 | 175 | 28 | - | Low |
| 1 | 1p36.33 | Deletion-Duplication | 101 | 120 | 221 | 24 | Failed | - |
| 1 | 1q24.2 | Deletion | 32 | 7 | 39 | 14 | - | Low |
| 1 | 1q25.3 | Deletion-Duplication | 28 | 50 | 78 | 18 | Succeeded | High |
| 1 | 1q31.1 | Deletion-Duplication | 104 | 20 | 124 | 17 | - | Low |
| 2 | 2q32.1 | Deletion-Duplication | 137 | 36 | 173 | 23 | - | Low |
| 3 | 3p12.3 | Deletion-Duplication | 50 | 51 | 101 | 21 | Succeeded | High |
| 3 | 3q23 | Deletion-Duplication | 17 | 60 | 77 | 18 | Failed | - |
| 4 | 4p16.1 | Deletion-Duplication | 45 | 48 | 93 | 24 | - | Low |
| 4 | 4q12 | Deletion-Duplication | 69 | 77 | 146 | 24 | - | Low |
| X | Xq27.3 | Deletion-Duplication | 83 | 30 | 113 | 12 | Succeeded | High |
| 15 | 15q25.2-q25.3 | Deletion | 14 | 1 | 15 | 8 | Succeeded | High |
| 15 | 15q13.1-q13.2 | Deletion-Duplication | 15 | 11 | 26 | 12 | Succeeded | High |
