## Extended Data Table 6 for "Autism in a dish: ES cell models of autism with copy number variations reveal cell-type-specific vulnerability"

**Extended Data Table 6. Synteny analysis; Targeted CNVs**

| Human Chromosome | Homo sapiens genes<br>Human (GRCh38.p12) | Human location | | Mus musculus homologues<br>Mouse (GRCm38.p6) | Mus musculus location |
|---|---|---|---|---|---|
| **1q21.1** | GPR89A (ENSG00000117262) | 1:145607988-145670650 | → | Gpr89 (ENSMUSG00000028096) | 3:96868281-96905346 |
| | PDZK1 (ENSG00000174827) | 1:145670852-145708148 | → | Pdzk1 (ENSMUSG00000038298) | 3:96829284-96870926 |
| | CD160 (ENSG00000117281) | 1:145719471-145739288 | → | Cd160 (ENSMUSG00000038304) | 3:96798763-96829351 |
| | RNF115 (ENSG00000265491) | 1:145738868-145824095 | → | Rnf115 (ENSMUSG00000028098) | 3:96727664-96791638 |
| | POLR3C (ENSG00000186141) | 1:145824088-145844402 | → | Polr3c (ENSMUSG00000028099) | 3:96711490-96727628 |
| | NUDT17 (ENSG00000186364) | 1:145845630-145848954 | → | Nudt17 (ENSMUSG00000028100) | 3:96706067-96708562 |
| | PIAS3 (ENSG00000131788) | 1:145848522-145859836 | → | Pias3 (ENSMUSG00000028101) | 3:96696384-96706070 |
| | ANKRD35 (ENSG00000198483) | 1:145866560-145885866 | → | Ankrd35 (ENSMUSG00000038354) | 3:96670131-96691032 |
| | ITGA10 (ENSG00000143127) | 1:145891208-145910111 | → | Itga10 (ENSMUSG00000090210) | 3:96645584-96664519 |
| | PEX11B (ENSG00000131779) | 1:145911350-145918717 | → | Pex11b (ENSMUSG00000028102) | 3:96635376-96645366 |
| | RBM8A (ENSG00000265241) | 1:145917714-145927678 | → | Rbm8a (ENSMUSG00000038374) | 3:96629933-96633791 |
| | | | → | Rbm8a2 (ENSMUSG00000078184) | 1:175977777-175979114 |
| | AC243547.3 (ENSG00000280778) | 1:145927258-145977811 | | No homologues | |
| | LIX1L (ENSG00000271601) | 1:145933423-145958017 | → | Lix1l (ENSMUSG00000049288) | 3:96601149-96626171 |
| | ANKRD34A (ENSG00000272031) | 1:145959441-145964575 | → | Ankrd34a (ENSMUSG00000049097) | 3:96596636-96599775 |
| | POLR3GL (ENSG00000121851) | 1:145964690-145978848 | | No homologues | |
| | TXNIP (ENSG00000265972) | 1:145992435-145996579 | → | Txnip (ENSMUSG00000038393) | 3:96557957-96561883 |
| | HJV (ENSG00000168509) | 1:146017468-146036746 | → | Hjv (ENSMUSG00000038403) | 3:96525172-96529210 |
| | NBPF10 (ENSG00000271425) | 1:146064711-146229000 | | No homologues | |
| | NOTCH2NLA (ENSG00000264343) | 1:146146203-146229026 | | No homologues | |
| | PPIAL4H (ENSG00000270339) | 1:146344131-146344790 | → | Ppia (ENSMUSG00000071866) | 11:6415443-6419817 |
| | NBPF12 (ENSG00000268043) | 1:146938744-146996202 | | No homologues | |
| | PRKAB2 (ENSG00000131791) | 1:147155106-147172550 | → | Prkab2 (ENSMUSG00000038205) | 3:97658193-97673812 |
| **2p16.3** | NRXN1 (ENSG00000179915) | 2:49918503-51225575 | → | Nrxn1 (ENSMUSG00000024109) | 17:90033631-91093071 |
| **3q29** | TFRC (ENSG00000072274) | 3:196027183-196082096 | → | Tfrc (ENSMUSG00000022797) | 16:32608920-32632794 |
| | ZDHHC19 (ENSG00000163958) | 3:196197452-196211437 | → | Zdhhc19 (ENSMUSG00000052363) | 16:32496265-32561966 |
| | SLC51A (ENSG00000163959) | 3:196211487-196243178 | → | Slc51a (ENSMUSG00000035699) | 16:32474504-32487879 |
| | PCYT1A (ENSG00000161217) | 3:196214222-196287957 | → | Pcyt1a (ENSMUSG00000005615) | 16:32430921-32475070 |
| | AC069257.3 (ENSG00000272741) | 3:196248230-196318222 | → | Tctex1d2 (ENSMUSG00000014075) | 16:32419702-32429099 |
| | TCTEX1D2 (ENSG00000213123) | 3:196291219-196318299 | → | Tctex1d2 (ENSMUSG00000014075) | 16:32419702-32429099 |
| | TM4SF19-TCTEX1D2 (ENSG00000273331) | 3:196316082-196338373 | → | Tm4sf19 (ENSMUSG00000079625) | 16:32400506-32408227 |
| | TM4SF19 (ENSG00000145107) | 3:196319342-196338503 | → | Tm4sf19 (ENSMUSG00000079625) | 16:32400506-32408227 |
| | UBXN7 (ENSG00000163960) | 3:196347662-196432430 | → | Ubxn7 (ENSMUSG00000053774) | 16:32332257-32393747 |
| | RNF168 (ENSG00000163961) | 3:196468783-196503768 | → | Rnf168 (ENSMUSG00000014074) | 16:32277459-32301434 |
| | SMCO1 (ENSG00000214097) | 3:196506879-196515346 | → | Smco1 (ENSMUSG00000046345) | 16:32271468-32274781 |
| | WDR53 (ENSG00000185798) | 3:196554177-196568674 | → | Wdr53 (ENSMUSG00000022787) | 16:32247227-32257083 |
| | FBXO45 (ENSG00000174013) | 3:196568611-196589059 | → | Fbxo45 (ENSMUSG00000035764) | 16:32230112-32247158 |
| | NRROS (ENSG00000174004) | 3:196639694-196662004 | → | Nrros (ENSMUSG00000052384) | 16:32142785-32165594 |
| | PIGX (ENSG00000163964) | 3:196639775-196736007 | → | Pigx (ENSMUSG00000023791) | 16:32084416-32099740 |
| | CEP19 (ENSG00000174007) | 3:196706277-196712250 | → | Cep19 (ENSMUSG00000035790) | 16:32099800-32108069 |
| | PAK2 (ENSG00000180370) | 3:196739857-196832647 | → | Pak2 (ENSMUSG00000022781) | 16:32016290-32079342 |
| | SENP5 (ENSG00000119231) | 3:196867856-196934714 | → | Senp5 (ENSMUSG00000022772) | 16:31959672-32003287 |
| | NCBP2 (ENSG00000114503) | 3:196935402-196942594 | → | Ncbp2 (ENSMUSG00000022774) | 16:31948513-31961781 |
| | NCBP2AS2 (ENSG00000270170) | 3:196942674-196943543 | → | 0610012G03Rik (ENSMUSG00000107002) | 16:31947050-31948494 |
| | PIGZ (ENSG00000119227) | 3:196946356-196969060 | | No homologues | |
| | MELTF (ENSG00000163975) | 3:196988621-197029817 | → | Meltf (ENSMUSG00000022780) | 16:31878810-31899020 |
| | DLG1 (ENSG00000075711) | 3:197042560-197299330 | → | Dlg1 (ENSMUSG00000022770) | 16:31663443-31875129 |
| | DLG1 (ENSG00000075711) | 3:197042560-197299330 | → | Dlg1 (ENSMUSG00000022770) | 16:31663443-31875129 |
| | BDH1 (ENSG00000161267) | 3:197509783-197573323 | → | Bdh1 (ENSMUSG00000046598) | 16:31422280-31458901 |
| **7q11.23** | TRIM50 (ENSG00000146755) | 7:73312536-73328082 | → | Trim50 (ENSMUSG00000053388) | 5:135353295-135368005 |
| | FKBP6 (ENSG00000077800) | 7:73328161-73358637 | → | Fkbp6 (ENSMUSG00000040013) | 5:135291704-135350044 |
| | FZD9 (ENSG00000188763) | 7:73433778-73436120 | → | Fzd9 (ENSMUSG00000049551) | 5:135248938-135251230 |
| | BAZ1B (ENSG00000009954) | 7:73440406-73522293 | → | Baz1b (ENSMUSG00000002748) | 5:135187264-135246129 |
| | BCL7B (ENSG00000106635) | 7:73536356-73557690 | → | Bcl7b (ENSMUSG00000029681) | 5:135168283-135181855 |
| | TBL2 (ENSG00000106638) | 7:73567537-73578791 | → | Tbl2 (ENSMUSG00000005374) | 5:135149657-135165760 |
| | MLXIPL (ENSG00000009950) | 7:73593194-73624543 | → | Mlxipl (ENSMUSG00000005373) | 5:135089890-135138382 |
| | VPS37D (ENSG00000176428) | 7:73667831-73672112 | → | Vps37d (ENSMUSG00000043614) | 5:135072900-135078266 |
| | DNAJC30 (ENSG00000176410) | 7:73680918-73683453 | → | Dnajc30 (ENSMUSG00000061118) | 5:135064202-135065862 |
| | BUD23 (ENSG00000071462) | 7:73683025-73705161 | → | Bud23 (ENSMUSG00000005378) | 5:135052957-135064959 |
| | STX1A (ENSG00000106089) | 7:73699206-73719672 | → | Stx1a (ENSMUSG00000007207) | 5:135023482-135051100 |
| | ABHD11 (ENSG00000106077) | 7:73736094-73738867 | → | Abhd11 (ENSMUSG00000040532) | 5:135009152-135012175 |
| | CLDN3 (ENSG00000165215) | 7:73768997-73770270 | → | Cldn3 (ENSMUSG00000070473) | 5:134986214-134987472 |
| | CLDN4 (ENSG00000189143) | 7:73799542-73832693 | → | Cldn13 (ENSMUSG00000008843) | 5:134914249-134915526 |
| | METTL27 (ENSG00000165171) | 7:73834590-73842516 | → | Mettl27 (ENSMUSG00000040557) | 5:134932368-134942637 |
| | TMEM270 (ENSG00000175877) | 7:73861159-73865890 | → | Tmem270 (ENSMUSG00000040576) | 5:134901593-134906733 |
| | ELN (ENSG00000049540) | 7:74027789-74069907 | → | Eln (ENSMUSG00000029675) | 5:134702593-134747323 |
| | LIMK1 (ENSG00000106683) | 7:74082933-74122525 | → | Limk1 (ENSMUSG00000029674) | 5:134656039-134688598 |
| | EIF4H (ENSG00000106682) | 7:74174245-74197101 | → | Eif4h (ENSMUSG00000040731) | 5:134619721-134639490 |

| | | | | | |
|---|---|---|---|---|---|
| | LAT2 (ENSG00000086730) | 7:74199652-74229834 | → | Lat2 (ENSMUSG00000040751) | 5:134600022-134615025 |
| | RFC2 (ENSG00000049541) | 7:74231499-74254458 | → | Rfc2 (ENSMUSG00000023104) | 5:134581366-134601805 |
| | CLIP2 (ENSG00000106665) | 7:74289475-74405943 | → | Clip2 (ENSMUSG00000063146) | 5:134489383-134552434 |
| | GTF2IRD1 (ENSG00000006704) | 7:74453790-74602604 | → | Gtf2ird1 (ENSMUSG00000023079) | 5:134357656-134456716 |
| **15q11.2** | NIPA1 (ENSG00000170113) | 15:22773063-22829789 | → | Nipa1 (ENSMUSG00000047037) | 7:55977567-56019954 |
| | NIPA2 (ENSG00000140157) | 15:22838641-22868384 | → | Nipa2 (ENSMUSG00000030452) | 7:55931287-55962476 |
| | CYFIP1 (ENSG00000273749) | 15:22867052-22981063 | → | Cyfip1 (ENSMUSG00000030447) | 7:55841745-55932602 |
| | TUBGCP5 (ENSG00000275835) | 15:22983192-23039572 | → | Tubgcp5 (ENSMUSG00000033790) | 7:55794154-55831677 |
| **15q13.3** | OTUD7A (ENSG00000169918) | 15:31475398-31870789 | → | Otud7a (ENSMUSG00000033510) | 7:63444751-63759028 |
| | CHRNA7 (ENSG00000175344) | 15:31923438-32173018 | → | Chrna7 (ENSMUSG00000030525) | 7:63098692-63212569 |
| **16p11.2** | SPN (ENSG00000197471) | 16:29662979-29670876 | → | Spn (ENSMUSG00000051457) | 7:127132232-127137823 |
| | QPRT (ENSG00000103485) | 16:29663279-29698699 | → | Qprt (ENSMUSG00000030674) | 7:127107114-127122226 |
| | C16orf54 (ENSG00000185905) | 16:29742463-29745990 | → | AI467606 (ENSMUSG00000045165) | 7:127091359-127093986 |
| | ZG16 (ENSG00000174992) | 16:29778256-29782973 | → | Zg16 (ENSMUSG00000049350) | 7:127050156-127087328 |
| | KIF22 (ENSG00000079616) | 16:29790719-29805385 | → | Kif22 (ENSMUSG00000030677) | 7:127027729-127042471 |
| | MAZ (ENSG00000103495) | 16:29806106-29811164 | → | Maz (ENSMUSG00000030678) | 7:127022130-127027037 |
| | PRRT2 (ENSG00000167371) | 16:29811382-29815892 | → | Prrt2 (ENSMUSG00000045114) | 7:127017531-127021211 |
| | AC009133.6 (ENSG00000280893) | 16:29812261-29820092 | | No homologues | |
| | PAGR1 (ENSG00000280789) | 16:29816152-29822489 | → | Gm42742 (ENSMUSG00000107068) | 7:127015051-127021097 |
| | | | → | Pagr1b (ENSMUSG00000092534) | 7:127001480-127016484 |
| | | | → | Pagr1a (ENSMUSG00000030680) | 7:127015033-127017352 |
| | AC120114.4 (ENSG00000281348) | 16:29817239-29830627 | | No homologues | |
| | MVP (ENSG00000013364) | 16:29820394-29848039 | → | Mvp (ENSMUSG00000030681) | 7:126986860-127014621 |
| | CDIPT (ENSG00000103502) | 16:29858357-29863414 | → | Cdipt (ENSMUSG00000030682) | 7:126975914-126980501 |
| | SEZ6L2 (ENSG00000174938) | 16:29871159-29899547 | → | Sez6l2 (ENSMUSG00000030683) | 7:126950563-126970606 |
| | ASPHD1 (ENSG00000174939) | 16:29900375-29919864 | → | Asphd1 (ENSMUSG00000046378) | 7:126945567-126949582 |
| | KCTD13 (ENSG00000174943) | 16:29905012-29926236 | → | Kctd13 (ENSMUSG00000030685) | 7:126928879-126945631 |
| | TMEM219 (ENSG00000149932) | 16:29940885-29973050 | → | Tmem219 (ENSMUSG00000060538) | 7:126886171-126922917 |
| | TAOK2 (ENSG00000149930) | 16:29973868-29992261 | → | Taok2 (ENSMUSG00000059981) | 7:126865678-126884703 |
| | HIRIP3 (ENSG00000149929) | 16:29992330-29996074 | → | Hirip3 (ENSMUSG00000042606) | 7:126861972-126865377 |
| | INO80E (ENSG00000169592) | 16:29995715-30005793 | → | Ino80e (ENSMUSG00000030689) | 7:126850960-126862377 |
| | DOC2A (ENSG00000149927) | 16:30005514-30023270 | → | Doc2a (ENSMUSG00000052301) | 7:126847416-126852705 |
| | C16orf92 (ENSG00000167194) | 16:30023334-30027736 | → | 4930451I11Rik (ENSMUSG00000045989) | 7:126830468-126831639 |
| | TLCD3B (ENSG00000149926) | 16:30024427-30052978 | → | Fam57b (ENSMUSG00000058966) | 7:126797668-126830219 |
| | AC093512.2 (ENSG00000285043) | 16:30053090-30070420 | → | Aldoa (ENSMUSG00000030695) | 7:126795234-126800751 |
| | | | → | Aldoart2 (ENSMUSG00000063129) | 12:55565239-55566896 |
| | | | → | Aldoart1 (ENSMUSG00000059343) | 4:72850583-72852634 |
| | ALDOA (ENSG00000149925) | 16:30064164-30070457 | → | Aldoa (ENSMUSG00000030695) | 7:126795234-126800751 |
| | | | → | Aldoart2 (ENSMUSG00000063129) | 12:55565239-55566896 |
| | | | → | Aldoart1 (ENSMUSG00000059343) | 4:72850583-72852634 |
| | PPP4C (ENSG00000149923) | 16:30075978-30085376 | → | Ppp4c (ENSMUSG00000030697) | 7:126785866-126792496 |
| | TBX6 (ENSG00000149922) | 16:30085793-30091887 | → | Tbx6 (ENSMUSG00000030699) | 7:126781483-126785560 |
| | YPEL3 (ENSG00000090238) | 16:30092314-30096915 | → | Ypel3 (ENSMUSG00000042675) | 7:126776955-126780514 |
| | GDPD3 (ENSG00000102886) | 16:30104810-30113537 | → | Gdpd3 (ENSMUSG00000030703) | 7:126766334-126775649 |
| | MAPK3 (ENSG00000102882) | 16:30114105-30123506 | → | Mapk3 (ENSMUSG00000063065) | 7:126759601-126765819 |
| | CORO1A (ENSG00000102879) | 16:30182827-30189076 | → | Coro1a (ENSMUSG00000030707) | 7:126699773-126707787 |
| **16p13.2** | TMEM114 (ENSG00000232258) | 16:8537605-8590193 | → | Tmem114 (ENSMUSG00000022715) | 16:8409275-8425136 |
| | METTL22 (ENSG00000067365) | 16:8621683-8649654 | → | Mettl22 (ENSMUSG00000039345) | 16:8470788-8490684 |
| | ABAT (ENSG00000183044) | 16:8674596-8784575 | → | Abat (ENSMUSG00000057880) | 16:8513429-8621568 |
| | TMEM186 (ENSG00000184857) | 16:8780384-8797642 | → | Tmem186 (ENSMUSG00000043140) | 16:8633229-8637712 |
| | PMM2 (ENSG00000140650) | 16:8788823-8849325 | → | Pmm2 (ENSMUSG00000022711) | 16:8637674-8662467 |
| | CARHSP1 (ENSG00000153048) | 16:8852942-8869012 | → | Carhsp1 (ENSMUSG00000008393) | 16:8658580-8672155 |
| | LITAFD (ENSG00000283516) | 16:8881634-8885351 | → | Gm5767 (ENSMUSG00000107252) | 16:8680770-8683911 |
| | USP7 (ENSG00000187555) | 16:8892097-8964514 | → | Usp7 (ENSMUSG00000022710) | 16:8689595-8792308 |
| | C16orf72 (ENSG00000182831) | 16:9091644-9121635 | → | 1810013L24Rik (ENSMUSG00000022507) | 16:8830100-8858922 |
| **17p11.2** | MPRIP (ENSG00000133030) | 17:17042545-17217679 | → | Mprip (ENSMUSG00000005417) | 11:59661305-59780860 |
| | PLD6 (ENSG00000179598) | 17:17200995-17206333 | → | Pld6 (ENSMUSG00000043648) | 11:59783897-59787645 |
| | AC055811.2 (ENSG00000264187) | 17:17202649-17237185 | | No homologues | |
| | FLCN (ENSG00000154803) | 17:17212212-17237188 | → | Flcn (ENSMUSG00000032633) | 11:59791408-59810016 |
| | COPS3 (ENSG00000141030) | 17:17246616-17281273 | → | Cops3 (ENSMUSG00000019373) | 11:59817795-59839838 |
| | NT5M (ENSG00000205309) | 17:17303335-17347663 | → | Nt5m (ENSMUSG00000032615) | 11:59839447-59880968 |
| | MED9 (ENSG00000141026) | 17:17476994-17493221 | → | Med9 (ENSMUSG00000061650) | 11:59948206-59962205 |
| | RASD1 (ENSG00000108551) | 17:17494437-17496395 | → | Rasd1 (ENSMUSG00000049892) | 11:59963181-59964944 |
| | PEMT (ENSG00000133027) | 17:17505563-17591708 | → | Pemt (ENSMUSG00000000301) | 11:59970614-60046489 |
| | RAI1 (ENSG00000108557) | 17:17681458-17811453 | → | Rai1 (ENSMUSG00000062115) | 11:60105013-60199197 |
| | SREBF1 (ENSG00000072310) | 17:17810399-17837011 | → | Srebf1 (ENSMUSG00000020538) | 11:60199089-60222581 |
| | TOM1L2 (ENSG00000175662) | 17:17843511-17972422 | → | Tom1l2 (ENSMUSG00000000538) | 11:60226714-60352905 |
| | DRC3 (ENSG00000171962) | 17:17972813-18016889 | → | Drc3 (ENSMUSG00000056598) | 11:60353329-60394341 |
| | ATPAF2 (ENSG00000171953) | 17:17977409-18039209 | → | Atpaf2 (ENSMUSG00000042709) | 11:60400626-60418457 |
| | GID4 (ENSG00000141034) | 17:18039408-18068405 | → | Gid4 (ENSMUSG00000018415) | 11:60417145-60450927 |
| | DRG2 (ENSG00000108591) | 17:18087892-18107970 | → | Drg2 (ENSMUSG00000020537) | 11:60454591-60468754 |

| | | | | | |
|---|---|---|---|---|---|
| | MYO15A (ENSG00000091536) | 17:18108706-18179802 | → | Myo15 (ENSMUSG00000042678) | 11:60469339-60528369 |
| | ALKBH5 (ENSG00000091542) | 17:18183078-18209954 | → | Alkbh5 (ENSMUSG00000042650) | 11:60536381-60558512 |
| | LLGL1 (ENSG00000131899) | 17:18225635-18244875 | → | Llgl1 (ENSMUSG00000020536) | 11:60699723-60714186 |
| | FLII (ENSG00000177731) | 17:18244815-18258738 | → | Flii (ENSMUSG00000002812) | 11:60714123-60727263 |
| | MIEF2 (ENSG00000177427) | 17:18260597-18266552 | → | Mief2 (ENSMUSG00000018599) | 11:60728398-60732951 |
| | TOP3A (ENSG00000177302) | 17:18271428-18315007 | → | Top3a (ENSMUSG00000002814) | 11:60740058-60777365 |
| | SMCR8 (ENSG00000176994) | 17:18315293-18328056 | → | Smcr8 (ENSMUSG00000049323) | 11:60777524-60788287 |
| **17q12** | ZNHIT3 (ENSG00000273611) | 17:36486629-36499310 | → | Znhit3 (ENSMUSG00000020526) | 11:84910950-84916366 |
| | MYO19 (ENSG00000278259) | 17:36495633-36543435 | → | Myo19 (ENSMUSG00000020527) | 11:84880148-84911226 |
| | PIGW (ENSG00000277161) | 17:36534987-36539310 | → | Pigw (ENSMUSG00000045140) | 11:84876315-84880285 |
| | GGNBP2 (ENSG00000278311) | 17:36544912-36589848 | → | Ggnbp2 (ENSMUSG00000020530) | 11:84832361-84870817 |
| | DHRS11 (ENSG00000278535) | 17:36591879-36600804 | → | Dhrs11 (ENSMUSG00000034449) | 11:84820856-84828994 |
| | MRM1 (ENSG00000278619) | 17:36601583-36608964 | → | Mrm1 (ENSMUSG00000018405) | 11:84813061-84819515 |
| | LHX1 (ENSG00000273706) | 17:36936785-36944612 | → | Lhx1 (ENSMUSG00000018698) | 11:84518284-84525535 |
| | AATF (ENSG00000275700) | 17:36948954-37056871 | → | Aatf (ENSMUSG00000018697) | 11:84422855-84513522 |
| | ACACA (ENSG00000278540) | 17:37084992-37406836 | → | Acaca (ENSMUSG00000020532) | 11:84129672-84401651 |
| | C17orf78 (ENSG00000278505) | 17:37375985-37392708 | → | Gm11437 (ENSMUSG00000051452) | 11:84148351-84167476 |
| | TADA2A (ENSG00000276234) | 17:37406886-37479725 | → | Tada2a (ENSMUSG00000018651) | 11:84078920-84129600 |
| | DUSP14 (ENSG00000276023) | 17:37489891-37513501 | → | Dusp14 (ENSMUSG00000018648) | 11:84048041-84069261 |
| | SYNRG (ENSG00000275066) | 17:37514807-37609472 | → | Synrg (ENSMUSG00000034940) | 11:83964428-84044578 |
| | DDX52 (ENSG00000278053) | 17:37609739-37643446 | → | Ddx52 (ENSMUSG00000020677) | 11:83942062-83963088 |
| | HNF1B (ENSG00000275410) | 17:37686431-37745105 | → | Hnf1b (ENSMUSG00000020679) | 11:83850063-83905819 |
| **Xq27.3** | FMR1 (ENSG00000102081) | X:147911951-147951125 | → | Fmr1 (ENSMUSG00000000838) | X:68678541-68717963 |
| **Xq28** | MECP2 (ENSG00000169057) | X:154021573-154137103 | → | Mecp2 (ENSMUSG00000031393) | X:74026592-74135363 |
