## Extended Data Table 7 for "Autism in a dish: ES cell models of autism with copy number variations reveal cell-type-specific vulnerability"

**Extended Data Table7. Gene expression in targeted loci.**

| CNVs | Genes | Average | logFC | P-value | |
|---|---|---|---|---|---|
| 1q21.1dup | *Acp6* | 0.039564957 | -0.441555157 | 0.19710355 | |
| | *Bcl9* | 0.550569496 | 1.203300946 | 1.66E-07 | **** |
| | *Chd1l* | 0.022094716 | -0.892865432 | 0.005691529 | *** |
| | *Fmo5* | 0.00385373 | 0.445550786 | 0.76645674 | |
| | *Gja5* | 0.000513831 | 0.615475788 | 0.894654509 | |
| | *Gja8* | - | - | - | |
| | *Gm24435* | - | - | - | |
| | *Gpr89* | 0.239188148 | 0.765416415 | 0.002012707 | *** |
| | *Pde4dip* | 0.182923696 | 0.669067765 | 0.010291665 | ** |
| | *Prkab2* | 0.143101824 | 1.26589135 | 5.64E-07 | **** |
| | *RP23-290C4.1* | - | - | - | |
| | *RP23-394K8.3* | - | - | - | |
| | *RP23-394K8.5* | - | - | - | |
| | *RP23-97A19.1* | - | - | - | |
| | *RP23-97A19.2* | - | - | - | |
| | *RP24-316D12.2* | - | - | - | |
| 2p16.3 | *Nrxn1* | 1.393736504 | 0.69866876 | 0.038626965 | ** |
| 3q29 | *0610012G03Rik* | 0.483847307 | -1.110100689 | 9.65E-06 | **** |
| | *Bdh1* | 0.065835994 | -1.441558335 | 1.47E-07 | **** |
| | *Bex6* | - | - | - | |
| | *Cep19* | 0.128521941 | -1.090240536 | 4.91E-05 | **** |
| | *Dlg1* | 0.078436185 | -1.081433435 | 0.000121639 | **** |
| | *Fbxo45* | 0.057960875 | -0.747319417 | 0.022641868 | ** |
| | *Gm15694* | - | - | - | |
| | *Gm15696* | - | - | - | |
| | *Gm15703* | - | - | - | |
| | *Gm15729* | - | - | - | |
| | *Gm15743* | 0.015120228 | -0.177113532 | 0.965302352 | |
| | *Gm20056* | - | - | - | |
| | *Gm24879* | - | - | - | |
| | *Gm25848* | - | - | - | |
| | *Gm26269* | - | - | - | |
| | *Gm26419* | - | - | - | |
| | *Gm5405* | - | - | - | |
| | *Hmgb1-ps6* | - | - | - | |
| | *Mfi2* | 0.001260019 | 0.357923743 | 1 | |
| | *Mir1946a* | - | - | - | |
| | *Ncbp2* | 0.255783863 | -1.162316788 | 4.63E-06 | **** |
| | *NCBP2-AS2* | - | - | - | |
| | *Nrros* | 0.000945014 | -2.165638213 | 0.027141199 | ** |
| | *Pak2* | 0.292954424 | -1.397365456 | 1.45E-08 | **** |
| | *Pcyt1a* | 0.082531246 | -1.332778011 | 6.29E-07 | **** |
| | *Pigx* | 0.293584434 | -0.925454977 | 0.000479654 | **** |
| | *Pigz* | 0.002520038 | -0.057113756 | 1 | |
| | *Rnf168* | 0.083791265 | -1.164463093 | 2.15E-05 | **** |
| | *Senp5* | 0.082846251 | -0.941978045 | 0.001145388 | *** |
| | *Slc51a* | 0.00063001 | -0.642076257 | 0.863598032 | |
| | *Smco1* | 0.000315005 | -0.227038758 | 1 | |
| | *Tctex1d2* | 0.154037326 | -0.736105454 | 0.011628988 | ** |
| | *Tfrc* | 0.212313206 | -0.524103335 | 0.11365276 | |
| | *Tm4sf19* | - | - | - | |
| | *Tnk2* | 0.022995347 | 0.263596361 | 0.8117429 | |
| | *Ubxn7* | 0.139232103 | -0.713550726 | 0.017015771 | ** |
| | *Wdr53* | 0.028665433 | -1.061028806 | 0.00176745 | *** |
| | *Zdhhc19* | - | - | - | |
| 7q11.23dup | *Adhd11* | - | - | - | |
| | *Adhd11os* | - | - | - | |
| | *Baz1b* | 0.448068312 | -0.418041494 | 0.124934293 | |
| | *Bcl7b* | 0.299292539 | 0.204611333 | 0.630821933 | |
| | *Cldn13* | - | - | - | |
| | *Cldn3* | 0.072963438 | 1.136026999 | 0.012332288 | ** |
| | *Cldn4* | 0.100344511 | 0.971823634 | 0.036344855 | ** |
| | *Clip2* | 0.265264038 | 1.4866467 | 3.96E-09 | **** |
| | *Dnajc30* | 0.340126741 | 0.48953498 | 0.083963026 | * |
| | *Eif4h* | 1.097617007 | 0.022608603 | 1 | |
| | *Eln* | 0.014719305 | -0.570895748 | 0.225032876 | |
| | *Fkbp6* | 0.010287686 | -0.065148936 | 1 | |
| | *Fzd9* | 0.004748163 | -2.187055605 | 1.57E-08 | **** |
| | *Gm16020* | - | - | - | |
| | *Lat2* | 0.017251659 | 0.80611775 | 0.056528601 | * |
| | *Limk1* | 0.282040881 | 1.222399096 | 1.16E-06 | **** |
| | *Mir7228* | - | - | - | |
| | *Mlxipl* | 0.003165442 | 0.189664963 | 0.980387649 | |
| | *Rfc2* | 0.383176752 | 0.161790918 | 0.732797826 | |
| | *Stx1a* | 0.197840124 | 1.452007594 | 2.00E-08 | **** |
| | *Syna* | 0.002374081 | 3.382310041 | 0.00621638 | *** |
| | *Tbl2* | 0.059668581 | 0.223453276 | 0.647807782 | |
| | *Trim50* | - | - | - | |
| | *Vps37d* | 0.178056111 | 1.243176747 | 1.63E-06 | **** |
| | *Wbscr22* | 0.213983878 | 0.153079109 | 0.774577563 | |
| | *Wbscr25* | 0.004115075 | 0.967272541 | 0.256663885 | |
| | *Wbscr27* | 0.029280338 | -0.78589028 | 0.008285569 | ** |
| | *Wbscr28* | 0.000158272 | -0.617689959 | 1 | |
| 15q11.2 | *Cyfip1* | 0.16271762 | -0.814945149 | 0.004695296 | *** |
| | *Gm17907* | - | - | - | |
| | *Nipa1* | 0.09794274 | -0.080138309 | 1 | |
| | *Nipa2* | 0.235296702 | -0.764457329 | 0.007572227 | *** |
| | *Olfr732* | - | - | - | |
| | *Olfr733* | - | - | - | |

| | | | | | |
|---|---|---|---|---|---|
| | *Olfr734* | - | - | - | |
| | *Tubgcp5* | 0.150230896 | -0.259377062 | 0.607893066 | |
| 15q13.3 | *4930554H23Rik* | - | - | - | |
| | *Chrna7* | 0.001281971 | -1.431132083 | 0.374751688 | |
| | *Otud7a* | 0.003845912 | -1.071236138 | 0.440772052 | |
| 16p11.2 | *4930451I11Rik* | - | - | - | |
| | *AI467606* | - | - | - | |
| | *Aldoa* | 1.044946979 | -0.245679819 | 0.61907321 | |
| | *Asphd1* | 0.042400411 | 1.215532595 | 0.002871621 | *** |
| | *Cdipt* | 0.214791557 | -0.099407438 | 1 | |
| | *Coro1a* | 0.109906329 | 1.177319159 | 0.000103719 | **** |
| | *Doc2a* | 0.010600103 | 1.576482579 | 0.025307462 | ** |
| | *Fam57b* | 0.490394231 | 0.882784439 | 0.000766187 | **** |
| | *Gdpd3* | 0.005579001 | -1.54235365 | 0.008636018 | *** |
| | *Hirip3* | 0.118832732 | -0.239194991 | 0.694334535 | |
| | *Ino80e* | 0.281181675 | -0.066039811 | 1 | |
| | *Kctd13* | 0.403361808 | 0.879931253 | 0.00088991 | **** |
| | *Kif22* | 0.063600617 | -0.657808425 | 0.07508426 | * |
| | *Mapk3* | 0.255518268 | 0.172752144 | 0.753647216 | |
| | *Maz* | 0.482583629 | 0.810926056 | 0.002138943 | *** |
| | *Mvp* | 0.03682141 | -0.684357006 | 0.110629249 | |
| | *Pagr1a* | 0.010600103 | 1.16144508 | 0.166181111 | |
| | *Ppp4c* | 0.592489958 | -0.224633022 | 0.677758066 | |
| | *Prrt2* | 0.0005579 | 0.839516985 | 1 | |
| | *Qprt* | 0.002231601 | -0.777154375 | 0.594287447 | |
| | *Sez6l2* | 0.205307255 | 0.792085155 | 0.006977593 | *** |
| | *Spn* | 0 | 0.424479486 | 1 | |
| | *Taok2* | 0.232644362 | 1.071142686 | 8.22E-05 | **** |
| | *Tbx6* | 0.003347401 | 1.646871907 | 0.258754674 | |
| | *Tmem219* | 0.18578075 | 0.20641299 | 0.673085978 | |
| | *Ypel3* | 0.768228505 | 0.54905891 | 0.05324585 | * |
| | *Zg16* | 0 | 0.424479486 | 1 | |
| 16p13.2 | *1810013L24Rik* | 0.11115871 | -0.286244862 | 0.512583677 | |
| | *Abat* | 0.079150957 | -2.515224781 | 1.20E-20 | **** |
| | *Carhsp1* | 0.586808816 | -0.929819655 | 0.000718365 | **** |
| | *Gm5767* | - | - | - | |
| | *Mettl22* | 0.070714804 | -0.603015129 | 0.075217122 | ** |
| | *Pmm2* | 0.05260189 | -1.536711399 | 7.56E-08 | **** |
| | *Tmem186* | 0.022827235 | -1.30303552 | 8.98E-05 | **** |
| | *Usp7* | 0.162519989 | -1.213195681 | 8.34E-06 | **** |
| 17p11.2 | *Alkbh5* | 0.222280153 | -0.88604906 | 0.000444951 | **** |
| | *Atpaf2* | 0.074000998 | -1.28464002 | 9.66E-07 | **** |
| | *Cops3* | 0.408529857 | -0.648257417 | 0.011462905 | ** |
| | *Drg2* | 0.078712672 | -1.333070347 | 2.03E-07 | **** |
| | *Flcn* | 0.062360392 | -0.937042638 | 0.001006289 | *** |
| | *Flii* | 0.052937044 | -1.509532136 | 1.13E-08 | **** |
| | *Gid4* | 0.057648718 | -1.468466583 | 2.65E-08 | **** |
| | *Llgl1* | 0.032427404 | -1.908459444 | 4.41E-12 | **** |
| | *Lrrc48* | 0.003880202 | -0.914585546 | 0.167936604 | |
| | *Med9os* | 0.008591876 | -2.2608978 | 1.99E-10 | **** |
| | *Mief2* | 0.015520809 | -1.678901629 | 1.97E-07 | **** |
| | *Mprip* | 0.18458676 | -2.189142381 | 1.10E-19 | **** |
| | *Myo15* | 0.000831472 | 0.637955477 | 0.912551387 | |
| | *Nt5m* | 0.097005054 | -1.385149884 | 3.85E-08 | **** |
| | *Pemt* | 0.020786797 | -0.823775258 | 0.018973037 | ** |
| | *Pld6* | 0.004711674 | 0.222917977 | 0.985292098 | |
| | *Rai1* | 0.047116741 | -2.353582945 | 2.41E-19 | **** |
| | *Rasd1* | 0.004711674 | -1.307596739 | 0.043702688 | ** |
| | *Smcr6* | - | - | - | |
| | *Srebf1* | 0.047948212 | -1.952524029 | 8.45E-14 | **** |
| | *Tom1l2* | 0.034090348 | -1.785781718 | 1.33E-10 | **** |
| | *Top3a* | 0.01025482 | -1.728826854 | 1.64E-06 | **** |
| 17q12 | *1700109G15Rik* | 0.000230131 | -0.986803263 | 0.835337781 | |
| | *4930502E09Rik* | - | - | - | |
| | *Aatf* | 0.10424918 | -1.231717138 | 1.17E-06 | **** |
| | *Acaca* | 0.093663171 | -1.636306016 | 5.61E-11 | **** |
| | *Ddx52* | 0.204356009 | -1.239704563 | 3.16E-07 | **** |
| | *Dhrs11* | 0.019791235 | 0.517540778 | 0.312293854 | |
| | *Dusp14* | 0.093202909 | -0.351214689 | 0.305775242 | |
| | *Ggnbp2* | 0.565661115 | -0.887016125 | 0.000296691 | **** |
| | *Gm11431* | - | - | - | |
| | *Gm11437* | - | - | - | |
| | *Gm23564* | - | - | - | |
| | *Gm37391* | - | - | - | |
| | *Hnf1b* | 0.015188622 | -1.117111285 | 0.002335858 | *** |
| | *Lhx1* | 0.05039861 | -1.697296646 | 1.84E-07 | **** |
| | *Lhx1os* | 0.096885 | -1.899910094 | 2.55E-11 | **** |
| | *Mrm1* | 0.015648884 | -0.6152444 | 0.155624941 | **** |
| | *Myo19* | 0.020481627 | -0.986803263 | 0.003703651 | *** |
| | *Pigw* | 0.01449823 | -1.183200476 | 0.000879977 | **** |
| | *Synrg* | 0.071570629 | -2.543751387 | 2.33E-24 | **** |
| | *Tada2a* | 0.126111591 | -0.937803044 | 0.000290399 | **** |
| | *Znhit3* | 0.234733253 | -1.15761767 | 2.01E-06 | **** |
| Xq27.3 | *Fmr1* | 0 | -11.59712671 | 1.56E-124 | **** |
| Xq28 | *Mecp2* | 0 | -11.36437822 | 2.24E-119 | **** |

| **p-value legend** | |
|---|---|
| **** | p<0.001 |
| *** | p<0.01 |
| ** | p<0.05 |
| * | p<0.1 |
