## Extended Data Table 8 for "Autism in a dish: ES cell models of autism with copy number variations reveal cell-type-specific vulnerability"

**Extended Data Table 8. Cell-type (cluster) specific Gene Ontology (GO) terms.**

| GO (#0) | Category | Description | Count | % | Log10(P) | Log10(q) |
|---|---|---|---|---|---|---|
| R-MMU-1474244 | Reactome Gene Sets | Extracellular matrix organization | 42 | 14.38 | -30.41 | -26.15 |
| GO:0001501 | GO Biological Processes | skeletal system development | 53 | 18.15 | -28.62 | -24.66 |
| GO:0001568 | GO Biological Processes | blood vessel development | 52 | 17.81 | -21.5 | -17.94 |
| GO:2000147 | GO Biological Processes | positive regulation of cell motility | 43 | 14.73 | -18.66 | -15.36 |
| GO:0032963 | GO Biological Processes | collagen metabolic process | 22 | 7.53 | -18.58 | -15.32 |
| GO:0070848 | GO Biological Processes | response to growth factor | 44 | 15.07 | -17.85 | -14.73 |
| GO:0001503 | GO Biological Processes | ossification | 35 | 11.99 | -17.62 | -14.61 |
| R-MMU-1474290 | Reactome Gene Sets | Collagen formation | 19 | 6.51 | -17.43 | -14.45 |
| GO:0050673 | GO Biological Processes | epithelial cell proliferation | 36 | 12.33 | -16.98 | -14.04 |
| GO:0048729 | GO Biological Processes | tissue morphogenesis | 44 | 15.07 | -16.34 | -13.46 |
| R-MMU-381426 | Reactome Gene Sets | Regulation of Insulin-like Growth Factor (IGF) transport and uptake by Insulin-like Growth Factor Bi | 20 | 6.85 | -15.52 | -12.68 |
| GO:0042060 | GO Biological Processes | wound healing | 30 | 10.27 | -13.93 | -11.14 |
| GO:0031589 | GO Biological Processes | cell-substrate adhesion | 29 | 9.93 | -13.86 | -11.1 |
| GO:0048589 | GO Biological Processes | developmental growth | 43 | 14.73 | -13.35 | -10.64 |
| GO:0051271 | GO Biological Processes | negative regulation of cellular component movement | 26 | 8.9 | -12.08 | -9.49 |
| R-MMU-2129379 | Reactome Gene Sets | Molecules associated with elastic fibres | 11 | 3.77 | -12.02 | -9.44 |
| GO:0048706 | GO Biological Processes | embryonic skeletal system development | 18 | 6.16 | -11.8 | -9.25 |
| mmu05205 | KEGG Pathway | Proteoglycans in cancer | 21 | 7.19 | -11.62 | -9.08 |
| GO:0008285 | GO Biological Processes | negative regulation of cell proliferation | 37 | 12.67 | -11.43 | -8.92 |
| GO:0016055 | GO Biological Processes | Wnt signaling pathway | 28 | 9.59 | -10.45 | -8.02 |

| GO (#1) | Category | Description | Count | % | Log10(P) | Log10(q) |
|---|---|---|---|---|---|---|
| R-MMU-975956 | Reactome Gene Sets | Nonsense Mediated Decay (NMD) independent of the Exon Junction Complex (EJC) | 52 | 18.06 | -71.8 | -67.54 |
| R-MMU-1640170 | Reactome Gene Sets | Cell Cycle | 50 | 17.36 | -25.33 | -22.47 |
| GO:0042254 | GO Biological Processes | ribosome biogenesis | 38 | 13.19 | -24.7 | -21.85 |
| GO:1903047 | GO Biological Processes | mitotic cell cycle process | 50 | 17.36 | -22.25 | -19.45 |
| GO:0042273 | GO Biological Processes | ribosomal large subunit biogenesis | 15 | 5.21 | -12.34 | -9.75 |
| GO:0045786 | GO Biological Processes | negative regulation of cell cycle | 31 | 10.76 | -12.33 | -9.75 |
| GO:0006260 | GO Biological Processes | DNA replication | 23 | 7.99 | -12.12 | -9.56 |
| GO:0051052 | GO Biological Processes | regulation of DNA metabolic process | 29 | 10.07 | -11.16 | -8.62 |
| GO:0042255 | GO Biological Processes | ribosome assembly | 13 | 4.51 | -10.7 | -8.21 |
| R-MMU-8852276 | Reactome Gene Sets | The role of GTSE1 in G2/M progression after G2 checkpoint | 9 | 3.12 | -9.65 | -7.23 |
| GO:0009123 | GO Biological Processes | nucleoside monophosphate metabolic process | 21 | 7.29 | -9.08 | -6.68 |
| GO:0065004 | GO Biological Processes | protein-DNA complex assembly | 16 | 5.56 | -8.48 | -6.14 |
| GO:1903046 | GO Biological Processes | meiotic cell cycle process | 17 | 5.9 | -8.44 | -6.11 |
| GO:0006275 | GO Biological Processes | regulation of DNA replication | 13 | 4.51 | -8.41 | -6.07 |
| GO:1990823 | GO Biological Processes | response to leukemia inhibitory factor | 19 | 6.6 | -6.96 | -4.78 |
| GO:0051785 | GO Biological Processes | positive regulation of nuclear division | 10 | 3.47 | -6.88 | -4.7 |
| R-MMU-176417 | Reactome Gene Sets | Phosphorylation of Emi1 | 4 | 1.39 | -6.71 | -4.55 |
| GO:0001701 | GO Biological Processes | in utero embryonic development | 25 | 8.68 | -6.64 | -4.49 |
| GO:0034502 | GO Biological Processes | protein localization to chromosome | 10 | 3.47 | -6.46 | -4.33 |
| GO:0031570 | GO Biological Processes | DNA integrity checkpoint | 11 | 3.82 | -6.25 | -4.13 |

| GO (#2) | Category | Description | Count | % | Log10(P) | Log10(q) |
|---|---|---|---|---|---|---|
| GO:0032990 | GO Biological Processes | cell part morphogenesis | 55 | 19.16 | -24.37 | -20.11 |
| GO:0051129 | GO Biological Processes | negative regulation of cellular component organization | 42 | 14.63 | -13.53 | -10.34 |
| GO:0050808 | GO Biological Processes | synapse organization | 31 | 10.8 | -12.38 | -9.26 |
| GO:0007420 | GO Biological Processes | brain development | 35 | 12.2 | -11.23 | -8.23 |
| R-MMU-422475 | Reactome Gene Sets | Axon guidance | 22 | 7.67 | -10.88 | -7.95 |
| GO:0051961 | GO Biological Processes | negative regulation of nervous system development | 24 | 8.36 | -9.42 | -6.7 |
| R-MMU-399956 | Reactome Gene Sets | CRMPs in Sema3A signaling | 7 | 2.44 | -9.22 | -6.52 |
| GO:0001764 | GO Biological Processes | neuron migration | 17 | 5.92 | -8.92 | -6.3 |
| GO:0048024 | GO Biological Processes | regulation of mRNA splicing, via spliceosome | 13 | 4.53 | -8.48 | -5.92 |
| GO:0097479 | GO Biological Processes | synaptic vesicle localization | 16 | 5.57 | -7.95 | -5.46 |
| GO:0006333 | GO Biological Processes | chromatin assembly or disassembly | 14 | 4.88 | -7.76 | -5.32 |
| GO:0097106 | GO Biological Processes | postsynaptic density organization | 8 | 2.79 | -7.61 | -5.2 |
| GO:0034643 | GO Biological Processes | establishment of mitochondrion localization, microtubule-mediated | 6 | 2.09 | -6.76 | -4.48 |
| GO:0035249 | GO Biological Processes | synaptic transmission, glutamatergic | 11 | 3.83 | -6.55 | -4.29 |
| mmu04728 | KEGG Pathway | Dopaminergic synapse | 12 | 4.18 | -6.34 | -4.12 |
| R-MMU-380320 | Reactome Gene Sets | Recruitment of NuMA to mitotic centrosomes | 10 | 3.48 | -6.29 | -4.08 |
| GO:1903827 | GO Biological Processes | regulation of cellular protein localization | 23 | 8.01 | -5.81 | -3.65 |
| GO:0097091 | GO Biological Processes | synaptic vesicle clustering | 5 | 1.74 | -5.4 | -3.27 |
| GO:0007610 | GO Biological Processes | behavior | 27 | 9.41 | -5.29 | -3.18 |
| GO:2000171 | GO Biological Processes | negative regulation of dendrite development | 6 | 2.09 | -5.06 | -3 |

| GO (#3) | Category | Description | Count | % | Log10(P) | Log10(q) |
|---|---|---|---|---|---|---|
| GO:0060322 | GO Biological Processes | head development | 45 | 15.9 | -17.41 | -13.24 |
| GO:0045596 | GO Biological Processes | negative regulation of cell differentiation | 46 | 16.25 | -15.66 | -12 |
| GO:0010001 | GO Biological Processes | glial cell differentiation | 24 | 8.48 | -13.36 | -9.95 |
| GO:0022612 | GO Biological Processes | gland morphogenesis | 19 | 6.71 | -12.5 | -9.28 |
| GO:0030855 | GO Biological Processes | epithelial cell differentiation | 36 | 12.72 | -12.21 | -9.04 |
| GO:0030278 | GO Biological Processes | regulation of ossification | 22 | 7.77 | -11.74 | -8.69 |
| GO:0050673 | GO Biological Processes | epithelial cell proliferation | 29 | 10.25 | -11.7 | -8.66 |
| GO:0001655 | GO Biological Processes | urogenital system development | 25 | 8.83 | -10.67 | -7.77 |
| GO:0007423 | GO Biological Processes | sensory organ development | 33 | 11.66 | -10.58 | -7.69 |
| GO:0048863 | GO Biological Processes | stem cell differentiation | 20 | 7.07 | -10.11 | -7.32 |
| GO:0003002 | GO Biological Processes | regionalization | 25 | 8.83 | -9.99 | -7.22 |
| GO:0045165 | GO Biological Processes | cell fate commitment | 21 | 7.42 | -9.24 | -6.54 |
| GO:0048589 | GO Biological Processes | developmental growth | 34 | 12.01 | -8.29 | -5.68 |
| GO:0008544 | GO Biological Processes | epidermis development | 21 | 7.42 | -8.08 | -5.5 |
| GO:0060485 | GO Biological Processes | mesenchyme development | 19 | 6.71 | -8.01 | -5.44 |
| GO:0021846 | GO Biological Processes | cell proliferation in forebrain | 8 | 2.83 | -7.66 | -5.15 |
| GO:0008285 | GO Biological Processes | negative regulation of cell proliferation | 30 | 10.6 | -7.55 | -5.06 |
| GO:0021872 | GO Biological Processes | forebrain generation of neurons | 11 | 3.89 | -7.36 | -4.9 |
| GO:0001570 | GO Biological Processes | vasculogenesis | 11 | 3.89 | -7.11 | -4.7 |
| GO:0021915 | GO Biological Processes | neural tube development | 15 | 5.3 | -7.1 | -4.7 |

| GO (#4) | Category | Description | Count | % | Log10(P) | Log10(q) |
|---|---|---|---|---|---|---|
| GO:0030036 | GO Biological Processes | actin cytoskeleton organization | 40 | 13.7 | -13.8 | -9.53 |
| R-MMU-72662 | Reactome Gene Sets | Activation of the mRNA upon binding of the cap-binding complex and eIFs, and subsequent bindir | 13 | 4.45 | -11.72 | -8.06 |
| GO:0030335 | GO Biological Processes | positive regulation of cell migration | 33 | 11.3 | -11.61 | -8.04 |
| GO:0031589 | GO Biological Processes | cell-substrate adhesion | 26 | 8.9 | -11.38 | -7.96 |
| mmu04670 | KEGG Pathway | Leukocyte transendothelial migration | 15 | 5.14 | -9.87 | -6.81 |
| R-MMU-445355 | Reactome Gene Sets | Smooth Muscle Contraction | 9 | 3.08 | -9.75 | -6.74 |
| GO:0051017 | GO Biological Processes | actin filament bundle assembly | 15 | 5.14 | -7.67 | -4.91 |
| GO:0044419 | GO Biological Processes | interspecies interaction between organisms | 23 | 7.88 | -7.55 | -4.83 |
| GO:0034330 | GO Biological Processes | cell junction organization | 18 | 6.16 | -7.42 | -4.72 |
| GO:0007160 | GO Biological Processes | cell-matrix adhesion | 16 | 5.48 | -7.34 | -4.66 |
| GO:0098609 | GO Biological Processes | cell-cell adhesion | 31 | 10.62 | -7.19 | -4.52 |
| R-MMU-446353 | Reactome Gene Sets | Cell-extracellular matrix interactions | 6 | 2.05 | -7.04 | -4.38 |
| GO:0051129 | GO Biological Processes | negative regulation of cellular component organization | 31 | 10.62 | -6.76 | -4.12 |
| GO:0000082 | GO Biological Processes | G1/S transition of mitotic cell cycle | 14 | 4.79 | -6.72 | -4.1 |
| GO:0042060 | GO Biological Processes | wound healing | 20 | 6.85 | -6.47 | -3.91 |
| GO:0002064 | GO Biological Processes | epithelial cell development | 16 | 5.48 | -6.18 | -3.66 |
| mmu05200 | KEGG Pathway | Pathways in cancer | 20 | 6.85 | -5.93 | -3.45 |

| | | | | | | |
|---|---|---|---|---|---|---|
| R-MMU-68827 | Reactome Gene Sets | CDT1 association with the CDC6:ORC:origin complex | 8 | 2.74 | -5.77 | -3.29 |
| mmu05205 | KEGG Pathway | Proteoglycans in cancer | 14 | 4.79 | -5.74 | -3.28 |
| GO:0008360 | GO Biological Processes | regulation of cell shape | 12 | 4.11 | -5.74 | -3.28 |

| GO (#5) | Category | Description | Count | % | Log10(P) | Log10(q) |
|---|---|---|---|---|---|---|
| GO:0001944 | GO Biological Processes | vasculature development | 44 | 15.66 | -15.35 | -11.12 |
| GO:0002009 | GO Biological Processes | morphogenesis of an epithelium | 30 | 10.68 | -9.7 | -6.28 |
| GO:0061061 | GO Biological Processes | muscle structure development | 33 | 11.74 | -9.06 | -5.7 |
| R-MMU-1474290 | Reactome Gene Sets | Collagen formation | 11 | 3.91 | -7.83 | -4.56 |
| GO:0071363 | GO Biological Processes | cellular response to growth factor stimulus | 28 | 9.96 | -7.59 | -4.46 |
| GO:0042060 | GO Biological Processes | wound healing | 21 | 7.47 | -7.41 | -4.33 |
| R-MMU-109582 | Reactome Gene Sets | Hemostasis | 25 | 8.9 | -7.32 | -4.33 |
| GO:0040017 | GO Biological Processes | positive regulation of locomotion | 27 | 9.61 | -7.19 | -4.27 |
| GO:0048608 | GO Biological Processes | reproductive structure development | 23 | 8.19 | -7.08 | -4.22 |
| mmu04141 | KEGG Pathway | Protein processing in endoplasmic reticulum | 14 | 4.98 | -7.06 | -4.22 |
| GO:0035270 | GO Biological Processes | endocrine system development | 12 | 4.27 | -6.37 | -3.69 |
| R-MMU-6798695 | Reactome Gene Sets | Neutrophil degranulation | 23 | 8.19 | -5.94 | -3.33 |
| GO:0034976 | GO Biological Processes | response to endoplasmic reticulum stress | 15 | 5.34 | -5.93 | -3.33 |
| GO:0030855 | GO Biological Processes | epithelial cell differentiation | 25 | 8.9 | -5.62 | -3.08 |
| GO:0055123 | GO Biological Processes | digestive system development | 11 | 3.91 | -5.58 | -3.04 |
| GO:0008285 | GO Biological Processes | negative regulation of cell proliferation | 26 | 9.25 | -5.5 | -2.97 |
| GO:0051145 | GO Biological Processes | smooth muscle cell differentiation | 8 | 2.85 | -5.13 | -2.64 |
| GO:0048625 | GO Biological Processes | myoblast fate commitment | 3 | 1.07 | -4.98 | -2.53 |
| GO:0070831 | GO Biological Processes | basement membrane assembly | 4 | 1.42 | -4.96 | -2.52 |
| GO:0007169 | GO Biological Processes | transmembrane receptor protein tyrosine kinase signaling pathway | 22 | 7.83 | -4.94 | -2.52 |

| GO (#6) | Category | Description | Count | % | Log10(P) | Log10(q) |
|---|---|---|---|---|---|---|
| GO:0045216 | GO Biological Processes | cell-cell junction organization | 22 | 7.53 | -15.67 | -11.41 |
| GO:0001890 | GO Biological Processes | placenta development | 22 | 7.53 | -14.31 | -10.35 |
| GO:0030029 | GO Biological Processes | actin filament-based process | 42 | 14.38 | -13.46 | -9.67 |
| mmu04530 | KEGG Pathway | Tight junction | 18 | 6.16 | -10.44 | -7.02 |
| R-MMU-446728 | Reactome Gene Sets | Cell junction organization | 11 | 3.77 | -8.7 | -5.48 |
| GO:0030855 | GO Biological Processes | epithelial cell differentiation | 31 | 10.62 | -8.62 | -5.47 |
| GO:0072659 | GO Biological Processes | protein localization to plasma membrane | 20 | 6.85 | -8.36 | -5.28 |
| mmu04670 | KEGG Pathway | Leukocyte transendothelial migration | 13 | 4.45 | -7.86 | -4.87 |
| GO:0090214 | GO Biological Processes | spongiotrophoblast layer developmental growth | 4 | 1.37 | -7.38 | -4.42 |
| GO:0002934 | GO Biological Processes | desmosome organization | 5 | 1.71 | -6.85 | -3.93 |
| GO:0098609 | GO Biological Processes | cell-cell adhesion | 30 | 10.27 | -6.67 | -3.77 |
| GO:0040017 | GO Biological Processes | positive regulation of locomotion | 26 | 8.9 | -6.3 | -3.5 |
| mmu04810 | KEGG Pathway | Regulation of actin cytoskeleton | 15 | 5.14 | -6.28 | -3.5 |
| R-MMU-4420097 | Reactome Gene Sets | VEGFA-VEGFR2 Pathway | 10 | 3.42 | -6.18 | -3.44 |
| GO:0043086 | GO Biological Processes | negative regulation of catalytic activity | 28 | 9.59 | -6.17 | -3.44 |
| GO:0043588 | GO Biological Processes | skin development | 17 | 5.82 | -5.89 | -3.21 |
| R-MMU-6798695 | Reactome Gene Sets | Neutrophil degranulation | 23 | 7.88 | -5.65 | -3.01 |
| GO:1904019 | GO Biological Processes | epithelial cell apoptotic process | 10 | 3.42 | -5.27 | -2.77 |
| R-MMU-1266738 | Reactome Gene Sets | Developmental Biology | 21 | 7.19 | -4.97 | -2.52 |
| CORUM:3539 | CORUM | Limd1-p62-Traf6-Prkcz complex | 3 | 1.03 | -4.93 | -2.51 |

| GO (#7) | Category | Description | Count | % | Log10(P) | Log10(q) |
|---|---|---|---|---|---|---|
| GO:0051301 | GO Biological Processes | cell division | 50 | 17.01 | -24.1 | -19.84 |
| GO:0030198 | GO Biological Processes | extracellular matrix organization | 32 | 10.88 | -19.32 | -15.85 |
| GO:0001501 | GO Biological Processes | skeletal system development | 38 | 12.93 | -15.41 | -12.37 |
| GO:0001568 | GO Biological Processes | blood vessel development | 42 | 14.29 | -13.92 | -11.02 |
| GO:0001503 | GO Biological Processes | ossification | 30 | 10.2 | -13.26 | -10.47 |
| GO:0050673 | GO Biological Processes | epithelial cell proliferation | 31 | 10.54 | -12.8 | -10.05 |
| R-MMU-1474290 | Reactome Gene Sets | Collagen formation | 15 | 5.1 | -12.21 | -9.48 |
| GO:0070848 | GO Biological Processes | response to growth factor | 36 | 12.24 | -11.89 | -9.2 |
| R-MMU-381426 | Reactome Gene Sets | Regulation of Insulin-like Growth Factor (IGF) transport and uptake by Insulin-like Growth Factor Bi | 15 | 5.1 | -9.88 | -7.32 |
| R-MMU-983189 | Reactome Gene Sets | Kinesins | 11 | 3.74 | -9.76 | -7.23 |
| GO:0051383 | GO Biological Processes | kinetochore organization | 8 | 2.72 | -9.72 | -7.2 |
| GO:0061640 | GO Biological Processes | cytoskeleton-dependent cytokinesis | 13 | 4.42 | -9.06 | -6.61 |
| R-MMU-2129379 | Reactome Gene Sets | Molecules associated with elastic fibres | 9 | 3.06 | -9.03 | -6.58 |
| GO:0001655 | GO Biological Processes | urogenital system development | 23 | 7.82 | -8.82 | -6.38 |
| GO:0007507 | GO Biological Processes | heart development | 31 | 10.54 | -8.53 | -6.14 |
| GO:0048589 | GO Biological Processes | developmental growth | 35 | 11.9 | -8.41 | -6.03 |
| GO:0032963 | GO Biological Processes | collagen metabolic process | 13 | 4.42 | -8.2 | -5.85 |
| GO:0030261 | GO Biological Processes | chromosome condensation | 9 | 3.06 | -7.68 | -5.42 |
| R-MMU-174143 | Reactome Gene Sets | APC/C-mediated degradation of cell cycle proteins | 11 | 3.74 | -7.68 | -5.42 |
| GO:0048144 | GO Biological Processes | fibroblast proliferation | 12 | 4.08 | -7.22 | -4.99 |

| GO (#8) | Category | Description | Count | % | Log10(P) | Log10(q) |
|---|---|---|---|---|---|---|
| GO:0048812 | GO Biological Processes | neuron projection morphogenesis | 68 | 23.13 | -37.1 | -32.83 |
| GO:0120035 | GO Biological Processes | regulation of plasma membrane bounded cell projection organization | 68 | 23.13 | -33.5 | -30.01 |
| GO:0050808 | GO Biological Processes | synapse organization | 46 | 15.65 | -24.95 | -21.77 |
| GO:0097479 | GO Biological Processes | synaptic vesicle localization | 30 | 10.2 | -21.98 | -18.83 |
| GO:0007610 | GO Biological Processes | behavior | 52 | 17.69 | -21.11 | -18 |
| GO:0050804 | GO Biological Processes | modulation of chemical synaptic transmission | 49 | 16.67 | -20.53 | -17.52 |
| R-MMU-112316 | Reactome Gene Sets | Neuronal System | 33 | 11.22 | -18.01 | -15.16 |
| GO:0035418 | GO Biological Processes | protein localization to synapse | 17 | 5.78 | -14.35 | -11.72 |
| GO:0048813 | GO Biological Processes | dendrite morphogenesis | 22 | 7.48 | -13.82 | -11.19 |
| GO:0007611 | GO Biological Processes | learning or memory | 26 | 8.84 | -12.81 | -10.22 |
| GO:0051129 | GO Biological Processes | negative regulation of cellular component organization | 38 | 12.93 | -10.62 | -8.12 |
| GO:1902904 | GO Biological Processes | negative regulation of supramolecular fiber organization | 17 | 5.78 | -10.24 | -7.77 |
| GO:0042391 | GO Biological Processes | regulation of membrane potential | 28 | 9.52 | -10.22 | -7.76 |
| GO:0008088 | GO Biological Processes | axo-dendritic transport | 13 | 4.42 | -10.2 | -7.74 |
| GO:0036465 | GO Biological Processes | synaptic vesicle recycling | 12 | 4.08 | -9.18 | -6.81 |
| GO:0098698 | GO Biological Processes | postsynaptic specialization assembly | 8 | 2.72 | -8.37 | -6.13 |
| GO:0098742 | GO Biological Processes | cell-cell adhesion via plasma-membrane adhesion molecules | 17 | 5.78 | -8.11 | -5.89 |
| mmu04728 | KEGG Pathway | Dopaminergic synapse | 14 | 4.76 | -8.03 | -5.81 |
| GO:0050905 | GO Biological Processes | neuromuscular process | 14 | 4.76 | -7.91 | -5.7 |
| GO:0007420 | GO Biological Processes | brain development | 30 | 10.2 | -7.86 | -5.67 |

| GO (#9) | Category | Description | Count | % | Log10(P) | Log10(q) |
|---|---|---|---|---|---|---|
| GO:0097479 | GO Biological Processes | synaptic vesicle localization | 32 | 11.07 | -24.57 | -20.31 |
| R-MMU-112315 | Reactome Gene Sets | Transmission across Chemical Synapses | 31 | 10.73 | -22.6 | -19.12 |
| GO:0050808 | GO Biological Processes | synapse organization | 42 | 14.53 | -21.52 | -18.21 |
| GO:0050804 | GO Biological Processes | modulation of chemical synaptic transmission | 48 | 16.61 | -20.05 | -16.9 |
| GO:0048812 | GO Biological Processes | neuron projection morphogenesis | 47 | 16.26 | -18.85 | -15.82 |
| GO:0035418 | GO Biological Processes | protein localization to synapse | 18 | 6.23 | -15.77 | -12.95 |
| GO:0120035 | GO Biological Processes | regulation of plasma membrane bounded cell projection organization | 44 | 15.22 | -14.48 | -11.73 |
| GO:0007610 | GO Biological Processes | behavior | 42 | 14.53 | -14 | -11.3 |
| mmu04728 | KEGG Pathway | Dopaminergic synapse | 17 | 5.88 | -11.1 | -8.51 |
| GO:0043269 | GO Biological Processes | regulation of ion transport | 37 | 12.8 | -10.63 | -8.08 |
| GO:0008088 | GO Biological Processes | axo-dendritic transport | 13 | 4.5 | -10.29 | -7.75 |
| GO:0007626 | GO Biological Processes | locomotory behavior | 20 | 6.92 | -9.15 | -6.66 |
| GO:0007215 | GO Biological Processes | glutamate receptor signaling pathway | 13 | 4.5 | -9.1 | -6.62 |
| GO:0072657 | GO Biological Processes | protein localization to membrane | 29 | 10.03 | -8.94 | -6.5 |
| GO:0036465 | GO Biological Processes | synaptic vesicle recycling | 11 | 3.81 | -8.12 | -5.72 |

| | | | | | | |
|---|---|---|---|---|---|---|
| GO:0048168 | GO Biological Processes | regulation of neuronal synaptic plasticity | 10 | 3.46 | -7.45 | -5.09 |
| R-MMU-111885 | Reactome Gene Sets | Opioid Signalling | 10 | 3.46 | -7.33 | -4.99 |
| GO:0007416 | GO Biological Processes | synapse assembly | 15 | 5.19 | -7.32 | -4.99 |
| GO:0032984 | GO Biological Processes | protein-containing complex disassembly | 16 | 5.54 | -7.08 | -4.78 |
| GO:0051952 | GO Biological Processes | regulation of amine transport | 12 | 4.15 | -6.92 | -4.64 |

| GO (#10) | Category | Description | Count | % | Log10(P) | Log10(q) |
|---|---|---|---|---|---|---|
| GO:0045664 | GO Biological Processes | regulation of neuron differentiation | 53 | 18.09 | -20.71 | -16.45 |
| R-MMU-112315 | Reactome Gene Sets | Transmission across Chemical Synapses | 28 | 9.56 | -19.05 | -15.52 |
| GO:0097479 | GO Biological Processes | synaptic vesicle localization | 27 | 9.22 | -18.62 | -15.2 |
| GO:0050804 | GO Biological Processes | modulation of chemical synaptic transmission | 42 | 14.33 | -15.21 | -12.19 |
| GO:0007610 | GO Biological Processes | behavior | 42 | 14.33 | -13.79 | -10.9 |
| GO:0035418 | GO Biological Processes | protein localization to synapse | 16 | 5.46 | -13.13 | -10.31 |
| GO:0050808 | GO Biological Processes | synapse organization | 32 | 10.92 | -12.89 | -10.11 |
| GO:0031345 | GO Biological Processes | negative regulation of cell projection organization | 22 | 7.51 | -12.57 | -9.85 |
| GO:0042391 | GO Biological Processes | regulation of membrane potential | 29 | 9.9 | -10.98 | -8.37 |
| GO:0043269 | GO Biological Processes | regulation of ion transport | 37 | 12.63 | -10.46 | -7.9 |
| GO:0036465 | GO Biological Processes | synaptic vesicle recycling | 12 | 4.1 | -9.2 | -6.75 |
| mmu04728 | KEGG Pathway | Dopaminergic synapse | 15 | 5.12 | -9 | -6.57 |
| GO:0097091 | GO Biological Processes | synaptic vesicle clustering | 7 | 2.39 | -8.48 | -6.08 |
| GO:0007215 | GO Biological Processes | glutamate receptor signaling pathway | 12 | 4.1 | -7.97 | -5.59 |
| GO:0007420 | GO Biological Processes | brain development | 30 | 10.24 | -7.9 | -5.53 |
| GO:1990778 | GO Biological Processes | protein localization to cell periphery | 21 | 7.17 | -7.5 | -5.15 |
| mmu04727 | KEGG Pathway | GABAergic synapse | 11 | 3.75 | -7.26 | -4.93 |
| GO:0031110 | GO Biological Processes | regulation of microtubule polymerization or depolymerization | 11 | 3.75 | -7.26 | -4.93 |
| GO:0016358 | GO Biological Processes | dendrite development | 19 | 6.48 | -7.22 | -4.91 |
| R-MMU-5576892 | Reactome Gene Sets | Phase 0 - rapid depolarisation | 8 | 2.73 | -6.88 | -4.63 |

| GO (#11) | Category | Description | Count | % | Log10(P) | Log10(q) |
|---|---|---|---|---|---|---|
| R-MMU-1640170 | Reactome Gene Sets | Cell Cycle | 70 | 23.73 | -45.01 | -40.74 |
| GO:0006260 | GO Biological Processes | DNA replication | 28 | 9.49 | -16.62 | -13.74 |
| R-MMU-174143 | Reactome Gene Sets | APC/C-mediated degradation of cell cycle proteins | 17 | 5.76 | -14.8 | -11.97 |
| GO:0000226 | GO Biological Processes | microtubule cytoskeleton organization | 37 | 12.54 | -13.65 | -10.86 |
| GO:0010564 | GO Biological Processes | regulation of cell cycle process | 37 | 12.54 | -13.28 | -10.51 |
| GO:0051383 | GO Biological Processes | kinetochore organization | 9 | 3.05 | -11.43 | -8.71 |
| R-MMU-8953854 | Reactome Gene Sets | Metabolism of RNA | 30 | 10.17 | -10.15 | -7.47 |
| GO:0061640 | GO Biological Processes | cytoskeleton-dependent cytokinesis | 14 | 4.75 | -10.14 | -7.47 |
| mmu04110 | KEGG Pathway | Cell cycle | 15 | 5.08 | -9.44 | -6.87 |
| R-MMU-69275 | Reactome Gene Sets | G2/M Transition | 15 | 5.08 | -8.96 | -6.42 |
| GO:0006974 | GO Biological Processes | cellular response to DNA damage stimulus | 33 | 11.19 | -7.94 | -5.51 |
| GO:0140013 | GO Biological Processes | meiotic nuclear division | 15 | 5.08 | -7.24 | -4.89 |
| R-MMU-176408 | Reactome Gene Sets | Regulation of APC/C activators between G1/S and early anaphase | 7 | 2.37 | -7.06 | -4.72 |
| R-MMU-72662 | Reactome Gene Sets | Activation of the mRNA upon binding of the cap-binding complex and eIFs, and subsequent bindir | 9 | 3.05 | -6.77 | -4.48 |
| GO:0006323 | GO Biological Processes | DNA packaging | 14 | 4.75 | -6.73 | -4.44 |
| GO:0008608 | GO Biological Processes | attachment of spindle microtubules to kinetochore | 7 | 2.37 | -6.71 | -4.44 |
| R-MMU-983189 | Reactome Gene Sets | Kinesins | 8 | 2.71 | -6.11 | -3.89 |
| R-MMU-73894 | Reactome Gene Sets | DNA Repair | 18 | 6.1 | -6.06 | -3.85 |
| GO:0009263 | GO Biological Processes | deoxyribonucleotide biosynthetic process | 5 | 1.69 | -5.95 | -3.74 |
| R-MMU-3700989 | Reactome Gene Sets | Transcriptional Regulation by TP53 | 15 | 5.08 | -5.91 | -3.71 |

| GO (#12) | Category | Description | Count | % | Log10(P) | Log10(q) |
|---|---|---|---|---|---|---|
| mmu04141 | KEGG Pathway | Protein processing in endoplasmic reticulum | 23 | 7.99 | -15.67 | -11.41 |
| GO:0002009 | GO Biological Processes | morphogenesis of an epithelium | 37 | 12.85 | -14.34 | -10.38 |
| GO:0001944 | GO Biological Processes | vasculature development | 42 | 14.58 | -13.57 | -9.79 |
| GO:0043062 | GO Biological Processes | extracellular structure organization | 23 | 7.99 | -9.81 | -6.55 |
| GO:0001701 | GO Biological Processes | in utero embryonic development | 29 | 10.07 | -9.07 | -6.04 |
| GO:0071363 | GO Biological Processes | cellular response to growth factor stimulus | 29 | 10.07 | -7.94 | -5.08 |
| GO:0034976 | GO Biological Processes | response to endoplasmic reticulum stress | 16 | 5.56 | -6.53 | -3.78 |
| GO:0009100 | GO Biological Processes | glycoprotein metabolic process | 19 | 6.6 | -6.4 | -3.68 |
| GO:0035270 | GO Biological Processes | endocrine system development | 12 | 4.17 | -6.26 | -3.57 |
| GO:0035886 | GO Biological Processes | vascular smooth muscle cell differentiation | 7 | 2.43 | -6.2 | -3.54 |
| R-MMU-6811434 | Reactome Gene Sets | COPI-dependent Golgi-to-ER retrograde traffic | 10 | 3.47 | -6.14 | -3.49 |
| GO:0009611 | GO Biological Processes | response to wounding | 23 | 7.99 | -6.1 | -3.46 |
| GO:0061061 | GO Biological Processes | muscle structure development | 28 | 9.72 | -6.06 | -3.43 |
| GO:0018126 | GO Biological Processes | protein hydroxylation | 6 | 2.08 | -5.96 | -3.37 |
| GO:0040017 | GO Biological Processes | positive regulation of locomotion | 25 | 8.68 | -5.88 | -3.33 |
| GO:2000027 | GO Biological Processes | regulation of animal organ morphogenesis | 14 | 4.86 | -5.72 | -3.2 |
| CORUM:414 | CORUM | (ER)-localized multiprotein complex, in absence of Ig heavy chains | 4 | 1.39 | -5.33 | -2.87 |
| GO:0007492 | GO Biological Processes | endoderm development | 8 | 2.78 | -5.23 | -2.8 |
| GO:0048806 | GO Biological Processes | genitalia development | 7 | 2.43 | -5.14 | -2.72 |
| mmu04918 | KEGG Pathway | Thyroid hormone synthesis | 8 | 2.78 | -5.05 | -2.65 |

| GO (#13) | Category | Description | Count | % | Log10(P) | Log10(q) |
|---|---|---|---|---|---|---|
| R-MMU-8953854 | Reactome Gene Sets | Metabolism of RNA | 51 | 18.15 | -28.58 | -24.46 |
| R-MMU-72613 | Reactome Gene Sets | Eukaryotic Translation Initiation | 23 | 8.19 | -19.42 | -15.86 |
| CORUM:3047 | CORUM | Parvulin-associated pre-rRNP complex | 13 | 4.63 | -13.06 | -10.18 |
| GO:0016071 | GO Biological Processes | mRNA metabolic process | 37 | 13.17 | -12.79 | -9.93 |
| GO:0042273 | GO Biological Processes | ribosomal large subunit biogenesis | 14 | 4.98 | -11.27 | -8.47 |
| GO:0072594 | GO Biological Processes | establishment of protein localization to organelle | 23 | 8.19 | -8.57 | -5.93 |
| GO:0097193 | GO Biological Processes | intrinsic apoptotic signaling pathway | 20 | 7.12 | -8.28 | -5.66 |
| GO:0002181 | GO Biological Processes | cytoplasmic translation | 11 | 3.91 | -7.14 | -4.59 |
| GO:0042274 | GO Biological Processes | ribosomal small subunit biogenesis | 9 | 3.2 | -6.52 | -4 |
| GO:0072331 | GO Biological Processes | signal transduction by p53 class mediator | 12 | 4.27 | -6.37 | -3.87 |
| GO:1903829 | GO Biological Processes | positive regulation of cellular protein localization | 17 | 6.05 | -6.02 | -3.57 |
| GO:0031647 | GO Biological Processes | regulation of protein stability | 16 | 5.69 | -5.79 | -3.35 |
| GO:0006986 | GO Biological Processes | response to unfolded protein | 10 | 3.56 | -5.52 | -3.09 |
| GO:0042594 | GO Biological Processes | response to starvation | 12 | 4.27 | -5.42 | -3 |
| GO:0045727 | GO Biological Processes | positive regulation of translation | 10 | 3.56 | -5.18 | -2.79 |
| mmu04142 | KEGG Pathway | Lysosome | 10 | 3.56 | -5.05 | -2.68 |
| GO:0000470 | GO Biological Processes | maturation of LSU-rRNA | 5 | 1.78 | -4.51 | -2.22 |
| mmu04115 | KEGG Pathway | p53 signaling pathway | 7 | 2.49 | -4.37 | -2.1 |
| GO:0042256 | GO Biological Processes | mature ribosome assembly | 3 | 1.07 | -4.29 | -2.04 |
| R-MMU-4570464 | Reactome Gene Sets | SUMOylation of RNA binding proteins | 3 | 1.07 | -4.29 | -2.04 |

| GO (#14) | Category | Description | Count | % | Log10(P) | Log10(q) |
|---|---|---|---|---|---|---|
| GO:0050673 | GO Biological Processes | epithelial cell proliferation | 35 | 11.82 | -15.94 | -11.68 |
| GO:0097435 | GO Biological Processes | supramolecular fiber organization | 41 | 13.85 | -13.88 | -10.13 |
| GO:2000147 | GO Biological Processes | positive regulation of cell motility | 37 | 12.5 | -13.79 | -10.13 |
| GO:0030324 | GO Biological Processes | lung development | 21 | 7.09 | -10.92 | -7.78 |
| R-MMU-381426 | Reactome Gene Sets | Regulation of Insulin-like Growth Factor (IGF) transport and uptake by Insulin-like Growth Factor Bi | 16 | 5.41 | -10.89 | -7.78 |
| GO:0045785 | GO Biological Processes | positive regulation of cell adhesion | 28 | 9.46 | -10.82 | -7.75 |
| R-MMU-1474244 | Reactome Gene Sets | Extracellular matrix organization | 22 | 7.43 | -10.1 | -7.09 |
| mmu04510 | KEGG Pathway | Focal adhesion | 19 | 6.42 | -9.96 | -6.98 |
| GO:0042060 | GO Biological Processes | wound healing | 25 | 8.45 | -9.83 | -6.89 |
| GO:0060485 | GO Biological Processes | mesenchyme development | 21 | 7.09 | -9.23 | -6.38 |
| GO:0048729 | GO Biological Processes | tissue morphogenesis | 33 | 11.15 | -8.83 | -6.01 |
| R-MMU-72689 | Reactome Gene Sets | Formation of a pool of free 40S subunits | 13 | 4.39 | -8.63 | -5.85 |
| GO:0010942 | GO Biological Processes | positive regulation of cell death | 33 | 11.15 | -8.57 | -5.8 |

| | | | | | | |
|---|---|---|---|---|---|---|
| GO:0030855 | GO Biological Processes | epithelial cell differentiation | 31 | 10.47 | -8.47 | -5.73 |
| mmu04115 | KEGG Pathway | p53 signaling pathway | 11 | 3.72 | -8.42 | -5.69 |
| GO:0042063 | GO Biological Processes | gliogenesis | 21 | 7.09 | -7.9 | -5.29 |
| GO:0030111 | GO Biological Processes | regulation of Wnt signaling pathway | 20 | 6.76 | -7.82 | -5.24 |
| GO:0001501 | GO Biological Processes | skeletal system development | 27 | 9.12 | -7.7 | -5.15 |
| GO:0045596 | GO Biological Processes | negative regulation of cell differentiation | 34 | 11.49 | -7.64 | -5.1 |
| GO:0070848 | GO Biological Processes | response to growth factor | 29 | 9.8 | -7.46 | -4.97 |

| GO (#15) | Category | Description | Count | % | Log10(P) | Log10(q) |
|---|---|---|---|---|---|---|
| GO:0045664 | GO Biological Processes | regulation of neuron differentiation | 47 | 16.61 | -16.87 | -12.61 |
| GO:0048812 | GO Biological Processes | neuron projection morphogenesis | 39 | 13.78 | -13.28 | -9.32 |
| GO:0007389 | GO Biological Processes | pattern specification process | 31 | 10.95 | -12.17 | -8.75 |
| GO:0021537 | GO Biological Processes | telencephalon development | 23 | 8.13 | -12.06 | -8.7 |
| GO:0050768 | GO Biological Processes | negative regulation of neurogenesis | 24 | 8.48 | -10.15 | -7.12 |
| GO:0006208 | GO Biological Processes | pyrimidine nucleobase catabolic process | 5 | 1.77 | -7.56 | -4.79 |
| GO:0021953 | GO Biological Processes | central nervous system neuron differentiation | 16 | 5.65 | -6.94 | -4.23 |
| GO:0051098 | GO Biological Processes | regulation of binding | 21 | 7.42 | -6.6 | -3.95 |
| GO:0001764 | GO Biological Processes | neuron migration | 14 | 4.95 | -6.47 | -3.84 |
| GO:0003357 | GO Biological Processes | noradrenergic neuron differentiation | 4 | 1.41 | -6.27 | -3.66 |
| GO:0006338 | GO Biological Processes | chromatin remodeling | 11 | 3.89 | -5.33 | -2.79 |
| GO:0060412 | GO Biological Processes | ventricular septum morphogenesis | 7 | 2.47 | -5.19 | -2.67 |
| GO:0048732 | GO Biological Processes | gland development | 20 | 7.07 | -5.01 | -2.52 |
| GO:0021516 | GO Biological Processes | dorsal spinal cord development | 5 | 1.77 | -4.96 | -2.49 |
| CORUM:572 | CORUM | PYR complex | 4 | 1.41 | -4.78 | -2.37 |
| R-MMU-156827 | Reactome Gene Sets | L13a-mediated translational silencing of Ceruloplasmin expression | 9 | 3.18 | -4.63 | -2.23 |
| GO:0007623 | GO Biological Processes | circadian rhythm | 12 | 4.24 | -4.43 | -2.07 |
| GO:0021545 | GO Biological Processes | cranial nerve development | 6 | 2.12 | -4.23 | -1.91 |
| GO:0051101 | GO Biological Processes | regulation of DNA binding | 9 | 3.18 | -4.08 | -1.78 |
| GO:0010608 | GO Biological Processes | posttranscriptional regulation of gene expression | 19 | 6.71 | -4.03 | -1.74 |

| GO (#16) | Category | Description | Count | % | Log10(P) | Log10(q) |
|---|---|---|---|---|---|---|
| R-MMU-1799339 | Reactome Gene Sets | SRP-dependent cotranslational protein targeting to membrane | 55 | 22.54 | -83.15 | -78.89 |
| CORUM:3047 | CORUM | Parvulin-associated pre-rRNP complex | 18 | 7.38 | -21.93 | -19.06 |
| GO:0022613 | GO Biological Processes | ribonucleoprotein complex biogenesis | 36 | 14.75 | -19.16 | -16.31 |
| GO:0042273 | GO Biological Processes | ribosomal large subunit biogenesis | 16 | 6.56 | -14.72 | -11.9 |
| GO:0042274 | GO Biological Processes | ribosomal small subunit biogenesis | 12 | 4.92 | -10.72 | -7.98 |
| GO:0051301 | GO Biological Processes | cell division | 25 | 10.25 | -7.52 | -4.82 |
| R-MMU-176417 | Reactome Gene Sets | Phosphorylation of Emi1 | 4 | 1.64 | -7 | -4.32 |
| GO:1904666 | GO Biological Processes | regulation of ubiquitin protein ligase activity | 6 | 2.46 | -6.88 | -4.21 |
| GO:0051052 | GO Biological Processes | regulation of DNA metabolic process | 20 | 8.2 | -6.4 | -3.77 |
| R-MMU-176408 | Reactome Gene Sets | Regulation of APC/C activators between G1/S and early anaphase | 6 | 2.46 | -6.17 | -3.56 |
| CORUM:161 | CORUM | SWAP complex | 3 | 1.23 | -5.17 | -2.65 |
| GO:0000470 | GO Biological Processes | maturation of LSU-rRNA | 5 | 2.05 | -4.81 | -2.32 |
| R-MMU-8852276 | Reactome Gene Sets | The role of GTSE1 in G2/M progression after G2 checkpoint | 5 | 2.05 | -4.57 | -2.12 |
| GO:0051321 | GO Biological Processes | meiotic cell cycle | 14 | 5.74 | -4.54 | -2.1 |
| GO:0032526 | GO Biological Processes | response to retinoic acid | 7 | 2.87 | -3.98 | -1.61 |
| GO:0006119 | GO Biological Processes | oxidative phosphorylation | 7 | 2.87 | -3.77 | -1.43 |
| GO:0006304 | GO Biological Processes | DNA modification | 7 | 2.87 | -3.77 | -1.43 |
| GO:0001731 | GO Biological Processes | formation of translation preinitiation complex | 3 | 1.23 | -3.46 | -1.17 |
| GO:0071353 | GO Biological Processes | cellular response to interleukin-4 | 4 | 1.64 | -3.42 | -1.14 |
| R-MMU-8866652 | Reactome Gene Sets | Synthesis of active ubiquitin: roles of E1 and E2 enzymes | 4 | 1.64 | -3.36 | -1.1 |
