## Extended Data Table 9 for "Autism in a dish: ES cell models of autism with copy number variations reveal cell-type-specific vulnerability"

**Extended Data Table 9. Gene Ontology analysis; ASD associated 12 CNVs**

| GO | Category | Description | Count | % | Log10(P) | Log10(q) |
|---|---|---|---|---|---|---|
| R-MMU-72706 | Reactome Gene Sets | GTP hydrolysis and joining of the 60S ribosomal subunit | 89 | 8.43 | -93.36 | -89.1 |
| GO:0022613 | GO Biological Processes | ribonucleoprotein complex biogenesis | 93 | 8.81 | -31.35 | -28.46 |
| CORUM:3047 | CORUM | Parvulin-associated pre-rRNP complex | 30 | 2.84 | -25.84 | -22.98 |
| GO:0022618 | GO Biological Processes | ribonucleoprotein complex assembly | 59 | 5.59 | -25.26 | -22.41 |
| mmu05016 | KEGG Pathway | Huntington's disease | 55 | 5.21 | -25.2 | -22.37 |
| GO:0034248 | GO Biological Processes | regulation of cellular amide metabolic process | 70 | 6.63 | -19.77 | -17.04 |
| R-MMU-1640170 | Reactome Gene Sets | Cell Cycle | 86 | 8.14 | -19.5 | -16.79 |
| GO:0008380 | GO Biological Processes | RNA splicing | 67 | 6.34 | -19.04 | -16.34 |
| GO:0009167 | GO Biological Processes | purine ribonucleoside monophosphate metabolic process | 55 | 5.21 | -17.68 | -15.05 |
| R-MMU-8953897 | Reactome Gene Sets | Cellular responses to external stimuli | 69 | 6.53 | -17.3 | -14.73 |
| GO:0120035 | GO Biological Processes | regulation of plasma membrane bounded cell projection organization | 58 | 13.65 | -16.65 | -14.04 |
| GO:0006413 | GO Biological Processes | translational initiation | 31 | 2.94 | -15.14 | -12.66 |
| GO:0006457 | GO Biological Processes | protein folding | 37 | 3.5 | -14.68 | -12.21 |
| R-MMU-422475 | Reactome Gene Sets | Axon guidance | 48 | 4.55 | -14.29 | -11.84 |
| R-MMU-8852276 | Reactome Gene Sets | The role of GTSE1 in G2/M progression after G2 checkpoint | 13 | 3.16 | -14.15 | -11.68 |
| GO:0032990 | GO Biological Processes | cell part morphogenesis | 36 | 16.07 | -13.61 | -11.18 |
| GO:1903827 | GO Biological Processes | regulation of cellular protein localization | 72 | 6.82 | -13.56 | -11.12 |
| GO:0007005 | GO Biological Processes | mitochondrion organization | 65 | 6.16 | -12.78 | -10.37 |
| R-MMU-1280218 | Reactome Gene Sets | Adaptive Immune System | 77 | 7.29 | -12.41 | -10.03 |
| GO:0033365 | GO Biological Processes | protein localization to organelle | 90 | 8.52 | -12.3 | -9.94 |
